# gcSV: a unified framework for comprehensive structural variant detection

**DOI:** 10.1101/2025.02.10.637589

**Authors:** Gaoyang Li, Yadong Liu, Bo Liu, Long Qian, Yadong Wang

## Abstract

The characterization of structural variants (SVs) is fundamental to genomic studies and advanced computational approaches are on demand to exert the ability of the ubiquitous high-throughput sequencing data. Herein, we propose gcSV, a read-length agnostic alignment-based approach to well-handle the issues of genome repeats, SV breakpoints and read alignments/assemblies for comprehensive, cost-effective and versatile SV calling. For long reads, its yield is 20-38% higher than state-of-the-art tools in HG002 benchmark. For hybrid sequencing, it provides a cost-effective solution (2-4x long plus 30-60x short reads) to achieve even higher yield than that of state-of-the-art tools using 30x long reads. For short reads, gcSV also achieves over 8% higher precision without any loss of sensitivity. Furthermore, gcSV confidently brings over 93,000 novel SVs comparing to the official callset of 1000 Genomes Project Phase4. The results suggest that gcSV is promising to make valuable SV discoveries in many cutting-edge studies.

## Introduction

Structural variants (SVs) are genomic sequence alternations over 50 bp which account for the largest divergences among human genomes and exhibit population stratification^1, 2^. SVs are involved in distinct genome-wide mutational processes^3, 4^, leading to significant variations in both genetic and epigenetic landscapes^5, 6^. Moreover, SVs are strongly associated with various phenotypes and diseases, particularly serving as the primary drivers of oncogenesis and genetic disorders^7-9^. Due to its important roles, the characterization of SV is fundamental to many genomic studies. With the rapid development of high-throughput sequencing (HTS) technologies, it is on wide demand for tailored approaches to comprehensively discover SV events from HTS data.

State-of-the-art HTS-based SV calling approaches roughly fall in two categories, i.e., alignment-based and assembly-based. Alignment-based approaches^10-20^ detects SVs by the non-colinear alignment of SV-spanning reads to the reference, which manifests in forms of split alignments, discordant read-pairs, large clippings and gain or loss of local read depths^21-23^, depending on read length, quality and mappability. Subsequently, the types and locations of SVs are determined by either clustering reads associated with the focal SV and applying statistical inference^15-17^, or local read assembly and realignment^10, 19^. As many tools are tailored for various read lengths, depths and qualities, alignment-based approach is flexible to data configurations and hence ubiquitously used in large-scale genomics studies^24-26^. Moreover, efforts have also been made to handle the SVs of specific types and/or in specific genomic regions^27-29^. Furthermore, in many studies, pipelines are built to integrate multiple tools to improve overall yields^30, 31^. Regardless of these efforts, alignment-based approaches are heavily influenced by the complexity of alignments, especially fall short of resolving SVs in highly repetitive genomic regions and/or in complex forms, since the alignment signatures of SVs are highly ambiguous and heterogeneous in both cases.

On the other hand, assembly-based SV calling^32, 33^ initially implements whole genome assembly and then discovers SVs from the alignments between the assembled contigs and the reference. Given that long read-based assemblers are capable of producing chromosome level contigs^34-37^, assembly-based approaches leverage the high mappability of contigs for the detection of very large and/or complex SVs while avoiding potential reference bias. However, in practice, to handle repeats and extend the contig length, such approaches heavily rely on high-coverage long-read sequencing data, sometimes requiring multiple types of sequencing, as in many telomere-to-telomere (T2T) assembly studies^38-40^. This premise incurs prohibitive costs especially for large-scale studies. Moreover, state-of-the-art assemblers still struggle with mis-assemblies^41-43^ and haplotype-level assembly^44^, which inevitably lead to false positives or affect SV genotyping.

Herein, we present Genome Context-driven Structural Variation Caller (gcSV), a read-length agnostic alignment-based SV calling approach that enables comprehensive and cost-effective SV discovery. gcSV fully considers the various issues in reference genome and HTS data that affects the analysis of alignment-based SV signatures, such as genome repeats, SV breakpoints, non-colinear read alignment and local assembly, to break the bottlenecks of comprehensive loci recognition, precise signature clustering and repeat-robust allele reconstruction, thereby achieving substantially higher yields of SV calling than state-of-the-art tools with equal or lower coverage data. Furthermore, gcSV implements a unified framework to natively integrate short read sequencing (SRS) and long read sequencing (LRS) reads, enabling the use of hybrid sequencing data for cost-effective SV calling. Applying gcSV on commonly used coverage (i.e., 30x) of LRS data, we observed notable improvements in the sensitivity of SV calling without compromising the precision. When applied in conjunction with SRS data, gcSV reduced the requirement for LRS data by over 80% while still achieving yields of major SV callers such as Sniffles ^15^ and cuteSV^17^. For SRS only scenario, gcSV outperformed state-of-the-art pipelines consisting of multiple tools and pangenomes such as GATK-SV^1^ and DRAGEN^30^, meanwhile showcasing distinctive abilities to detect SVs which were thought intractable based on short reads. Besides the potential to enhance SV detection at reduced sequencing cost for ongoing and upcoming population-scale genomics studies^45, 46^, we demonstrated the power of gcSV in re-analyzing existing short-read genome sequencing data to unveil previously hard-to-identify SVs.

## Results

### Overview of gcSV

gcSV is an alignment-based approach that allows LRS, SRS and hybrid sequencing data as inputs. In gcSV, the three major steps of SV calling—SV site recognition, SV-spanning read clustering, SV calling and genotyping—are devised with principles tailored to the contexts of reference genome structure, SV events and data signatures, respectively, to achieve sensitive and precise calls. gcSV’s key design principles are outlined as following (Fig. 1, Online Methods).

**Fig 1.**
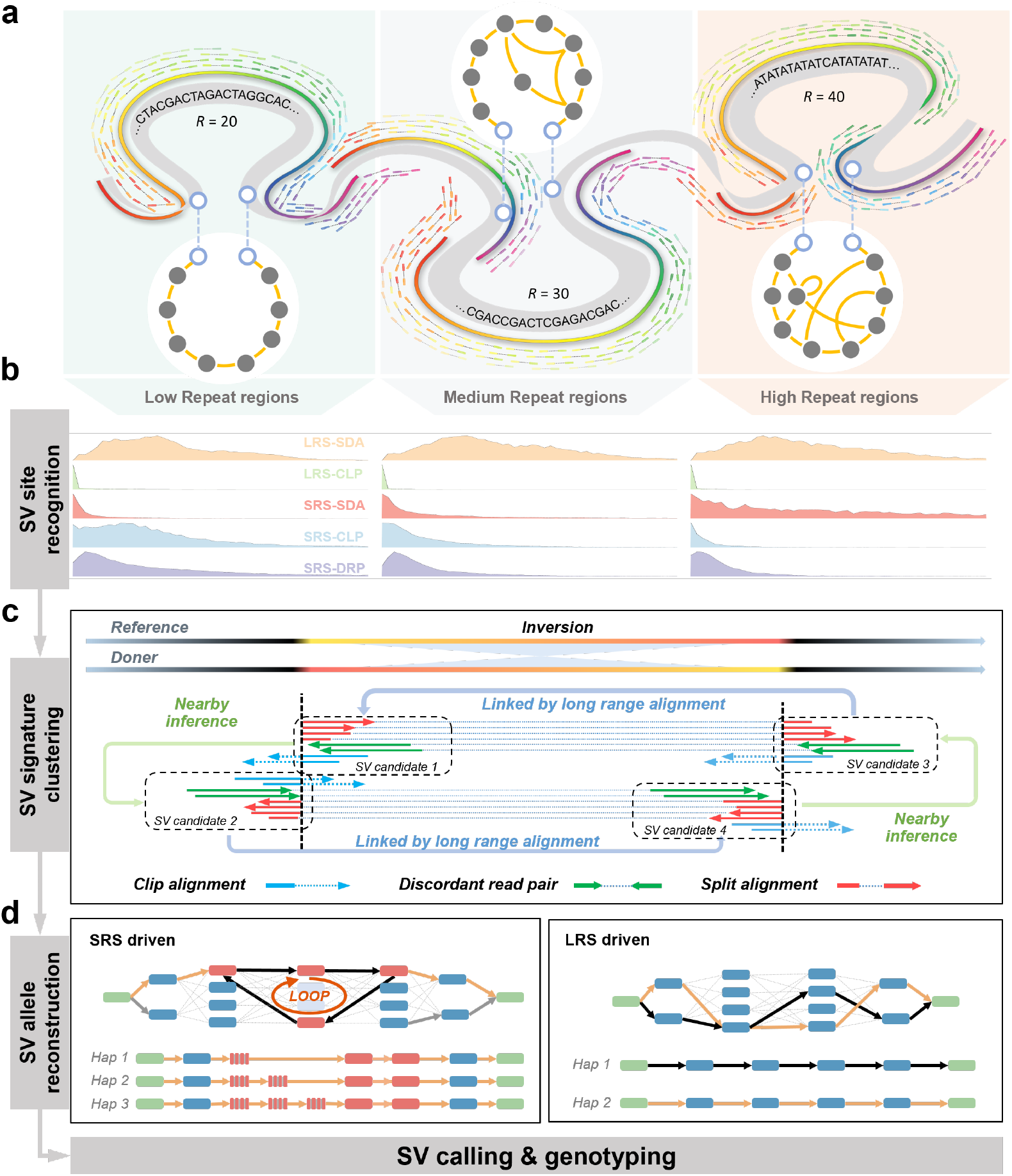
A schematic illustration of gcSV. **a**, Schematic of gcSV, an alignment-based SV calling approach for both LRS and SRS data. **b**, Repeat context-driven SV site recognition. gcSV learns the distributions of all types of alignment signals, including sequence divergences in alignments (SDA, for LRS and SRS), large clippings (CLP, for LRS and SRS) and discordant read pairs (DRP, only for SR), within SV spanning reads under various degrees of local repetitiveness. **c**, Fine-grained SV signature clustering. gcSV constructs probabilistic models to infer virtual SV breakpoints using alignments signals of the clustered reads, and then connects the clusters around various breakpoints in contexts of genomic positions or long-range information of read pairs or split alignments. Inversion is shown as an example. **d**, Heuristic walking-based SV reconstruction. gcSV generates candidate SV alleles/contigs by either coverage heuristics-based (solely using k-mers, left panel) walks or long read-guided (if available, right panel) walks on a local de Bruijn graph, and then uses a k-mer frequency-based approach to test the generated contigs to infer the most likely SV-allele(s) and implement genotyping.

Step 1: Repeat context-driven SV site recognition. Due to the omnipresence of repetitive sequences in human genome, in these regions, read alignment signals indicative of SV-alleles, such as split alignments or discordant read-pairs, may be largely attenuated or even vanish from mistakenly colinear alignments (Supplementary Fig. 1). Unlike most SV callers that disregard the complexity of read alignments under various genomic repeat contexts, gcSV proposes a repeat context-driven approach to learn the distributions of all types of signals within SV spanning reads under various degrees of local repetitiveness. These distributions effectively capture the divergence of signals at various degrees of repetitiveness, unveiling distinctive SV signatures in highly repetitive regions such as dense CIGARs. Further, gcSV adopts a statistical model based on the distributions to well-handle the attenuation and distortion of SV signatures and implement comprehensive SV loci recognition.

Step 2: Fine-grained SV signature clustering. Due to the complexity of underlying SV events, reads spanning an SV locus may carry confounding signals. In particular, to infer SV events from read signals, current practices of local read clustering^12, 13, 15, 17^ often fall short to identify long-range parity of signals for large SVs, to resolve mixed signals in heterozygous SV sites, and to address ambiguous alignment in repetitive alleles. gcSV addresses these issues by a bottom-up approach. Firstly, with an iterative algorithm, gcSV adaptively composes the probability distribution of the latent SV breakpoint by the aligned reads and discards the unlikely reads, thereby maximizing the overall read clustering purity (Supplementary Fig. 2a). Secondly, gcSV uses a greedy integration approach to further combine adjacent or linked clusters having compatible signals to pool reads of the same SV event together (Supplementary Fig. 2b). By this approach, gcSV not only aggregates interspersed reads to join distanced SV breakpoints of large deletions, inversions, translocations and/or genomic repeats (e.g., mobile element insertions), but also disentangles divergent SV signatures for precise SV reconstruction.

Step 3: Heuristic walking-based SV reconstruction. Even with comprehensively collected spanning reads, highly repetitive SV alleles are hard to reconstruct precisely. To overcome this difficulty, gcSV implements a constrained exhaustive search by either coverage heuristics-based (solely using k-mers) or long read-guided (if available) walks along a local de Bruijn graph (Supplementary Fig. 3). The heuristic walking broadly generates a set of plausible contigs irrespective of read-length limitations imposed by SRS data. Further, gcSV incorporates all information of read (both short and long) realignment to the candidate contigs, and uses a statistical model on k-mer frequencies to select the most likely contigs for making SV calls and genotyping. Such an approach solves the dilemma of commonly used local assembly strategies^10, 19^ that produce either conservative but shortened SV alleles or aggressive but imprecise SV alleles.

### Benchmarks suggest the high yields and versatility of gcSV

We used a well-studied human sample (GIAB HG002) to assess the SV calling ability of gcSV in various sequencing scenarios (short read-, long read- and hybrid sequencing). One long-read (PacBio HiFi, 30x coverage) and two short-read (Illumina, 60x and 35x coverages, respectively) real sequencing datasets (Supplementary Table 1) were used to compose various data configurations by down-sampling and/or mixing. NIST HG002 Q100 SV benchmark V1.0^47, 48^ (NIST-Q100) was employed as ground truth and Truvari^49^ was used as the assessment tool (by its “single” mode, i.e., allowing each variant to participate in up to one match, unless specified otherwise). In all scenarios, gcSV achieved the highest yields compared to state-of-the-art tools, demonstrating its outstanding ability and versatility, as well as the potential of cost-effective SV calling (Fig. 2 and Supplementary Tables 2-4).

**Fig 2.**
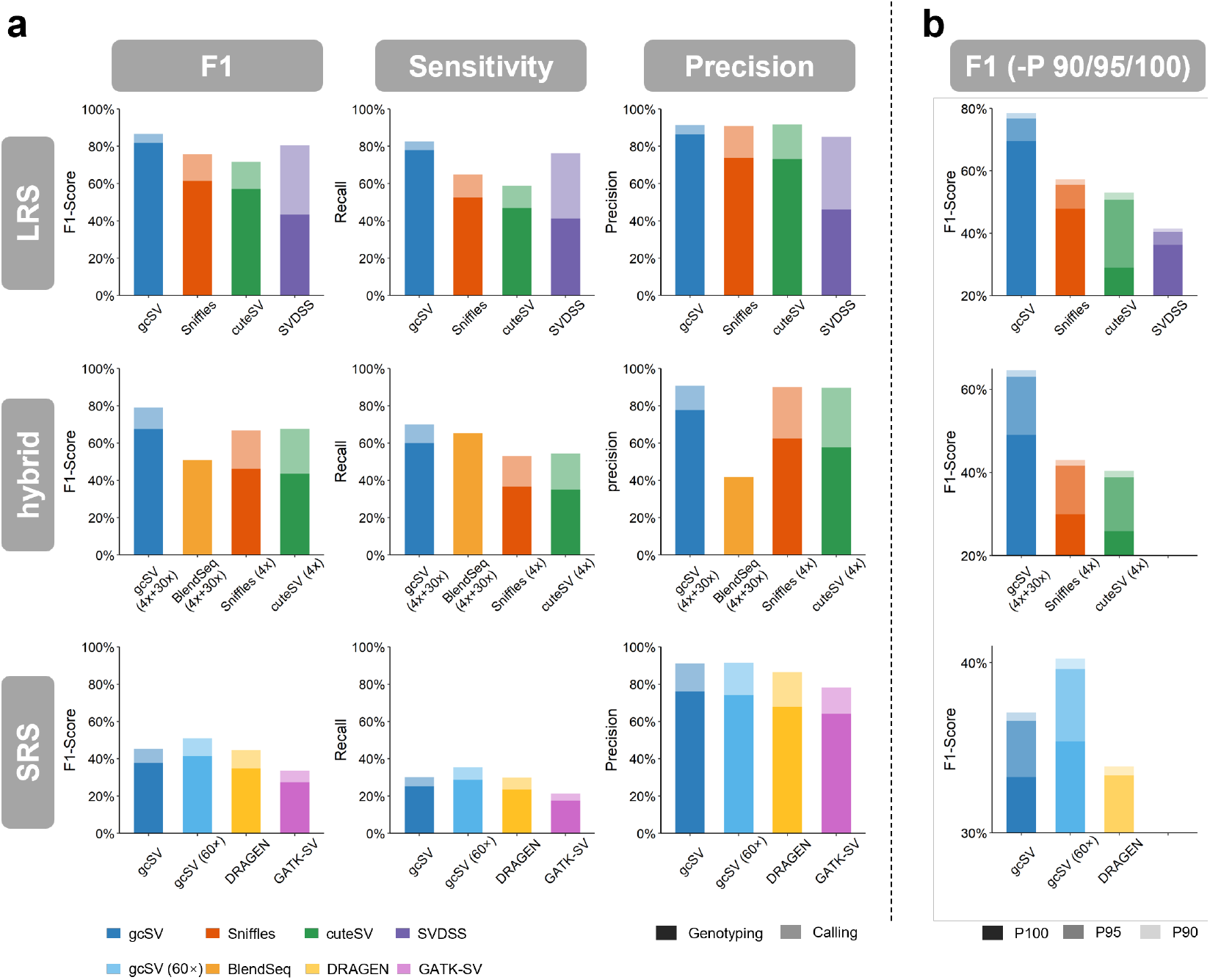
Overall performance evaluation of gcSV applied to various data configurations. **a**, The three rows indicate the benchmark evaluations of various SV callers for long read, hybrid and short read sequencing data, respectively. The three columns correspond to F1-scores, sensitivities and precisions of the callers, respectively. The dark- and light-colored bars indicate the metrics with and without genotyping. In the hybrid row, gcSV and BlendSeq are evaluated based on 4x long reads plus 30x short reads, whereas Sniffles and cuteSV are evaluated based on 4x long reads only. For short reads, scores of gcSV, DRAGEN and GATK-SV are shown at 35x short reads, meanwhile, the yield of gcSV with 60x short reads is also listed as a reference (marked by “60x”). **b**, The yields of the callers under more restrict configurations for the –p parameter of Truvari (-p 90, 95 and 100 for dark, medium and light bars, respectively).

In the scenario of long read-based SV calling, with commonly used sequencing coverage (30x PacBio HiFi reads), gcSV shows a substantial improvement on the overall SV calling yield (F1-score with genotyping, F1-GT), i.e., 81.89% (Fig. 2a, first column), which is achieved by both higher sensitivity (Recall-GT, 77.93%) and precision (Precision-GT, 86.27%). This yield is about 20-38% higher than the that of Sniffles (61.47%), cuteSV (60.73%) and SVDSS (43.43%). Moreover, gcSV has higher yield for the discovery of SV events (F1-score without genotyping, F1-SV), i.e., 86.67%, about 6-11% higher than that of the other three tools, indicating that gcSV is good at reconstructing SV alleles. Meanwhile, the genotype concordance of gcSV is also highest among the tools (Supplementary Table 2, GT-concordance column), indicating its accurate SV genotyping markedly reduced the gap in yields before and after genotyping. For the other tools, such degradations are more pronounced. The GT-concordance of Sniffles and cuteSV were about 13-15% lower than that of gcSV and they showed even larger F1-GT degradations. For SVDSS, although its Recall-SV was fairly high (76.29%), the final yield was affected by its poor genotyping (GT-concordance: 53.98%) that Recall-GT and Precision-GT drastically dropped.

In the scenario of hybrid sequencing-based SV calling, we found the combination of 4x long reads plus 60x short reads were sufficient to achieve higher sensitivity, precision and overall yield as called by gcSV (respectively 56.70%, 76.13% and 64.99%, with genotyping, Supplementary Table 3 and Fig. 2a, second column) than that of Sniffles, cuteSV or SVDSS using 30x long reads. This result indicates that gcSV provides a highly cost-effective solution to SV calling with equal or higher ability to state-of-the-art tools. In contrast, Sniffles, cuteSV and SVDSS run on 4x long reads generated much lower yields (42.55%, 40.35% and 35.69% for F1-GT, respectively). Moreover, the results were compared to that of Blend-Seq^50^, a recently proposed hybrid SV-calling pipeline. Under the same data configuration (4x PacBio HiFi reads + 30x Illumina reads), the F1-GT of Blend-Seq is about 13% lower than that of gcSV (50.92% vs 64.38%, Supplementary Table 3). This result is reasonable since gcSV natively incorporates all reads throughout the SV calling pipeline, while Blend-Seq uses long reads for SV detection and short reads for genotyping independently.

In the scenario of short read-based SV calling, with 60x Illumina short reads, gcSV shows an outstanding sensitivity without loss of precision (Fig. 2a, third column). The overall yield (F1-GT: 41.39%) is markedly higher than popular tools such as Manta^10^, Lumpy^11^ and Delly^12^ (refer to the following subsections). To our best knowledge, this is the highest yield ever seen in the NIST-Q100 callset for short read data. Moreover, we compared gcSV to two state-of-the-art pipelines, GATK-SV^1^ and DRAGEN^30^ on a 35x Illumina dataset. Such pipelines combine multiple tools and/or take advantage of pan-genomes to enhance SV calling for specific SV types and/or genomic regions to optimize overall yields. In comparison, through a single algorithm and without assuming specific SV priors, gcSV achieved performances on par with or higher than the two integrated platforms. Especially, the precision-GT of gcSV is noticeably higher (gcSV: 76.15%, GATG-SV: 64.11%, and DRAGEN: 67.75%, Supplementary Table 4). This high precision could facilitate the detection of novel SVs in a reliable way, to serve the aim of many large-scale genomics studies. Moreover, it is feasible to integrate gcSV into other pipelines as a fundamental tool.

Additionally, gcSV shows more accurate SV reconstruction under stringent parameters (-P, 0.9, 0.95 and 1) of the Truvari assessment. These parameters govern that an SV call is considered as true positive only if the alternative allele has at least 90/95/100% overlap with ground truth. Fig. 2b shows the decay in F1-GTs between the stringent and the default settings (-P 0.7) is the smallest for gcSV, indicating that a larger proportion of SVs are precisely recovered by gcSV, i.e., the sizes and positions highly coincided with the truth (nearly exact match). It is worthnoting that, in the long-read scenario, the number of SV being perfectly recovered (-P 1) by gcSV is over 40% higher than that of the best runner-up (i.e., Sniffles).

The performance of gcSV is also suited to large-scale genomics studies. Evaluating on a computing server having medium hardware configuration (2 Intel(R) Xeon(R) Gold 6240 CPUs @ 2.60 GHz, 36 cores in total, 256 GB RAM, running CentOS operating system), gcSV showed affordable runtime for various data configurations (30x long reads: 0.83 CPU hours, 4x+30x/60x hybrid sequencing data: 3.35 and 7.94 CPU hours, 30x/60x short reads: 3.14 and 7.39 CPU hours, respectively) as well as relatively small memory footprints (30x long reads: 1.23 GB, 4x+30x/60x hybrid sequencing data: 4.76 and 8.39 GB, 30x/60x short reads: 4.24 and 8.29 GB, respectively).

In the following subsections, we analyze in detail the performance of gcSV on LRS, hybrid sequencing, and SRS data, respectively.

### The ability of gcSV for long read-based SV calling

Recently, LRS has played an increasing role in genomic studies for its ability to reveal large SVs and resolve variants in diverse genomic contexts. We compared gcSV to state-of-the-art LRS callers to find its performance much less affected by SV size, type and local repetitiveness.

Firstly, gcSV shows a higher ability to handle large SVs. We categorized all the ground truth SVs into six bins (i.e., 50-200bp, 200-500bp, 500-1kp, 1k-2kbp, 2k-5kbp and >5kbp). While other tools’ yields dropped drastically for large SVs (Sniffles at 500-1kbp; cuteSV at 500-1kbp; SVDSS at 2k-5kbp), gcSV’s yield remained high for SVs >2kb and it produced considerable yield for >5kbp where the results of other tools are poor (Fig. 3a, Supplementary Fig. 4a and Supplementary Table 5).

**Fig 3.**
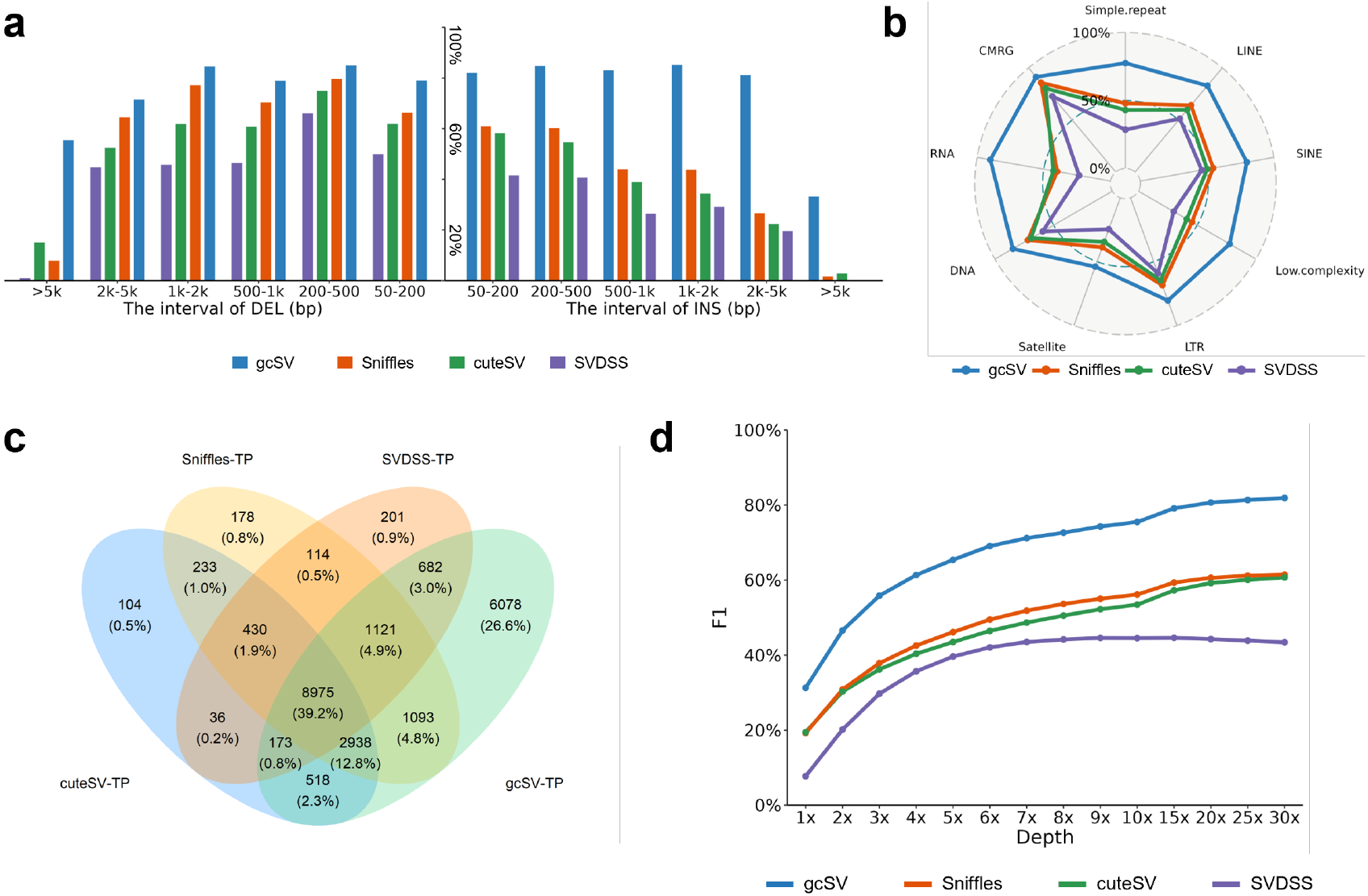
The ability of gcSV for long read-based SV calling. **a**, The F1-scores (with genotyping) of gcSV and state-of-the-art long read-based callers (Sniffles, cuteSV and SVDSS) for various types and sizes of SVs. Deletions (DEL, left) and insertions (INS, right) are binned by SV length as indicated on the x-axis. **b**, The F1-scores (with genotyping) of the callers in various kinds of genomic regions (according to the annotations of RepeatMasker and CMRG). **c**, The Venn diagram of the true positive calls produced by various approaches. **d**, The F1-scores (with genotyping) of the callers for various read coverages (1-30x).

Secondly, gcSV shows unbiased ability to SV types (Supplementary Table 5). We separately assessed the yields for deletion (DEL) and insertions/duplications (INS) events reported in NIST-Q100 callset. gcSV achieved not only higher, but also balanced yields for DELs and INSs. A similar trend was observed for SVDSS as well, indicating local assembly-based approach is robust to SV types. In contrast, Sniffles and cuteSV inferred much less INS than DEL (Fig. 3a). It is also worthnoting that gcSV has the ability to handle complex SVs such as the combinations of multiple events in various types (Supplementary Fig. 5).

Thirdly, gcSV detected SVs robustly in repetitive regions (Supplementary Table 6). We used RepeatMasker to bin the ground truth SVs into nine categories and separately assessed the callers in these regions (Fig. 3b). All callers exhibited balanced yields across most repeat categories, possibly due to the benefit of long reads as well as their tailored designs. However, for some extreme repeat categories, such as Simple Repeat, Low Complexity DNA and Repetitive RNA, the other callers had quite low yields (especially with genotyping) while gcSV maintained high yields. We also investigated the results on GIAB challenging medical genes (CMRG)^51^ and gcSV outperformed other callers as expected, suggesting its ability to handle difficult SVs.

Nevertheless, each tool contributed distinctive true positive SV calls (Venn diagrams, Fig. 3c), suggesting that higher yields could be made by heuristically combining the callsets of state-of-the-art callers. In addition, we observed that a large proportion of false positives/negatives calls of gcSV might have been resulted from inconsistent variant representation with the ground truth, while in fact the calls per se might have been highly similar to (or even exactly matched) the ground truth (Supplementary Fig. 6). To eliminate this ambiguity, we employed truvari with its refine module to re-assess gcSV calls and the result indicates that the adjusted yield obviously increased (F1: 95.58%, precision: 98.97%, Recall: 92.41%).

Further, we down-sampled the dataset to 1-25x (Fig. 3d, Supplementary Fig. 4b and Supplementary Table 2) to assess the yields on lower sequencing depths. We found that on 4x LRS data, gcSV already surpassed other tools on 30x LRS data in genotyping yield (F1-GT), suggesting a much more thorough exploitation of the LRS data by gcSV. Its yield saturated at 10-15x, indicative of a notable point to the cost-performance tradeoff.

### The ability of gcSV for hybrid sequencing-based SV calling

Given that gcSV produces significant yields with extremely low-depth LRS data, we investigated its capacity to jointly use LRS and SRS data to enhance SV calling while keeping the overall cost low. We evaluate 12 hybrid data configurations each combining 1-5x or 10x long reads with 30x/60x short reads. As shown in Fig. 4a and Supplementary Table 3, in all configurations, the yields of hybrid data without genotyping (F1-SV) are higher than that of LRS only, and more SRS certainly enhanced SV detection (i.e., 60x vs. 30x SRS). In particular, we noticed that with the integrated analysis of gcSV, SRS are strong supplements to extremely low depths LRS. The F1-GT of gcSV with 1x LRS+60x SRS (50.47%) came close to that of Blend-Seq with 4x LRS+30x SRS (50.92%), and so for 2-3x LRS+60x SRS configurations versus that of cuteSV and Sniffles with 30x LRS.

**Fig 4.**
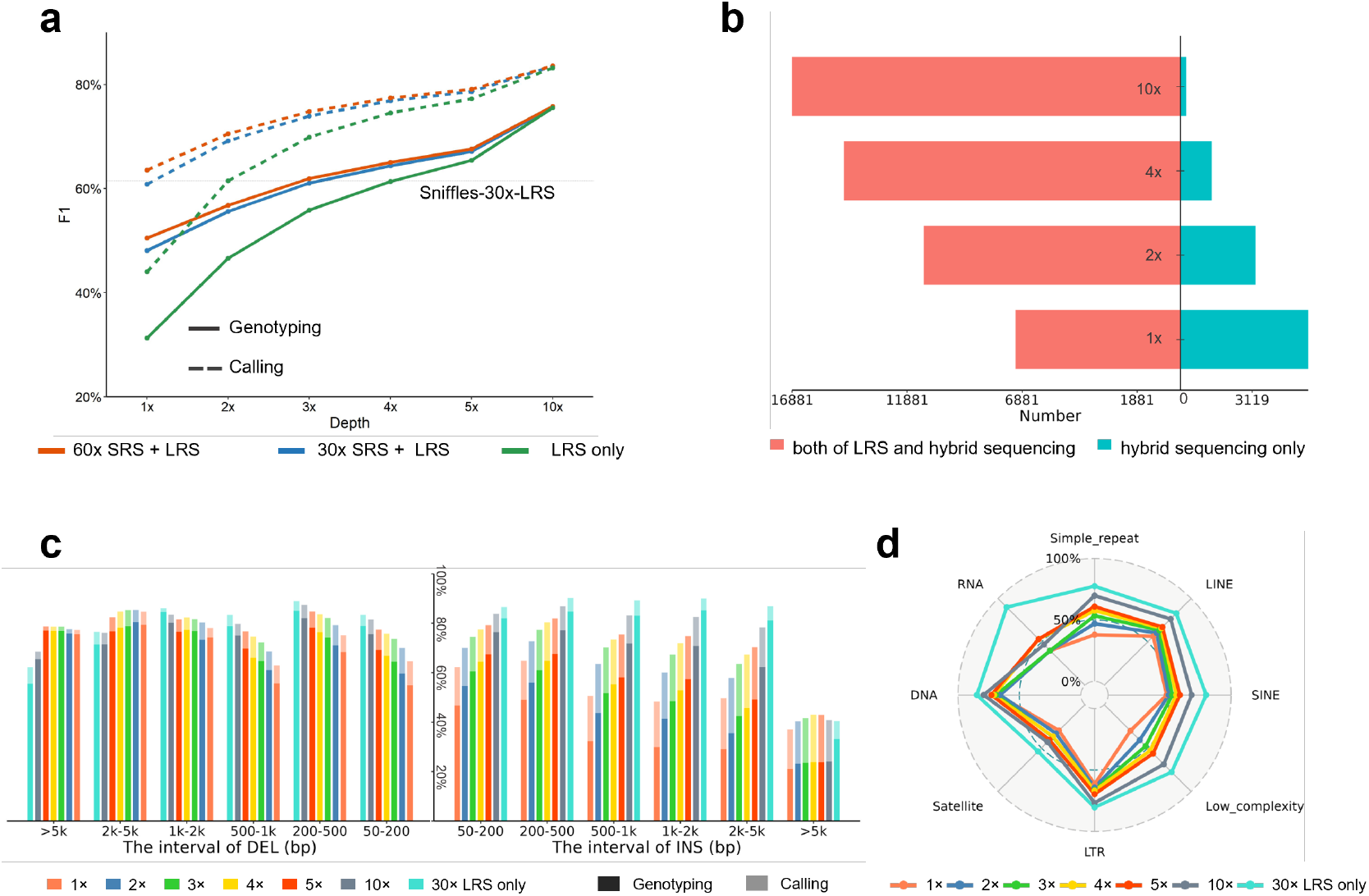
The ability of gcSV for hybrid sequencing-based SV calling. **a**, The yields of gcSV with 4x long reads plus various amounts of short reads (red: 60x, blue: 30x, green: none). The solid and dashed lines indicate the F1-scores with and without genotyping, respectively. The horizontal dashed line indicates the F1-GT of Sniffles with 30x LRS data. **b**, The comparison of gcSV calls using hybrid sequencing data and long reads only. The SV calls of various coverages (1x, 2x, 4x and 10x) of long reads with and without 60x short reads are compared in pairs. The red (left) and cyan (right) bars indicate numbers of true positive calls produced by both data configurations and by hybrid sequencing data only, respectively. **c**, The yields of various hybrid sequencing configurations for various types and sizes of SVs. Deletions (DEL, left) and insertions (INS, right) are binned by SV length as indicated on the *x*-axis. The yields of 1–10x long reads plus 60x short reads are shown in various colored bars (also compared with that of using 30x long reads only). The dark and light colors indicate the F1-scores with and without genotyping, respectively. **d**, The F1-scores (with genotyping) of various configurations (1x, 2x 3x, 4x 5x, 10x long reads plus 60x short reads and 30x long reads only) in various kinds of genomic regions (according to the annotations of RepeatMasker and CMRG).

This supplement effect is evident in the comparison between (true positives in) the callsets based on hybrid data and LRS data of the same depths (Fig. 4b). For 1-2x LRS, a large proportion of SVs were distinctively called with hybrid data. However, this effect attenuates for >4x long reads, because gcSV were able to discover SVs with very low numbers of supporting long reads. Nevertheless, the contribution of high-depth SRS is non-neglectable. The above results suggest that the unified SV calling framework of gcSV maximizes the power of hybrid sequencing data in detecting SVs. Using gcSV, 2x LRS+60x SRS could be a feasible solution for highly cost-effective SV calling in large-scale genomics studies. Meanwhile, 4-5x LRS+60x SRS is also an option to higher yield.

The yields of various hybrid configurations on SV sizes and types are shown in Fig. 4c and Supplementary Table 7. gcSV on hybrid data kept its yield up to moderate sized SVs (SV < 5kbp), although slightly tilted toward DELs, because INSs are more difficult to reconstruct with short reads. Despite the drop in yields for very large (>5kbp) insertions/duplications, very large deletions were more likely to be found from hybrid data than from pure LRS data. A plausible explanation is that deletion signatures of short read are easier to capture and be utilized for the local assembly. Moreover, subsequent SV filtration and genotyping benefited from the higher statistical power provided by the abundance of short reads.

The yields in various genomic regions are shown in Fig. 4d and Supplementary Table 8. For most repeat categories, the F1-SV yields of various hybrid configurations are quite close to that of 30x LRS. The drop in repeat handling for hybrid data is only pronounced in highly repetitive regions, due to the inherent insufficiency in read length for even high-depth SRS to resolve these regions (refer to the next subsection for more details).

### The ability of gcSV for short read-based SV calling

We benchmarked gcSV on three HG002 SRS datasets in 60x, 30x and 15x coverages, and compared it with Manta, Delly and Lumpy, three short read-based SV callers commonly employed in state-of-the-art pipelines. Two integral platforms DRAGEN and GATK-SV were further drawn into comparison. Due to compatibility issues, for the DRAGEN platform, we used the 35x HG002 dataset as in its published study^52^ and quote its performance therein.

The results indicate gcSV achieved the highest overall yields without genotyping (F1-SV) for all datasets, outperforming all standalone tools by a considerable margin (Fig. 5a and Supplementary Table 4), while the yields of DRAGEN and GATK-SV pipelines were closer to that of gcSV. Considering the strong interference of repeats to short read mapping, we separately assessed the sensitivities and precisions of the SV callers in NIST-Q100 alignment “easy” and “hard” regions^53^. The results (Fig. 5a and Supplementary Table 9) indicate that in easy regions, gcSV was highly reliable (95.73%-97.08% precision-SVs for various depths), and achieved significant gains in sensitivity over all competitors (about 11% and 20% higher than DRAGEN and GATK-SV, respectively). In hard regions, gcSV had a performance on par with DRAGEN and GATK-SV (Fig. 5a), although in absolute terms all of them are relatively low. For gcSV, this was attributed to the difficulties in realigning reads to multiple candidate alleles in the presence of complex repeats, even when the SV alleles themselves were finely reconstructed. Thus, the statistical power of both SV detection and genotyping was affected. This issue can be largely addressed by the inclusion of LRS data (refer to Supplementary Fig. 7 for illustrations).

**Fig 5.**
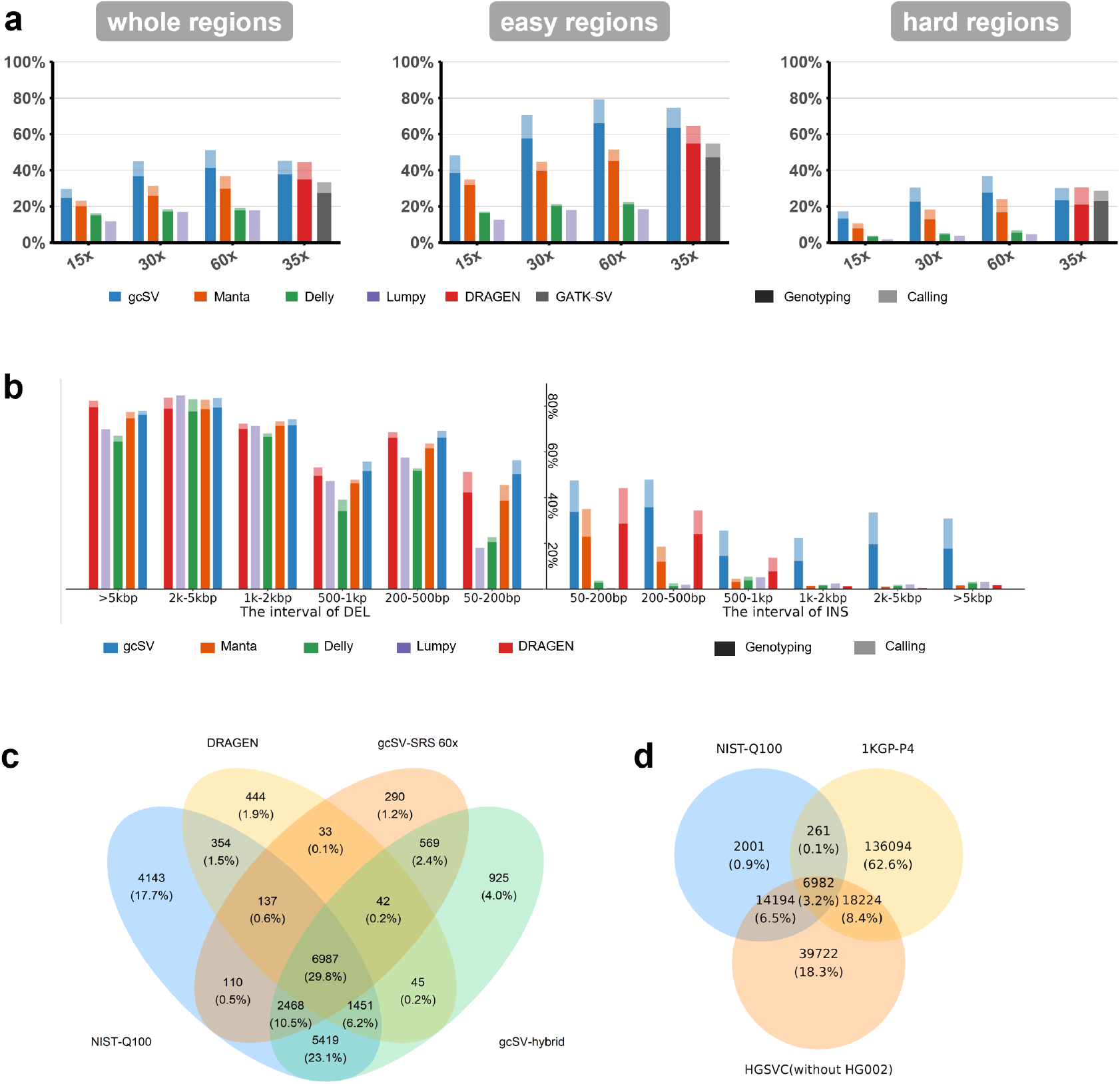
The ability of gcSV for short read-based SV calling. **a**, The yields of the callers (gcSV, Manta, Delly, Lumpy, DRAGEN and GATK-SV) for various (whole, hard, easy) genomic regions using various coverages of short reads (15x, 30x, 60x, 35x). The “whole” regions include all the regions defined by HG002 ground truth. “Hard” and “easy” regions are regions hard and easy for short reads to be aligned, respectively, as designated in previous studies^53^. **b**, The yields of the callers for various types and sizes of SVs. Deletions (DEL, left) and insertions (INS, right) are binned by SV length as indicated on the *x*-axis. The dark and light colors indicate the F1-scores with and without genotyping, respectively. It is also worthnoting the yield of DRAGEN is based on 35x SRS dataset which is directly quoted from its own study. For other callers, their yields on 60x SRS dataset are shown. **c**, The Venn diagram of SV calls of gcSV (using short reads and hybrid sequencing data, respectively), DRAGEN and HG002 ground truth. **d**, The Venn diagram of HG002 ground truth, 1KGP Phase4 official callset and HGSVC official callset (the SVs of HG002 are excluded beforehand).

Further, we assessed SV yields in various size and type categories and found that gcSV had a more balanced performance (Fig. 5b and Supplementary Table 10). While all tools exhibited high yields for DELs, the yields of INSs were uniformly low, in consistence with previous observations^30, 50^. However, gcSV stood out as the only tool yielding a considerable amount of large insertions (>1kbp, see Supplementary Fig. 8 for examples). It is also worth noting that gcSV’s ability to detect large deletions is slightly reduced, although still comparable, relative to that of DRAGEN. As most of the deletions missed by gcSV were from highly repetitive regions, which are exceedingly challenging for single-reference-based approaches, we speculated that DRAGEN’s advantage likely stems from the joint use of its pan-genome reference (yet unpublished) and certain components specifically designed for such events (see Supplementary Fig. 9 for examples).

The callsets of gcSV and DRAGEN (both 35x SRS) were compared (Venn diagram, Fig. 5c), revealing unique true positives in each. This highlights the complementary nature of different methods and the potential of integration to address complex short-read alignments in varied genome contexts. On a different note, supplementing short reads with limited long-read data (4x LRS+60x SRS hybrid) significantly boosted gcSV’s yield. This improvement arises primarily because long reads lend sufficient confidence for SV calling by distinguishing the true positive alleles from multiple ambiguous candidates. We observed from the intermediate results that, based on short reads, gcSV detected over 89.18% of the ground truth SV loci, but only 35.4% can be successfully called. However, with the help of only one or a few long reads, the proportion increased to 67.5%.

Lastly, we evaluated the potential of integrating a pan-reference for SV calling. As DRAGEN did not disclose its pan-reference details, we drew an indirect comparison between NIST-Q100 callset versus the official SV callsets of 1000 Genomes Project (1KGP) Phase4^6^ (3,202 samples, 173,366 SVs in total) and Human Genome Structural Variation Consortium^54^ (HGSVC, 12 samples after excluding HG002, 110,747 SVs in total). We observed that 2,001 of the SVs in NIST-Q100 callset were not covered by the other callsets (Fig. 5d), suggesting that a limited usage of pan-reference made by 1KGP and HGSVC callsets, which underscores the pivotal role of *de novo* SV detection by gcSV in genomic studies.

### The re-analysis of large-scale genomics datasets with gcSV

Having shown the performance on SRS data, we interrogated gcSV’s ability in re-analyzing the many existing large-scale genomics datasets. We implemented gcSV on the 1KGP SRS dataset (3,202 samples, on average 30x short reads) and compared its results (gcSV-callset) to the official callset of 1KGP Phase4 (1KGP-callset), which was produced by a combined pipeline of GATK-SV, SVTools and Absinthe. Overall, gcSV reported 32% more SV events (228,025) than that of the 1KGP-callset (173,366), and 93,242 of them were novel (not reported in 1KGP-callset). Moreover, an 18% increase in per sample SV number (11,405 vs 9,639) was observed, consistently across populations (Fig. 6a).

**Fig 6.**
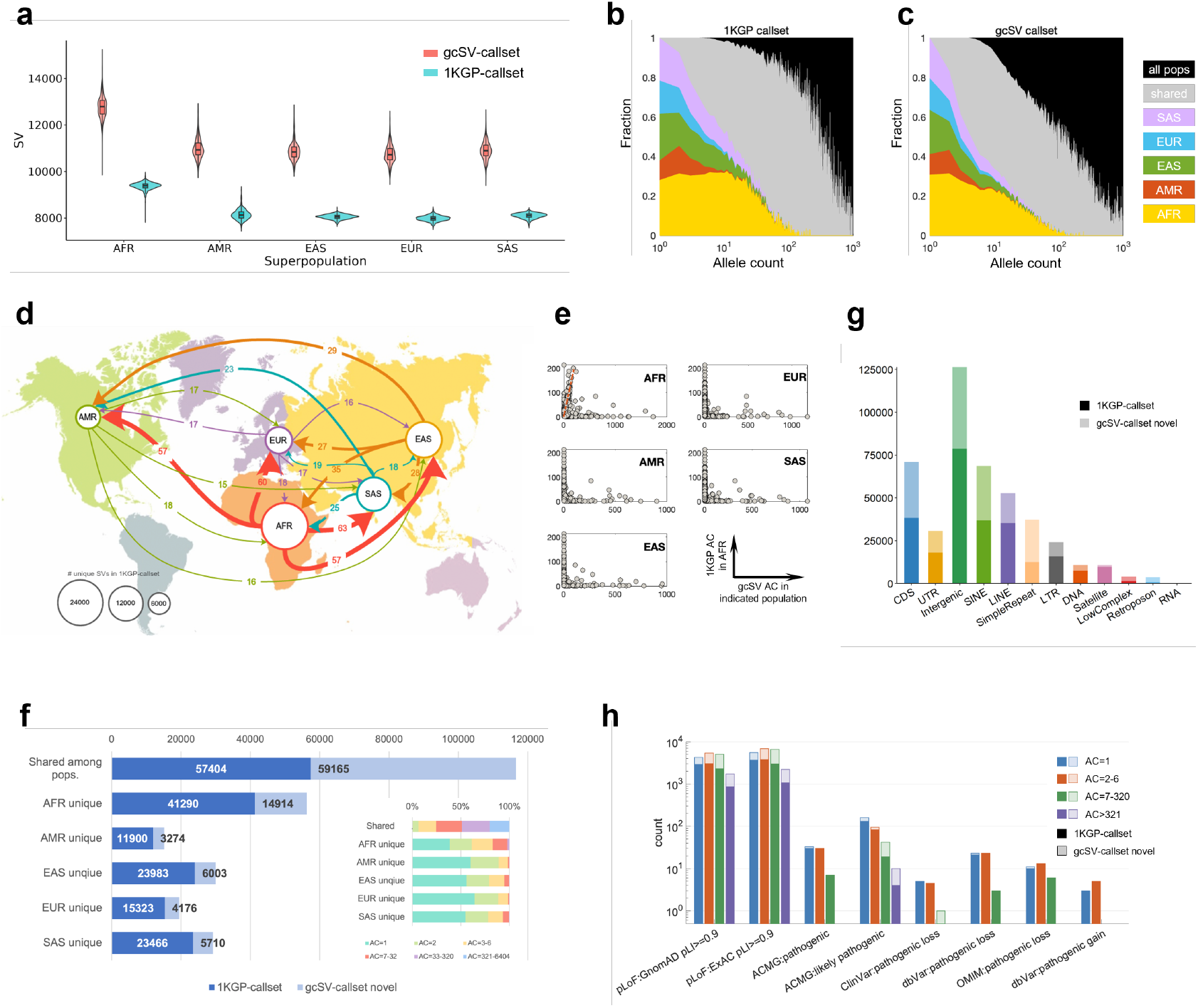
The re-analysis of 1KGP datasets by gcSV. **a**, Distribution of population-wise per-sample SV numbers in gcSV-callset and 1KGP-callset. The abbreviations for superpopulations are: AFR – African, AMR – American, EAS – East Asian, EUR – European, SAS – South Asian. **b&c**, Distributions of population unique and shared SV alleles. Shared represents alleles shared by at least two but not all superpopulations in 1KGP-callset (b) and gcSV-callset (c). **d**, Shifts in SV allele distributions between superpopulations. Arrows indicate that alleles deemed unique to the source superpopulation by 1KGP-callset are regarded common (frequency > 0.05) in the destination superpopulation by gcSV-callset. The circle size scales with the number of alleles unique to each superpopulation in gcSV-callset. The arrow width and the number marked on each arrow indicate the number of alleles re-discovered as common in the destination superpopulation. **e**, Allele counts of AFR unique SVs in 1KGP-callset vs. those of the same SVs in gcSV-callset in the same superpopulation or in another superpopulation, as indicated in each panel. Red dashed line in the first panel shows 1:1 ratio. **f**, Numbers of population unique and shared SVs in 1KGP-callset (dark blue) and distinctly identified in gcSV-callset (light blue). Inset shows the allele count distribution of SVs distinctly identified in gcSV-callset. **g**, Functional/genome region annotation of SVs in the 1KGP-callset (dark colors) and SVs distinctly identified in gcSV-callset (light colors). **h**, Phenotypic annotation of SVs in the 1KGP-callset (dark colors) and SVs distinctly identified in gcSV-callset (boxed light colors). For each annotation group, SVs are partitioned into four allele frequency bins.

We investigate the population-wise SV distributions, as they are routinely used to construct population genomic backgrounds and provide important clues for population genetic studies. Compared to 1KGP-callset, gcSV-callset displayed more smooth and faster decays in the number of population unique alleles with increasing allele counts (Fig. 6b&c). For all superpopulations, this updated trend revealed a discernible bi-modal distribution pattern, with the first mode presumably representing alleles undergoing neutral processes, and the second mode likely encompassing common alleles selectively retained in specific populations. Next, we inspected in details changes in allele distributions in gcSV-callset. Remarkably, gcSV pervasively recovered SV alleles previously regarded unique to a certain superpopulation in samples from other superpopulations (Fig. 6d). For example, 0.29% of African-unique SVs in 1KGP-callset were found to be of common alleles (frequency ≥0.05) in at least one other superpopulation, and 0.17% were found to be common in all five superpopulations. In addition, 1.52%∼2.84% of the population-unique SVs had their population AFs strongly elevated (Fig. 6e, Supplementary Fig. 10). Second, of the 93,242 SVs discovered solely by gcSV, 63% were shared among populations. However, the rest were regarded population unique, resulting in 24–36% increases of population-unique SVs for each individual superpopulation (Fig. 6f). Although over 80% of the newly identified population-specific SVs were extremely rare (AF<0.002, Fig. 6f inset), given gcSV’s high precision, most of them could be plausible. The newly discovered alleles and updated frequencies for existing alleles can provide resources for many association studies.

With regard to genome-wide distribution of SV alleles, the most pronounced increases in the number of discovered SVs were observed for tandem repeat (TR) regions (∼ 2-fold increase vs. gcSV-callset), along with other repetitive elements such as SINE, LINE, LTR and retrotransposons (Fig. 6g), which are still regions hard for state-of-the-art SRS-based callers to detect. While short TRs can be relatively reliably called from SRS data^55, 56^, gcSV were able to discover kilo-base scale TR expansions, extending TR unit variations to > ±500 copies relative to the reference genome and revealing stable fractions of population-unique TR alleles (Supplementary Fig. 11).

Lastly, we performed functional and clinical-related annotations for all gcSV-distinctive SVs by seven resources, i.e., GnomAD^57^, ExAC^58^, 1KGP4^6^, ACMG^59^, OMIM^60^, dbVar^61^ and ClinVar^62^ (Fig. 6h). A considerable number of gcSV-distinctive SVs were annotated by significant terms: 7,312 and 9,639 were annotated as pLoF by GnomAD and ExAC, respectively; and 71 were annotated as pathogenic and likely pathogenic by ACMG. In OMIM, dbVar and ClinVar databases, three large deletions (10–40 kb) on chromosomes 3, 7 and X were identified as extremely rare alleles in EUR and AFR superpopulations as related to retinal dystrophy, spastic paraplegia and congenital hypothyroidism, respectively; and a 50 bp insertion, common to all populations, was found to be related to primary ciliary dyskinesia (Supplementary Table 11).

## Discussion

The characterization of SVs is a cornerstone of genomic research. Despite considerable efforts in developing computational tools for SV calling from HTS data, the sensitivity and precision of state-of-the-art tools still fall short of achieving ideal performance. Furthermore, in the emerging T2T genomics era, SV detection for LRS data continues to undergo active methodological refinement to address recalcitrant genomic regions. Additionally, devising a practical and cost-effective solution for SV calling remains an unresolved challenge, particularly for million-scale genomic cohort studies. gcSV introduces a novel alignment-based SV calling approach that employs context-driven SV loci recognition and repeat-resilient SV reconstruction within a unified framework to address these challenges cohesively. Benchmarks demonstrate that gcSV transcends current performance thresholds in long read-based SV calling with 20-38% yield improvement over state-of-the-art callers. On short read data, gcSV also improves the essential ability of SV calling, outperforming state-of-the-art pipelines that integrate multiple existing tools and pan-genome reference (Fig. 2a).

Increased sensitivity constitutes the critical determinant in achieving yield optimization. With the fine modeling of SV signatures in various contexts of genomic local repeats, gcSV captures novel evidences such as dense CIGARs in reads from vanishing alignment signals in highly repetitive regions. Next, the adaptive read clustering approach finely classifies reads to the correct SV event and enhances SV signatures. Both steps ensure that SV sites can be comprehensively discovered from weak signals and with low read numbers, and a larger fraction of reads are reasonably retained for further analyses than in previous methods. Consistently, gcSV achieves comparable yields to other LRS callers with over 85% reduction in LRS depths (4x vs 30x, Fig. 3d). Meanwhile, it generates a considerable number of difficult SVs (such as large insertions, Fig. 5b) which remain elusive to state-of-the-art SRS callers like DRAGEN and GATK-SV. This results in more balanced yields for insertions and deletions, achieving a more representative depiction of the structural diversity in individual human genomes.

The precision of SV calling is also rigorously refined by fine-grained signature clustering and local assembly-based reconstruction. The former step uses iterative read clustering to comprehensively collect interspersed SV-spanning reads. The outcome is highly purified read clusters devoid of confounding signals and well-resolved SV breakpoints in close proximity. The next step implements heuristic walk-based local assembly in de Bruijn graph framework and then statistically infer SV alleles from re-alignment signals. For LRS data, previous work (e.g., SVDSS) consistently supports that local assembly enhances SV reconstruction accuracy. For SRS data, this step enables the “blind” generation of highly repetitive SV candidates. Combining realignment coverage information, it is able to pinpoint repetitive SVs, a task so far reckoned intractable with SRS data. Benchmark suggests gcSV reconstructed SVs with precision on the single-base level, yielding over 40% more SVs with sequences matching the ground truth (Fig. 2b).

Besides elevated yields on LRS and SRS data alone, gcSV offers a distinct opportunity for cost-effective SV calling, i.e., using 2-4x LRS plus 30-60x SRS data to achieve equal or higher yields than state-of-the-art tools (Fig. 4a). This is made possible by the unified representation and analytical framework of long and short reads. gcSV’s natively integrative approach allows both read types to play complementary roles (Fig. 4b). For example, in SV site recognition, long and short read-specific signals are consolidated to reduce false negatives. Subsequently during allele reconstruction, the few long reads provide long-range path association while the abundant short reads offer high statistical power for contig generation and inference. Moreover, in regions insufficiently covered by low-depth LRS data, SVs can be reliably called from SRS data to supplement the overall callset.

gcSV is readily integratable into existing genome analysis platforms. As a demonstration, we used gcSV for the pervasive re-discovery of SVs in the 1KGP dataset. gcSV reported a surplus of 32% high-confidence SVs than the official callset, with 12% of novel SVs deemed potentially pathogenic (Fig. 6a&h). Moreover, we observed marked shifts in SV allele frequencies and their population specificities (Fig. 6b-f). Lastly, given its targeted strategy toward repetitive genome regions, gcSV identified a notable fraction of repeat-associated SVs (Fig. 6g), providing a fresh view for studies of tandem repeat variations and their phenotypic consequences.

Importantly, several technical issues of SV discovery require further exploration. Firstly, very large SVs over 10kbp, as well as specific SV types or extremely complex SV clusters, present hurdles necessitating the incessant improvements in local assembly methods. Secondly, variable SV representation, as a result of the ambiguity in contig-to-reference alignments, has remained an issue for gcSV. To minimize its impact on downstream annotation and analysis, appropriate alignment scoring or novel representation methods (such as the graphical pan-genome) are anticipated. Lastly, our benchmark suggested that many SV sites could have been detected by gcSV with short reads, but were rejected due to low confidence levels. Intensive research is required to effectively utilize these copious yet obscure signals. In addition, although we have demonstrated gcSV’s power on hybrid data, it remains an open question regarding the optimal sequencing strategies and parameter configurations for subsequent SV calling.

## Online methods

### Repeat context-driven SV site recognition

We used long and short read sequencing data from Human Pan Reference Consortium (HPRC)^63^ to explore the distributions of alignment signatures around SV loci. Multiple kinds of colinear and non-colinear signatures were investigated, including sequence divergences in alignments (SDAs), large clippings (CLPs) and discordant read pairs (DRPs, only for short reads). Herein, SDA is defined by both of dense sequence divergences (mismatches and indels) shown by their CIGAR information and split alignment. We combine such signatures primarily due to that few of the reads (for both long and short) have split alignments in practice. Moreover, such signatures are collected from full-length alignments and can be handled in a unified approach.

Signature distributions were constructed to measure the local intensity of signature in various contexts of repetitiveness which was implemented in the following steps. 1) The reference was divided into 600 bp size windows with 300bp overlaps. For each window, a measure of local repetitiveness, *R*, was calculated as the maximum length (*R*) of repeated *K*-mers in the window:

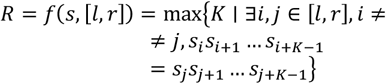

where, *s* is the reference and [*l, r*] is the coordinates of the window. *R* rises with local sequence repetitiveness. 2) The assembly-based SV callsets of the HPRC samples were used as ground truth to mark the windows involved in SV regions (termed as SV-windows). For each SV-window, the numbers of reads having SDA, CLP and DRP signatures (termed as #SDA, #CLP, #DRP) were separately counted. 3) Two sets of two-dimensional (repetitiveness and signature) empirical distributions based on all reads within SV-windows were constructed for short- and long reads, respectively. The distribution of DRP was only applied for short reads. 4) The distributions of signatures in non-SV-windows were also built in a similar way to serve as background distributions.

These distributions depict the intensity of the signatures as well as their divergences under the effects of repeats, which is important to model the read alignments in various genome contexts. Mainly, five keypoints were observed as following.

1) SDA is informative to SV detection, moreover, it shows different characteristics for LRS and SRS data (Supplementary Fig 1b, first row). For long reads, SDA signal is strong and evenly distributed for all the degrees of repetitiveness. This is due to that most of the SVs are shorter than read length, so that the events can be directly reflected by long indels in the alignment records (CIGARs), meanwhile, for large SVs the spanning reads can also be split-aligned with the high mappability. For short reads, SDA has a negative correlation with the degree of repetitiveness, and is indicative to SVs in highly repetitive regions. This is mainly due to the fact that there are usually small variants (SNVs and indels) among repeat units. In this situation, the spanning reads are colinearly aligned under the strong effect of repeat, but having systematically increased edit distances. This provides new and critical evidence to SV detection in repetitive regions, however, not being fully considered by state-of-the-art tools.

2) In both of LRS and SRS data, the numbers of SV windows having high CLP (e.g., #CLP > 4) reduce with the increasing repetitiveness, indicating the attenuation of the signature in high repetitive regions (Supplementary Fig 1b, second row). This is not surprising since the SV alleles have higher similarity to local repeats so that the spanning reads can be fully aligned there with only small divergences. Moreover, in long read data, the number of SV windows having CLPs is much lower than that of SDA, because only very large SVs could trigger clippings during long read alignment and such events are rare. However, CLPs are non-neglectable since they are the only signatures available in such cases. The situation of short reads is quite different that CLP is the most informative signature. This stems from that the read parts involving SV events have poor mappability so that aligners fail to produce split alignments and can only leave clippings around SV breakpoints.

3) The intensity (the number of SV windows) of DRP is much lower than that of CLP (Supplementary Fig 1b, third row). This may be caused by many factors such as alignment scoring, relative read positions and local repetitiveness. More importantly, DRPs should be higher weighted during short read-based SV detection since they are not easy to occur by chance, especially with the colinear preference of short read aligners. It is also worthnoting that DRP signature also attenuates in highly repetitive regions.

4) Although attenuation exists, the sum of various signatures indicates that most of SV windows still contain plenty of signatures (Supplementary Fig 1b, fourth row). On the contrary, a small proportion of the SV windows do not have signatures regardless of the repetitiveness, especially for long reads. We found that most of these windows are near to the breakpoints of large SVs and the corresponding read parts are aligned with no large indels or clippings, but a number of small divergences like mismatches. However, this may not affect SV detection seriously since such SVs may span a number of SV windows and enough signatures can be captured in nearby windows.

5) As indicated by background distributions (second and fourth columns of Supplementary Fig 1b), SV signatures can also be occasionally detected in non-SV windows. This is mainly due to the outlier reads having abnormal sequencing errors or insert sizes which embody SV-like signatures. However, none of the three signature categories show specifically high intensity in single non-SV windows. Thus, this issue can be largely addressed by fully considering various kinds of signatures. It is also worthnoting that such cases more often happen in highly repetitive regions. Combining with the signature attenuation in SV regions, the windows in such regions could become indistinguishable which potentially affect SV detection. A feasible solution is to straightforwardly mark the highly repetitive regions (e.g., *R* > 40) as candidate sites during SV detection, as their absolute number is not very high (1.46% of human reference genome) and the computational cost is affordable. Moreover, this operation also makes sense since many SV hotspots are in such regions.

Based on these distributions, a model was built for the recognition of unknown SV windows. Both of the signature distributions of non-SV and SV windows are considered to achieve high sensitivity while prevent too-many false positives. Mainly, the model defines two sets of rules for long and short reads, respectively. For long read data, given the statistics of SDA and CLP signatures, a genomic region is recognized as an SV candidate site if it meets the condition that there are more than *T*_*LR*_ reads in the region having either >30 bp indel(s) or >300 bp clippings recorded by their CIGARs. For short read data, given the statistics of SDA, CLP and DRP signatures, a genomic region is recognized as SV candidate site if it meets one of the four conditions: 1) the *R* value is higher than 40; 2) the number of DRP reads is higher than MAX(*T*_*SDA*_, 2); 3) the number of SDA reads is higher than MAX(*T*_*CLP*_, 2); 4) the number of DMI reads is higher than MAX(*T*_*DRP*_, 2). Here, the default values of the thresholds *T*_*SDA*_, *T*_*CLP*_ and *T*_*DRP*_ are set to 10% of local read depth. These rules are on the design of keeping low type II error while preventing from very high type I error. Using these rules, the sensitivities of SV site recognition on HG002 benchmark are 98.14% (19,234 sites) for long reads and 89.18% (17,478 sites) for short reads.

### Fine-grained SV signature clustering

Although SV loci can be recognized with the repeat context-driven modeling, the signatures are still complex under the effects of the various types, sizes and breakpoints SV events. The SV spanning reads can be interspersed in the reference due to the multiple SV breakpoints (such as large deletions, inversions and translocations) or repeats (such as mobile element insertions). The signatures of multiple events could also intertwine and present highly complicated patterns in heterozygous or adjacent SV sites. Moreover, the reads having unusual sequencing errors or insert sizes also bring false positive signatures. Under such circumstance, it is still non-trivial to collect the signatures belonging to the same event comprehensively and precisely to provide purified read clusters for further processing. This may also lead to the failure or mistake of SV allele reconstruction.

To solve this problem, gcSV uses a two-phase approach to implement refined SV signature read clustering. The first phase (breakpoint phase) is to make highly refined signature clusters around SV breakpoints (refer to Supplementary Fig 2a for a schematic illustration). The major idea is to treat SV breakpoint as a latent random variable and composes a probability distribution for it using the alignments of the clustered reads. Further, the probability of the clustered reads belonging to the breakpoint can be estimated and the unlikely ones are removed to obtain more purified clusters. The second phase (event phase) is to connect the clusters around various breakpoints under the contexts of genomic positions or linking information implied by read pairs or split alignments (refer to Supplementary Fig 2b for a schematic illustration). This helps to comprehensively collect the signature reads of the same SV event which is useful to the following step, i.e., SV allele reconstruction and genotyping. Some of the implementation details are as following.

In the first phase, gcSV collects all the reads having SV signatures from the recognized SV windows (and also the nearby windows to avoid false negatives). Further, the reads are categorized by their signatures (SDA, CLP and DRP) and initially clustered by their mapped positions. For each of the initial clusters, a probability distribution for the latent SV breakpoint is heuristically composed regarding to the category of signatures and the SV involving parts of the reads. Further, gcSV uses the distribution to test each of the reads and reject the ones unlikely belonging to the breakpoint. Purified clusters are the composed and gcSV tries to use the rejected reads to re-compose new clusters in an iterative manner. In details, for various categories of clusters, the refinement is done as follows.

For an SDA cluster, the whole reads are seen as the SV involving parts. gcSV projects all the read bases to reference by read mapping positions (Supplementary Fig. 2a, the first and second panel). A discrete distribution is then generated on the read-projection region, i.e., for each position of the region, the probability is defined as the number of the bases projected to it (normalized by the total number of bases). Further, for each of the reads, gcSV retrieves the probability for its centroid (i.e., the mapping position of the middle point of the read in reference) for refinement (keeping/rejection).

For a CLP cluster, the clipping parts are seen as the SV involving parts. gcSV compose a virtual local reference to map all the clipped read parts to it (regardless of the sequence, Supplementary Fig. 2a, the third panel). The distribution is then generated by the numbers of mapped bases. Further, for each of the reads, gcSV retrieves the probability of its clipping position for refinement.

For a DRP cluster, the junctions between paired read-ends are seen as the SV involving parts. gcSV also compose a virtual local reference to calculate the middle points of the junctions and generate the distribution by the numbers of mapped bases (Supplementary Fig. 2a, the fourth panel). Further, for each of the reads, gcSV retrieves the probability of its junction points in refence for refinement.

In the second phase, gcSV uses a greedy approach to connect the refined clusters. In details, it lists and scans all the clusters from upstream to downstream. The clusters in nearby regions and/or being linked are straightforwardly combined, regardless of their categories. Furthermore, the combined clusters are then tested by repeat context-based model again to determine the candidates for SV-allele reconstruction. All the reads in the candidate clusters are separately collected for further processing.

This approach is useful to handle the SVs having multiple and ambiguous breakpoints to comprehensively collect involving reads. For example, an inversion event typically has two breakpoints and four involving windows and this approach can directly cluster them at once (Fig. 1c). It is more notable for the event related to interspersed repeats (Supplementary Fig. 2, bottom panel) such as mobile element insertion (MEIs). In such cases, the reads involved in the SV-alleles be intricately mapped to many repeat copies which is difficult to collect. However, the candidate windows are either nearby or linked so that the greedy approach can cluster all of them transitively. It is also not worrying that the greedy approach may mistakenly combine the clusters of various SV events together, since the cluster portioned from the same initial cluster are recorded beforehand and not allowed to connect. Moreover, the histogram of the distances between neighboring SV events (based on HPRC callset) indicate that SVs are overall sparsely distributed along the genome so that most of the clusters contain only one event. Meanwhile, gcSV also has the ability to reconstruct multiple SV-alleles and discover the events simultaneously if the reads of multiple events are combined in one cluster.

### Heuristic walking-based SV reconstruction

SV-alleles themselves could be also highly repetitive and similar to flanking regions so that it is difficult to reconstruct precisely. This is still a hard problem for state-of-the-art tools during calling and gcSV solves it by also using a novel generative approach which is also implemented in two phases.

In the first phase, gcSV implements a heuristic de Bruijn graph-based contig generation to produce candidate SV-alleles. The novelty is that, with coverage-based rules, the contigs can be generated solely based on the walk along the graph if the repeats cannot be solved with limited read length, which is implemented by the following three steps (Supplementary Fig. 3).

1. For an SV signature cluster, gcSV extends its genomic region by 500bp upstream and downstream, respectively, and collects all the involved reads (both short and long) as well as the local reference. The reads are then indexed by homopolymer-compressed (HPC) k-mers. Further, a de Bruijn graph is built using all the k-mers of the reads through k-mer counting. Virtual SV breakpoint(s) are then made by a position-based heuristic and gcSV selects two unique k-mers upstream and downstream the breakpoints as the source- and sink nodes, respectively.
2. gcSV use a depth-first search method to compose one or more candidate paths through the graph from the source to the sink nodes. Using only short reads, gcSV assigns a maximum number of allowed coverages (i.e., a threshold for times of visiting) for all the nodes of the graph and conduct depth-first search to generate all the legal paths. Herein, the visiting thresholds for the nodes are determined by the corresponding k-mer counts of the input reads beforehand which reflects the read coverages as well as the intrinsic repetitiveness of the k-mers. Moreover, a path is considered legal only if it starts and ends at the source and sink nods and all the passing k-mers have lower coverages than the max allowed number. Using this approach, the contigs (candidate SV-allels) are “blindly” generated with an affordable computational cost under the control of read coverage implicitly. It is also worthnoting that, if long reads are available, the candidate paths are generated under a more strict control, i.e., for each of them, all the nodes in the path must appear in the same long read and maintaining the same topological order.
3. All the generated paths (by both of long and short reads) are converted to candidate contigs. In this context of assembly, any pair of the contigs is then composed as a hypothesis under the diploid assumption of current version of gcSV. Moreover, gcSV separately implements k-mer counting for all the reads and each of the contigs for further processing.

In the second phase, gcSV uses k-mer frequency-based approaches to filter the contigs and separately tests each of the remaining ones to infer the most likely SV-allele(s). Herein, k-mer-based statistical approach is adopted instead of base-level read realignments due to that the reads usually cannot be confidently aligned to the repetitive SV-alleles. Moreover, as the k-mer statistics indirectly reflect the read coverage on the contigs, the approach is suited to distinguish the ambiguous candidates having various copy numbers which are also the common cases of variable number tandem repeats (VNTR). The implementation is done by the following two steps (Supplementary Fig. 3).

1. gcSV filters the contigs and re-maps the reads at first. The filtration is simply done by the comparison of k-mer sets between the contigs and the reads. That is, gcSV assumes that a contig is confident only if all its internal nodes can be explained by the reads and discards all the candidates which have one or more k-mers not being covered by the k-mer set of the reads. After that, gcSV tries to remap each of the reads to remaining contigs by non-gap alignment and records the ones if the sequence of the read can be fully contained by the contig.
2. gcSV assumes each of the remaining contigs as a candidate haplotype and tests each pair of the haplotypes to infer the most likely alleles to the site. Given a pair of haplotypes (*H*_1_, *H*_2_), its likelihood is measured by the following equation:

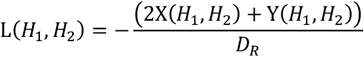

Herein, *D*_*R*_ indicates the read coverage of the dataset, X(*H*_1_, *H*_2_) indicates the number of reads cannot be remapped to any of *H*_1_ and *H*_2_, and Y(*H*_1_, *H*_2_) indicates the similarity between the k-mer counts of (*H*_1_, *H*_2_) and the reads which is computed as following:

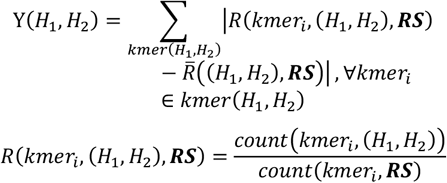

*kmer*(*H*_1_, *H*_2_) indicates the k-mer set of (*H*_1_, *H*_2_), ***RS*** indicates the read set, *count*(*kmer*_*i*_, (*H*_1_, *H*_2_)) and *count*(*kmer*_*i*_, ***RS***) indicate the counts of a specific k-mer (*kmer*_*i*_) on (*H*_1_, *H*_2_) and ***RS***, respectively. The haplotype-pair having highest likelihood is selected and realigned to reference (with the graph walking information) to infer SV breakpoints and make the call.

### SV genotyping and quality control

gcSV calculate the posterior probability *P*(*G*|*D*) of a specific genotype *G* given an observed set of related alignment *D* for genotyping. Assuming the list of the candidate alleles as *A* = {*r, x*_1_, … …, *x*_*n*_}, where *r* refers to reference allele and *x*_*i*_, *i* = 1, …, *n* refers to SV-alleles, there are |*A*|^2^ types of candidate genotypes, i.e., *G* = {*rr, rx*_1_, *rx*_2_, *rx*_3_, … … ., *x*_*n*−1_*x*_*n*_} and *P*(*G*|*D*) is formed as following:

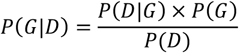

Further, the equation is reduced as *P*(*D*|*G*) × *P*(*G*) since *D* has been observed. The prior *P*(*G*) is set as:

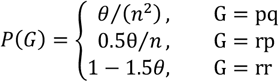

where *θ* is a constant (default value: 10^−5^) and *n* is the number of non-reference alleles.

Further, each alignment is considered as an independent event and the conditional probability *P*(*D*|*G*) is written as the product of the probabilities of the events:

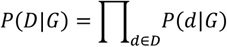

where *P*(*d*|*G*) is the probability of observing a specific abnormal alignment *d* given the genotype *G*,

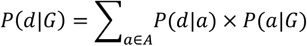

Similarly, *P*(*d*|*a*) represents the probability of observing a specific alignment *d* given a particular haplotype *a* . *P*(*a*|*G*) denotes the probability of observing a specific haplotype a under the condition of a given genotype *G*, which is taken from {0,025,120}.

gcSV applies DRP and the penalty scores of realigned reads to compute sf *P*(*d*|*a*). gcSV calculates the distribution of insert sizes for all paired-end reads and normalizes the results to obtain the probability density function *p* = *f*(*x*)for the insert size. For each DRP, gcSV mark the insert size as the distance of two breakpoint (termed as *isize*) and get the probability *P*(*d*|*a*) = *f*(*isize*). After realigned reads to different candidate haplotypes, gcSV recalculates the alignment scores by setting the penalty scores as 0, 3, 4, 1 for match, mismatch, gap open and gap extension, respectively. The penalty score has an upper limit (default is 5) and the total penalty score for realignment is denoted as *S, P*(*d*|*a*) is set as 10^−*S*^ consequently.

Under diploid assumption, there are three possible genotypes, that is homozygous reference (i.e., no variation), heterozygous, and homozygous variant (*G* = {*rr, rx, xx*}). The genotype *G* maximizing *P*(*G*|*D*) is selected and the quality score is computed by the following equation,

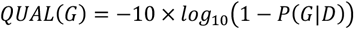

The results corresponding to homozygous reference genotypes are directly discarded. Additionally, results with quality scores below the threshold (default is 20) are labeled as *LOW_QUAL*, while others are labeled as *PASS*

### Benchmark on real sequencing datasets

Three real sequencing datasets of HG002 sample were downloaded by following previous studies where one is a 30x PacBio HiFi sequencing dataset and the other two are 60x and 35x Illumina short read sequencing datasets, respectively (refer to Supplementary Table 1 for the accessions). Moreover, the 30x LRS dataset was down-sampled as 1-10x, 15x, 20x and 25x and the 60x SRS dataset was down-sampled as 15x and 30x for more comprehensive benchmark, respectively. The down-sampling operations were conducted by Samtools^64^. For all the datasets, the reads were aligned to human reference genome (h37d5) beforehand by pbmm2 (long reads) and BWA-MEM (short reads), respectively, and the sorted BAM files were used as inputs.

The NIST-Q100 callset was downloaded from the official site of Genome in a Bottle Consortium and used as ground truth. The assessment of the yield metrics (F1-score, sensitivity, precision, GT concordance) were done by Truvari (run in ‘single’ mode). The pctseq (-p) parameter was set to 0. Only the variant calls marked as “pass” were considered and duplication calls were treated as insertions by setting dup-to-ins. The pctsize (-P) parameter was set as its default value (0.7), i.e., a called SV is considered as true positive if there is a SV record in the ground truth having 70% overlap with it. This is except for the strict SV matching assessment which the parameters were respectively set as 0.9, 0.95 and 1 to investigate the ability of the SV callers for the requirement of more accurate SV reconstruction.

For LRS benchmark, we implemented gcSV, Sniffles, cuteSV and SVDSS on the 1-10x, 15x, 20x, 25x and 30x PacBio HiFi dataset. It is worthnoting that we adjusted the parameters settings of Sniffles and cuteSVs for relatively low coverage datasets (Supplementary Table2) to maximize their sensitivity. For gcSV and SVDSS, the default settings were used for all the datasets. The results on the whole genome (defined NIST’s truth high confidence BED file) were assessed at first. Further, we separately investigate the yields of the callers on various types and sizes. The insertion and deletion types of SV calls were extracted from the callset (by the SVTYPE information recorded in vcf files) and assessed. To evaluate the various sized SVs, Truvari was implemented on the callsets with additional settings for sizemax and sizemin parameters correspondingly. We also used the annotations provided by RepeatMasker^65^ and CMRG^51^ to assess the yields of the callers on various categories of repeat regions (SINE, LINE, SimpleRepeat, LTR, DNA, Satellite, LowComplex, Retroposon and RNA, respectively) and difficult SVs. To compare the various callsets, the true positive calls of the various approaches were extracted and merged by using SURVIVOR^66^ (parameter setting: 10 1 1 1 0 50). Further, the calls were compared by implementing bcftools^67^ on the merged SV callset.

For hybrid sequencing benchmark, we implemented gcSV on 12 datasets which were generated by the combinations of 1x, 2x, 3x, 4x, 5x, 10x LRS data with 30x and 60x SRS data. The yields on whole genome, various SV types and sizes, various repetitive regions were assessed in a similar approach to that of LRS benchmark. Moreover, to assess the use of short reads in hybrid-based SV calling, we compared the corresponding callsets of 1x, 2x, 4x and 10x LRS data w/o 60x SRS data. The comparison was also conducted similarly to that of LRS benchmark. It is also worthnoting that, the source code of Blend-seq^50^ is still not fully open at the time of the submission of this manuscript, so that we directly quote its statistics on NIST-Q100 callset presented in its pre-print to compare gcSV with it.

For SRS benchmark, we directly implemented gcSV, Manta, Delly and Lumpy on the 15x, 30x and 60x SRS datasets. As DRAGEN and GATK-SV are also not fully open-source and easy to implement, we implemented gcSV on the 35x SRS dataset used in the DRAGEN study^30^ to make the benchmark comparable. Truvari was then implemented on the produced callsets of various genomic regions (whole genome and HARD and easy regions) to assess the yields of the callers, where the HARD and EASY regions for short reads were employed by referring to the previous study^50^. The yields on various SV types and sizes were assessed similarly. The whole SV sets of the ground truth, DRAGEN and gcSV (w/o long reads) were compared by using SURVIVOR and bcftools to plot the Venn diagram. Moreover, the ground truth, 1KGP Phase4 callset and HGSVC callset were merged by a different parameter setting (500 1 1 1 0 50) of SURVIVOR to compromise the short and long read-based SV calls. Their Venn diagram were then plotting by using bcftools.

We downloaded the read alignment files (in CRAM format) of all the 3,202 samples being sequenced in 1KGP Phase4 (Supplementary Table 1). gcSV was then implemented on each of the CRAM files to conduct SV calling (refer to Supplementary Notes for command lines). Further, the produced individual callsets (in VCF format) were merged by SURVIVOR (only the non-BND SVs passed quality control were considered). In the merged callset, the quality score of each SV was re-estimated by summing up the corresponding quality scores of all the supporting samples. Moreover, the SVs having <50 quality score were removed after the re-estimation. For each of the remaining SVs, the allele counts (ACs) and allele numbers (ANs) were then calculated, and it is also annotated by AnnotSV (parameters: -genomeBuild GRCh38 -SVminSize 30; Supplementary Note). The callset was also compared to the official callset of 1KGP Phase4 by using Truvari Bench (parameters: --passonly -p 0 --dup-to-in --pick single) to distinguish the novel SVs detected by gcSV.

Also refer to Supplementary Notes for the command lines used in the benchmark.

## Data availability

The GIAB HG002 Illumina and PacBio HiFi data, the high-coverage WGS data from the 1KGP cohort, GIAB benchmark sets, and relevant genomic regions are listed in Supplementary Table 1. All the version and download URL of tools used in this study are listed in Supplementary Table 12. Individual VCF files across the samples, along with the population VCF file obtained by reanalyzing the 1KGP4 high-coverage cohort (3,202 samples) for gcSV, are publicly available at https://github.com/hitbc/gcSV_call_results.

## Code availability

The gcSV program is written with c++ language, the source code (v1.0.0) and auxiliary scripts used to generate all the results in this study are provided at GitHub (https://github.com/hitbc/gcSV), which is available under GNU General Public License v3.0.

## Acknowledgements

This work was supported by the National Natural Science Foundation of China (No. 62331012, No. 62172125 and No. 62402140) and the Ministry of Science and Technology of China (2024YFF1206300, 2024YFF1206200, 2023YFF1205600 and 2017YFC0907500).

## Author contributions

Y. W. designed and supervised research; G. L. and B. L. developed the algorithm and software; Y. L., B. L. and L. Q. contributed to data interpretation; G. L. performed the algorithm benchmarking on the real data; G. L., Y. L. and Q. L. contributed to the re-analysis of 1KGP4 high-coverage cohort. G. L., Y. L., B. L., and L. Q. wrote the paper with input from all other authors. All authors read and approved the final manuscript.

## Competing interests

The authors declare no competing interests.

## Notes

### Competing Interest Statement

The authors have declared no competing interest.

