## Supplementary Material for "gcSV: a unified framework for comprehensive structural variant detection"

Supplementary Figure 1. The distributions of SV signatures in various repetitiveness context 3
Supplementary Figure 2. A schematic illustration on refined two-phase SV signature clustering 6
Supplementary Figure 3. A schematic illustration on generative SV reconstruction and genotyping 7
Supplementary Figure 4. The ability for long read-based SV calling (without genotyping) 9
Supplementary Figure 5. Examples of gcSV to detect complex SVs 10
Supplementary Figure 6. Examples of highly matched SV events with divergent representations 11
Supplementary Figure 7. A schematic illustration on the failure of SV allele reconstruction and genotyping with insufficient long reads 13
Supplementary Figure 8. Examples of gcSV to detect large insertions using short reads 14
Supplementary Figure 9. Examples of large deletions called by DRAGEN but not gcSV 15
Supplementary Figure 10. Allele counts of population-specific SVs in gcSV-callset and 1KGP-callset 16
Supplementary Figure 11. Tandem repeat-associated SVs in the gcSV-callset 17
Supplementary Table 1. The availability of the datasets used for benchmark 18
Supplementary Table 2. The yields of various approaches using long read sequencing data 19
Supplementary Table 3. The yields of various approaches using hybrid sequencing data 21
Supplementary Table 4. The yields of various approaches using short read sequencing data 22
Supplementary Table 5. The yields of various approaches for SV sizes and types (long reads) 23
Supplementary Table 6. The yields of various approaches in various genomic regions (long reads) 25
Supplementary Table 7. The yields of gcSV for various SV sizes and types (hybrid sequencing) 27
Supplementary Table 8. The yields of gcSV in various genomic regions (hybrid sequencing) 32
Supplementary Table 9. The yields of various approaches in the EASY/HARD regions for short reads 35
Supplementary Table 10. The yields of various approaches for various SV sizes and types (short reads) 37
Supplementary Table 11. OMIM, dbVar, ClinVar annotated pathogenic SVs discovered by gcSV in 1KGP dataset 40
Supplementary Table 12. All the resources used in this study 41
Supplementary Notes 42

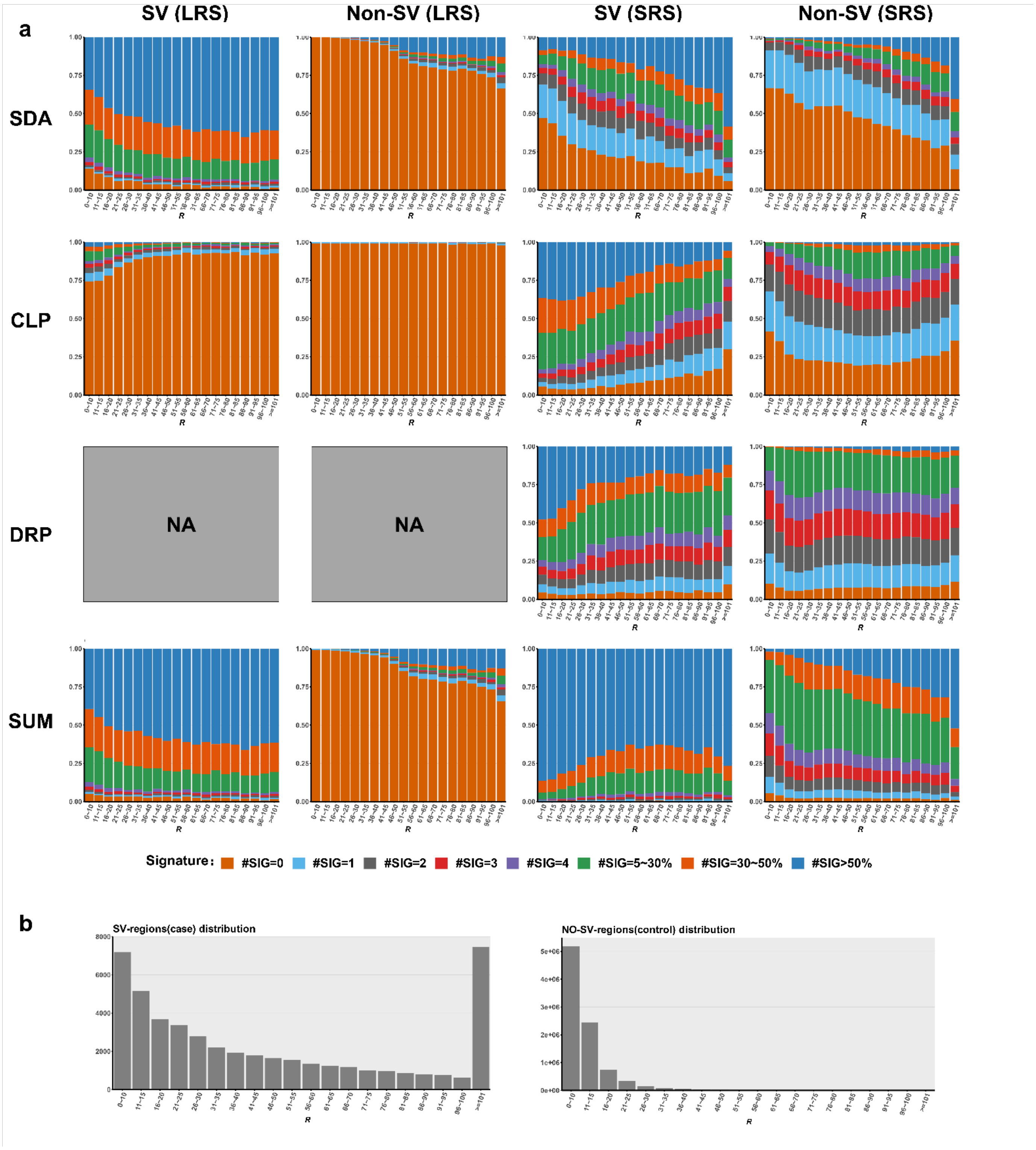

**Supplementary Figure 1. The distributions of SV signatures in various repetitiveness context**

**a,** The distributions of various kinds of signatures exhibited by SRS and LRS (from HPRC datasets). The four rows indicate the distributions of SDA, CLP, DRP (only available for SRS) signatures and their sums, respectively, and the four columns indicate the LRS and SRS signatures in SV- and non-SV windows, respectively. Each of the distribution is shown as a histogram (in the form of stacked bar plots). Each bar is corresponding to a specific range of local repetitiveness ($R$, 10 degrees per bar), and the colored blocks indicate the proportions of the windows in the context of $R$ that have specific numbers of signature reads (i.e., #SIG in figure, indicating the windows having 0, 1, 2, 3, 4, 5 reads in absolute terms and 5-30%, 30-50%, >50% of total number of reads, respectively). **b,** The histogram of the SV- and non-SV windows of human reference genome corresponding various $R$ ranges (shown in absolute terms of window numbers).

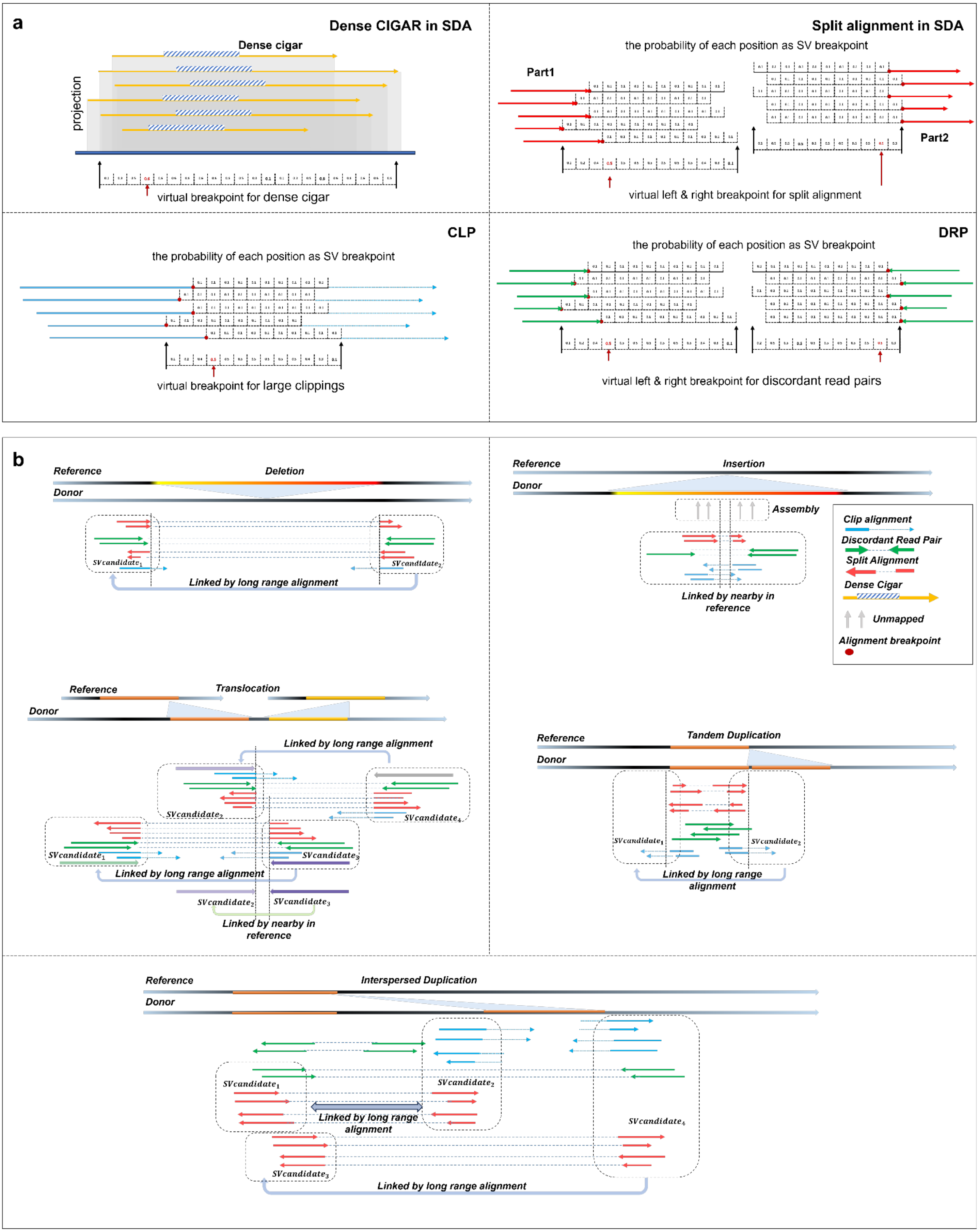
a
Dense CIGAR in SDA
Dense cigar
projection
virtual breakpoint for dense cigar
Split alignment in SDA
the probability of each position as SV breakpoint
Part1
Part2
virtual left & right breakpoint for split alignment
CLP
the probability of each position as SV breakpoint
virtual breakpoint for large clippings
DRP
the probability of each position as SV breakpoint
virtual left & right breakpoint for discordant read pairs
b
Reference
Deletion
Donor
Linked by long range alignment
Reference
Insertion
Donor
Assembly
Linked by nearby in reference
Clip alignment
Discordant Read Pair
Split Alignment
Dense Cigar
Unmapped
Alignment breakpoint
Reference
Translocation
Donor
Linked by long range alignment
Linked by long range alignment
Linked by nearby in reference
Reference
Tandem Duplication
Donor
Linked by long range alignment
Reference
Interspersed Duplication
Donor
Linked by long range alignment
Linked by long range alignment

**Supplementary Figure 2. A schematic illustration on refined two-phase SV signature clustering**

**a,** Phase1: virtual breakpoint-based read clustering and refinement. gcSV estimates the virtual breakpoint implied by SDA, CLIP, DRP read clusters in three different approaches. For SDA signature in dense cigar form (upper left panel, the dashed blocks indicate the dense cigars), gcSV projects all the bases of the reads to the reference and define the distribution of SV breakpoint on the positions having at least one base being mapped to. For each of the position, a probability is assigned as the number of bases being mapped (normalized by the total number of bases), further, gcSV separately tests each of the reads where the p-value is determined as the probability at the mapped centroid of the read in reference. For SDA signature in split alignment form (upper right panel), gcSV separately handles the multiple parts of the alignments. For each part, gcSV extends an $L$ bp (default value: 10 bp) block from the split point, projects the block to reference and assigns a weight (default value: 0.1) to each of the projected positions. Further, the null distribution is defined on all the reference positions having at least one projected base, and for each position the probability is set as the sum weights of all the bases being projected there. Further, gcSV tests each of the reads using this distribution. For CLP signature (lower left panel), gcSV extends an $L$ bp (default value: 10 bp) block from the clipping point and implements base projection and weighting in a similar way to that of split alignment. Further, the null distribution is also defined on all the reference positions being projected for read testing. For DRP signature (lower right panel), gcSV also handles the involved read pairs similarly. The difference is that the extended blocks are generated from the downstream and upstream alignment endpoints of the two paired reads, respectively. With the extended blocks, the null distribution is also composed based on the sum weights of the read projection positions. **b,** Phase2: the connection of refined clusters. gcSV greedily connects the read clusters in nearby regions and/or being linked to combine the read clusters belonging to the same SV event but having various types of signatures or being interspersed caused by multiple breakpoints and/or genomic repeats. For a deletion event (upper left panel), gcSV directly integrates the nearby clusters of SDA, CLP and DRP. Moreover, the DRP clusters can be further linked by read pairing information. For an insertion event (upper right panel), nearby CLP, SDA and DRP clusters are directly integrated similarly. Since insertion events can shorten the junctions of paired reads, in most cases the clusters can be comprehensively collected in a local region without regarding to the distant linking information. For a translocation event (middle left panel), it typically has two breakpoints and four involving clusters, nearby clusters were linked initially and then the long-range clusters are then clustered. For a tandem duplication event (middle right panel), CLP, SDA and DRP produce two breakpoints located around the boundaries of the duplication, separately. At each boundary, the nearby breakpoints from each category are first combined, and then the long-range clusters are straightforwardly merged. For an interspersed duplication event (lower panel), CLP and SDA produce four involving clusters, the split alignment signature support long-range information to combine all the clusters. Also refer to Fig. 1b for the illustration for inversion events.

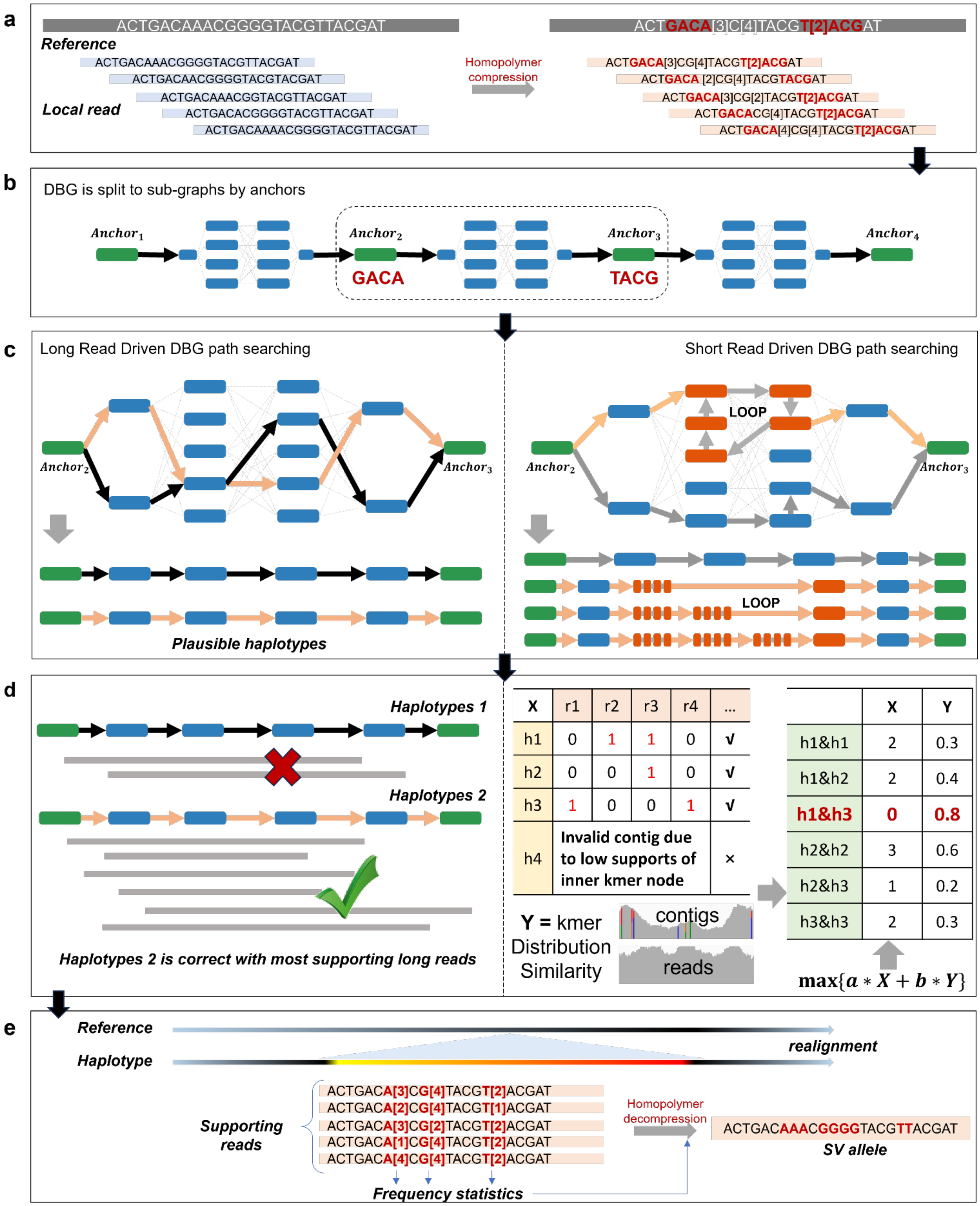

**Supplementary Figure 3. A schematic illustration on generative SV reconstruction and genotyping**

gcSV performs assembly context-driven hypothesis test approach to reconstruct SV alleles as following steps. **a,** Kmer extraction and indexing. gcSV index the local reference sequence and the clustered reads by homopolymer-compressed (HPC) *k*-mers. **b,** Graph construction. gcSV builds a de Bruijn graph (DBG) using all the HPC k-mers (with kmer counting and filtration) and split the DBG into multiple subgraphs by two unique k-mers (anchor) as the source- and sink nodes. **c,** Candidate allele generation. gcSV implements depth-first search to generate one or more valid paths that each of them starts and ends at one of the source- and the sink nodes, respectively. If long reads are available (left panel), the search is under the guidance of reads, i.e., the nodes of the path should be successively matched to the kmers in the reads. If only short reads are available (right panel), the search is guided by coverage-based heuristics that each of the kmers in the graph are assigned an upper limit of visiting times. During the searching process, the visiting times of the nodes are recorded and gcSV only allows to add the nodes having not been visited more than the upper limit to the generating path. With such heuristics, multiple paths having various copy numbers of local repeats can be produced as candidates, especially in tandem repeat regions. **d,** Allele testing. Any pair of the paths (alleles) is composed as a hypothesis under diploid assumption, and the k-mer-based statistical approach is used. gcSV filters out the candidates if one or more of their internal nodes cannot be covered (i.e., explained) by the kmer set of the reads. Further, the likelihoods of the hypotheses are separately computed (in a table form) for the selection of the most likely allele pair. **e,** SV calling. gcSV implements homopolymer decompression for the selected contigs and realigns them to the reference to infer the type and breakpoints of the SV events and reports.

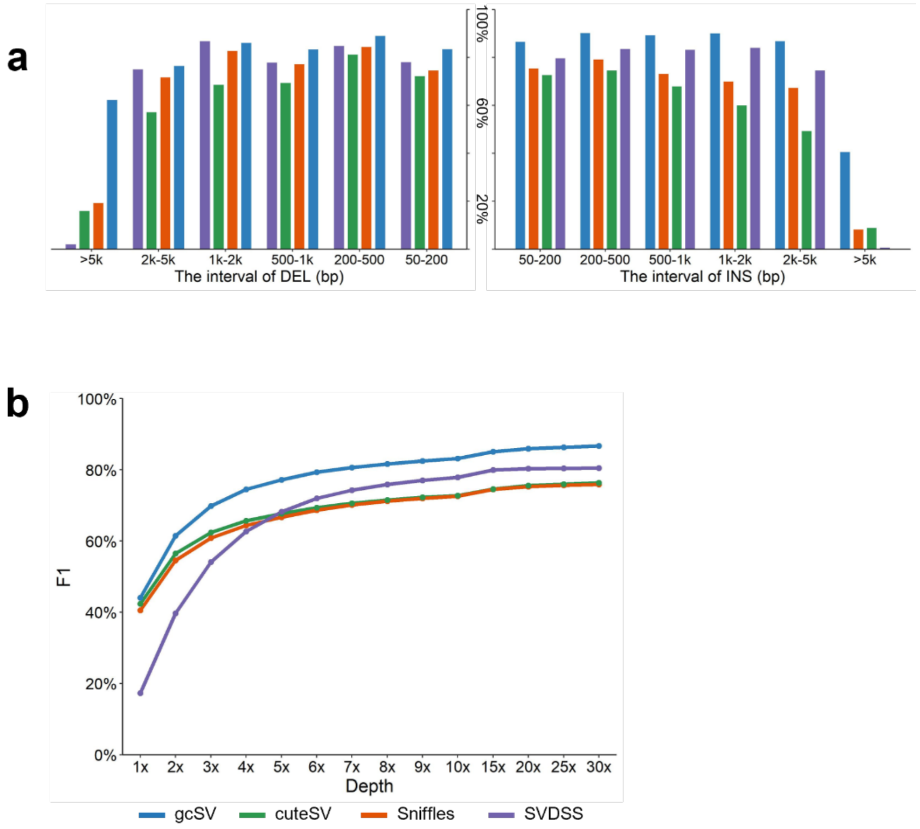

**Supplementary Figure 4. The ability for long read-based SV calling (without genotyping)**

**a,** The F1-scores (without genotyping) of the long read-based callers (gcSV, Sniffles, cuteSV and SVDSS) for various types and sizes of SVs. The horizontal axis indicates the types (left: deletions, right: insertions) and sizes (in the ranges of 50-200bp, 200-500bp, 500-1kbp, 1k-2kbp, 2k-5kbp and >5kbp, respectively). The results of gcSV, Sniffles, cuteSV and SVDSS are shown in various colored bars. The dark and light colors indicate their F1-scores with and without genotyping. **b,** The F1-scores (without genotyping) of the callers for various read coverages (1-30x).

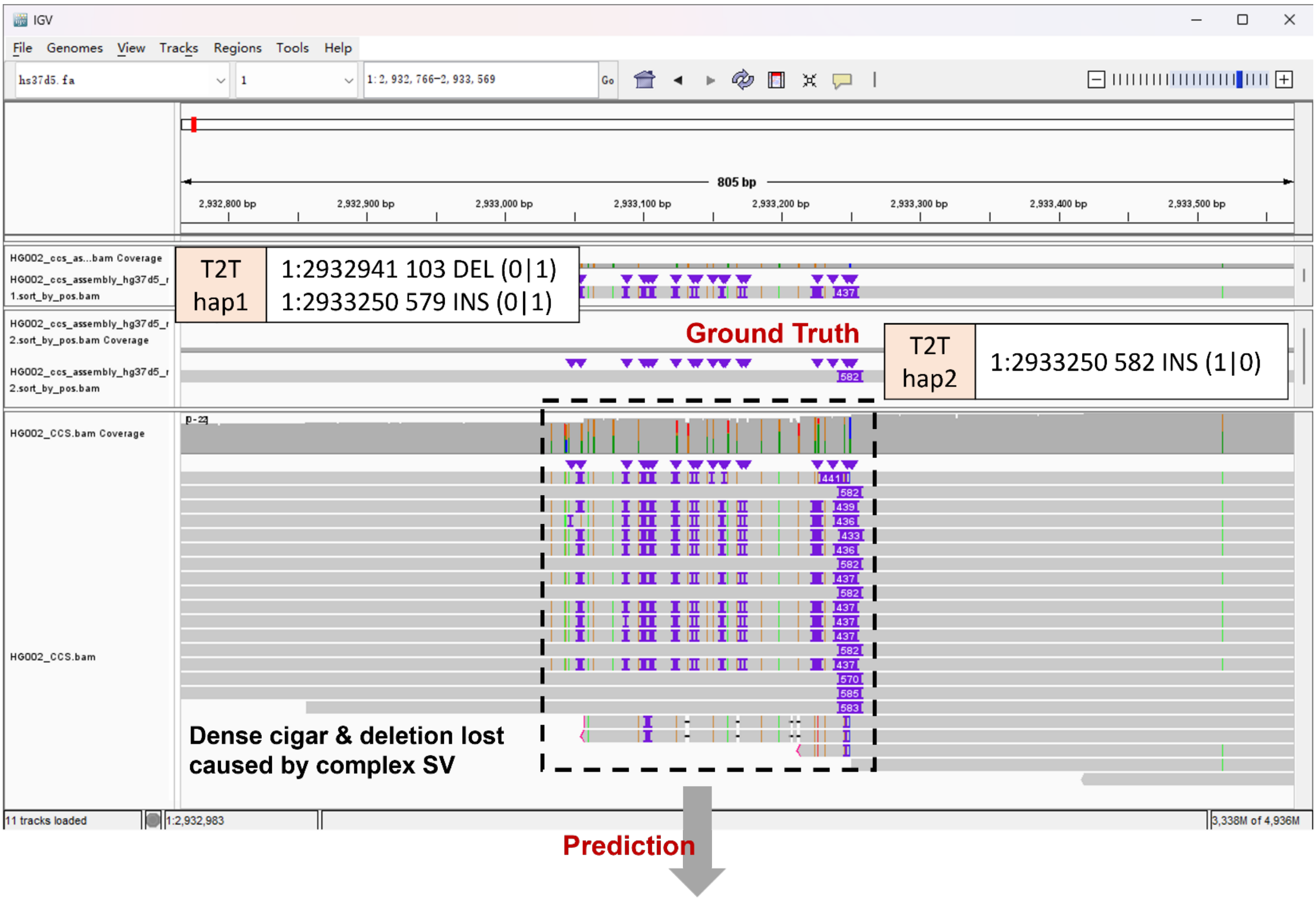

| Caller | Alleles in Hap1 | Alleles in Hap2 | SV allele match | Zygosity match | Sequence match |
|---|---|---|---|---|---|
| gcSV | 1:2932941 103 DEL (0\|1)<br>1:2933250 579 INS (0\|1) | 1:2933250 582 INS (1\|0) | √ | √ | √ |
| Sniffles | 1:2933251 582 INS (1/1) | | × | × | × |
| cuteSV | 1:2933250 496 INS (1/1) | | × | × | × |
| SVDSS | 1:2933247 437 INS (1/1)<br>1:2933250 582 INS (1/1) | | √ | × | × |

**Supplementary Figure 5. Examples of gcSV to detect complex SVs**

The Integrated Genomics Viewer (IGV) snapshot illustrates the detection of complex structural variations (SVs) in a local region (hs37d5, Chr1:2,933,000-2,933,400). According to the ground truth, this region is a heterozygous SV site having two alternative alleles, i.e., a combination of 103 bp deletion and 579 bp insertion (first allele) and a 582 bp insertion (second allele). In the 30x PacBio HiFi dataset, plenty of reads have dense CIGARs there (i.e., SDA signatures), however, the signatures are in quite complex forms, more importantly, no significant deletion signatures are provided by the read aligner, due to the crosstalk of local repetitiveness and the multiple SV events. Alignment-based callers (Sniffles and cuteSV) detected only one homozygous insertion, reporting sizes of 582 bp and 496 bp, respectively, since they heavily rely on CIGAR string parsing. SVDSS detects two homozygous insertions (437 bp and 282 bp) by its local assembly approach, however, this result is still not enough to reconstruct the SV alleles fully matched to the ground truth. Moreover, it also fails at genotyping which could be important to downstream analysis. Using the context-driven local assembly and genotyping, the output of gcSV is exactly matched to ground truth, for both of alleles and genotypes.

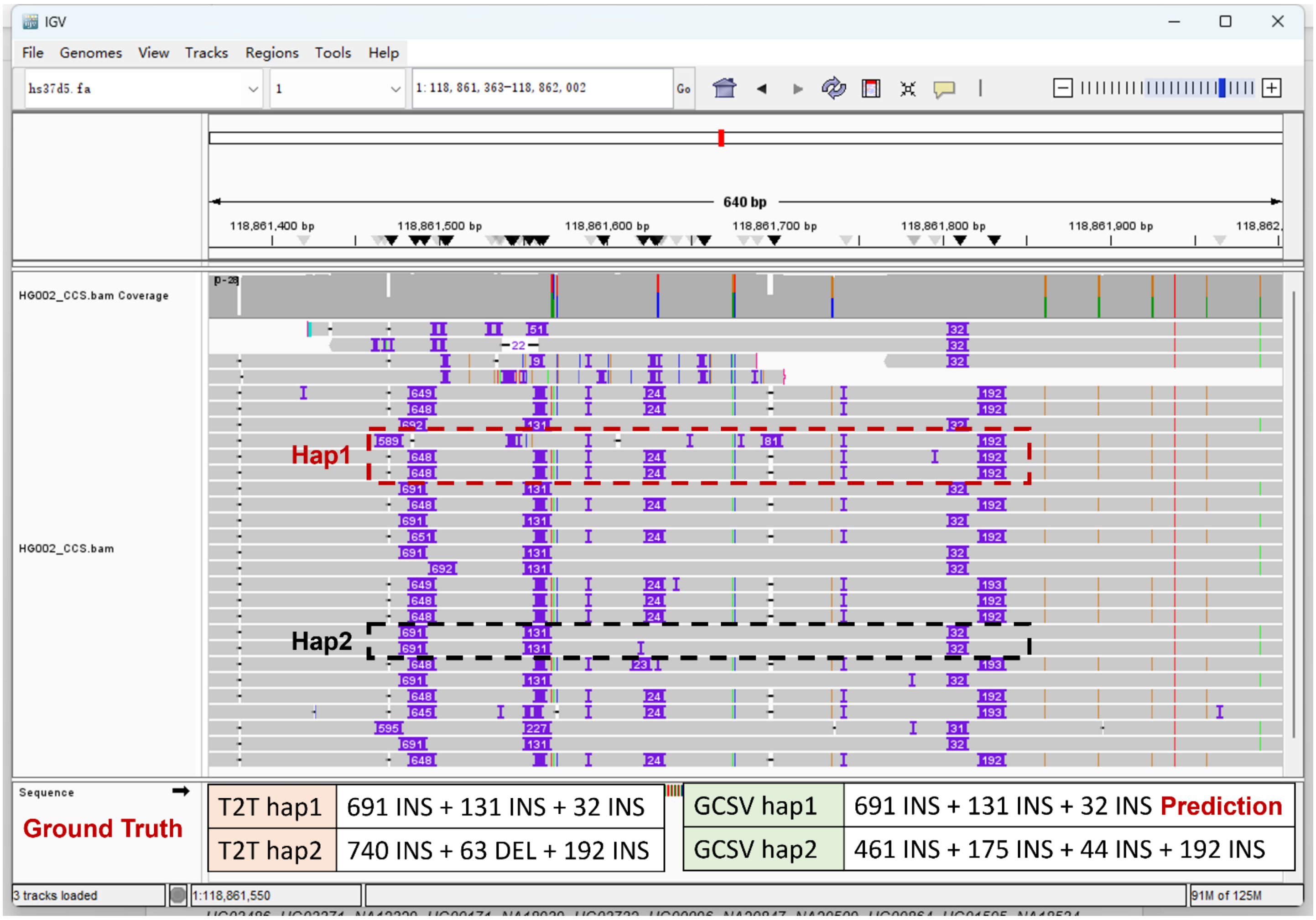

**Supplementary Figure 6. Examples of highly matched SV events with divergent representations**

The IGV snapshot for several called SVs in a local genome region (hs37d5, Chr1:118,861,163- Chr1:118,862,002). The HG002 ground truth suggests three insertions in one haplotype, i.e., 691 bp, 131 bp and 32 bp, while gcSV detects the same three insertions with identical insertion size (i.e., identical SV representations). However, in the other haplotype, the ground truth suggests a combination of a 740 bp insertion, a 63 bp deletion and a 192 bp insertion. gcSV reports four SV events as 416 bp insertion, 175 bp insertion, 44 bp insertion and 192 bp insertion, i.e., the former three are different from the ground truth linguistically. However, by putting the events back to the reference, the generated alleles of the donor genome by the ground truth and gcSV calls can be exactly matched to each other, indicating that the differences are caused by the various SV representations made by allele realignment, but not real false positive/negative calls in practice.

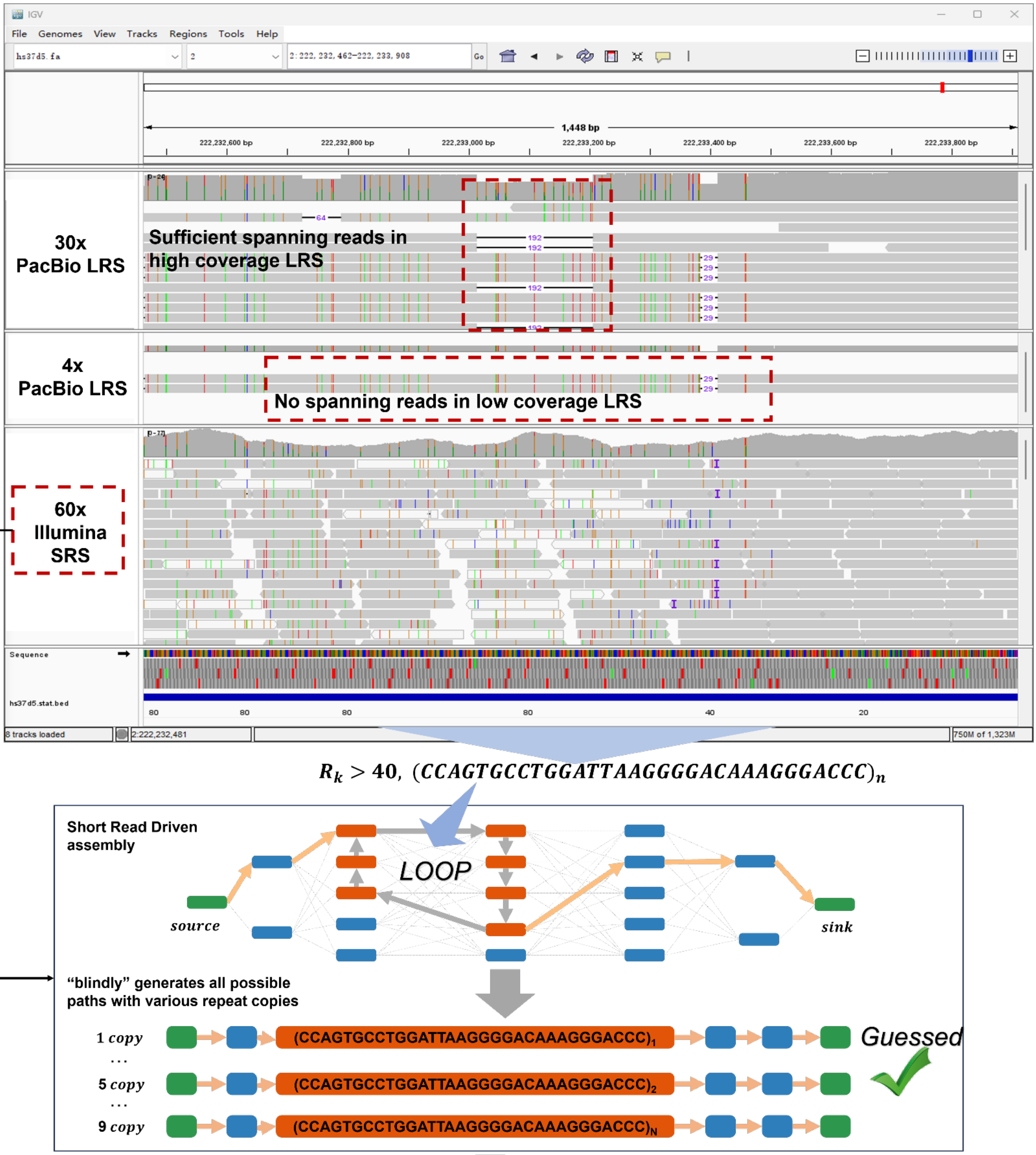

| | Position | Variant type | Deleted copies guessed (repeat length) |
|---|---|---|---|
| Ground truth | 2:222233014-222233206 | 192-bp deletion (VNTR) | 6 (32) |
| gcSV (60x SR only) | 2:222233014-222233142 | 128-bp deletion (VNTR) | 4 (32) × |
| gcSV (60x SR+ 4x LR) | 2:222233014-222233142 | 128-bp deletion (VNTR) | 4 (32) × |
| | **Without insufficient long reads for the SR only and low coverage LR to confirm the repeat and fail to reconstruct SV allele** | | |
| gcSV (30x LR only) | 2:222233014-222233206 | 192-bp deletion (VNTR) | 6 (32) √ |

**Supplementary Figure 7. A schematic illustration on the failure of SV allele reconstruction and genotyping with insufficient long reads**

The IGV snapshot for a VNTR in a highly repetitive region (hs37d5, Chr2:222,232,600-222,233,600, $R_k > 40$) highlights the challenges of insufficient long reads during SV allele reconstruction. The reference allele consists of 9 tandem repeat units ("CCAGTGCCTGGATTAAGGGGACAAAGGGACCC"), and the event is of 3 units according to the ground truth, resulting in a 192 bp deletion. In the 30x PacBio HiFi dataset (the upper part of the snapshot), a number of reads can span the region and provides the information of 192 bp deletions in their CIGARS, which helps to directly call the SV out. However, with the 60x Illumina dataset (the lower part of the snapshot), none of the reads can span the SV site, thus the SV allele reconstruction is quite hard under the effect of high local repetitiveness. gcSV tries to solely use heuristic graph walking (lower part of the figure) and and nine candidate alleles are generated (respectively 1-9 copies). Due to lack of long reads, gcSV uses the k-mer frequency-based approach to test each of them, however, an incorrect call (5 copies, 128bp deletion) is made under the effect of local repetitiveness and the divergence of read coverage. In the hybrid sequencing scenario (4x long reads, the middle part of the snapshot), the SV is expected to be correctly detected since only a few (even one) long reads can provide sufficient power for the allele testing. However, due to the divergence of read coverage, there is unexpectedly no long read in the region, thus leads to an unusual failure for gcSV. It is also worthnoting that DRAGEN does not give any call for this region.

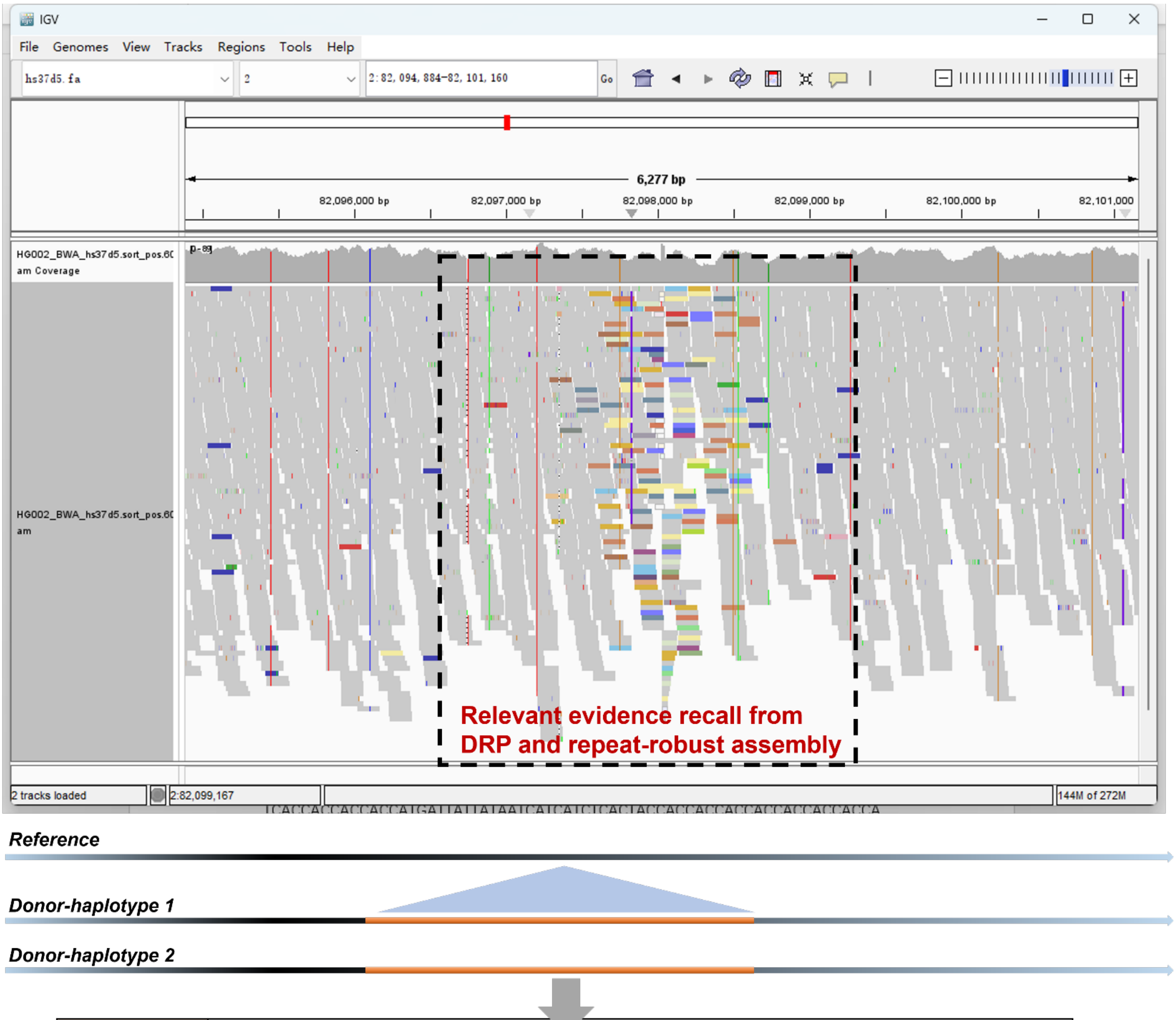

| | | | | | | |
|---|---|---|---|---|---|---|
| Position | 2:82098027 | | | | | |
| Type | 6056-bp large insertion (L1) | | | | | |
| Zygosity | Homozygosity | | | | | |
| Caller | gcSV | DRAGEN | GATK-SV | Manta | Delly | Lumpy |
| | √ | × | × | × | × | × |

**Supplementary Figure 8. Examples of gcSV to detect large insertions using short reads**

The IGV snapshot for a large insertion event at Chr2:82,098,027 which is a typical mobile element (L1) insertion. Using short reads, the event can only be detected by gcSV during the benchmark. gcSV achieves this goal by the refined read clustering which collects plenty of spanning reads by their DRP signatures (the colored ones in the figure). Moreover, the tailored local assembly approach also helps to precisely reconstruct the inserted L1 sequence (6056 bp).

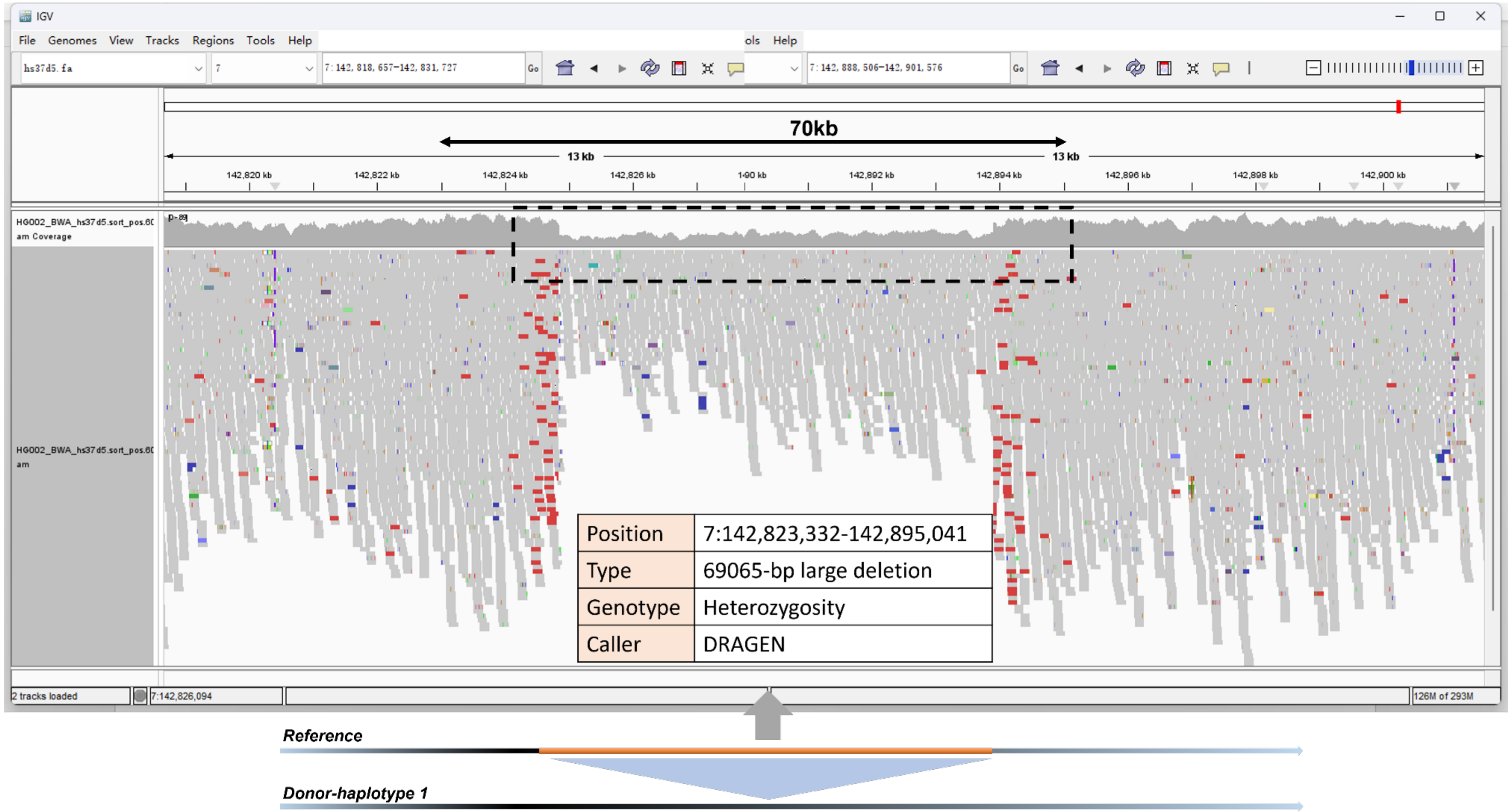

**Supplementary Figure 9. Examples of large deletions called by DRAGEN but not gcSV**

The IGV snapshot for a large deletion at Chr7:142,823,332-142,895,041 which is detected by DRAGEN only. This deletion is in highly repetitive regions and highly challenging to SRS-based SV detection without any specific prior such as pan-reference. Considering that the event is also not detected by GATK-SV (using Manta, Lumpy and many other specifically designed tools), we speculated that DRAGEN detect this event by the joint use of its unpublished pan-genome reference and certain components specifically designed for such events as well.

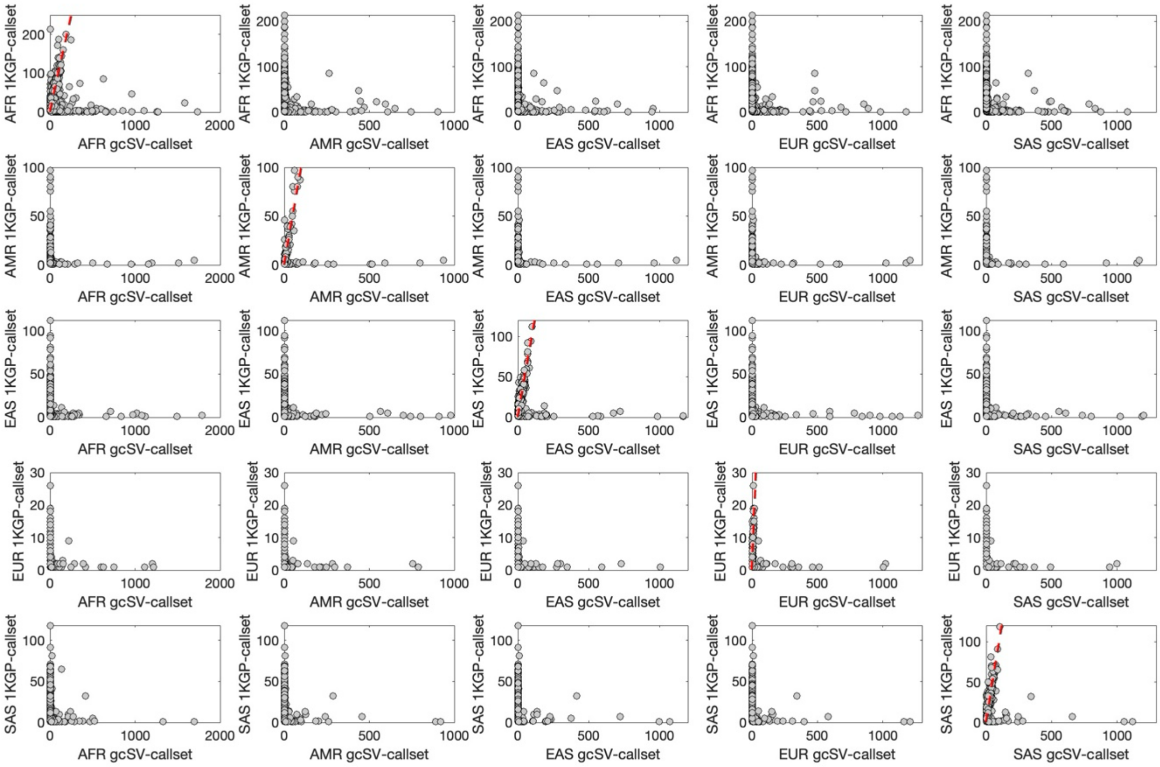

**Supplementary Figure 10. Allele counts of population-specific SVs in gcSV-callset and 1KGP-callset**

Only SVs regarded unique to the superpopulation indicated on the y-axis in the 1KGP-callset were plotted. The allele count of these SVs in the same superpopulation (diagonal) or in another superpopulation, in the gcSV-callset, is shown on the x-axis. The allele count of these SVs in the 1KGP-callset is shown on the y-axis. In the diagonal subplots, the red dashed lines mark the 1:1 ratio. In the main text, SVs with strongly elevated allele frequencies are those in the diagonal subplots whose allele frequencies in gcSV-callset became at least 2-fold of their allele frequencies in 1KGP-callset, in the same superpopulation.

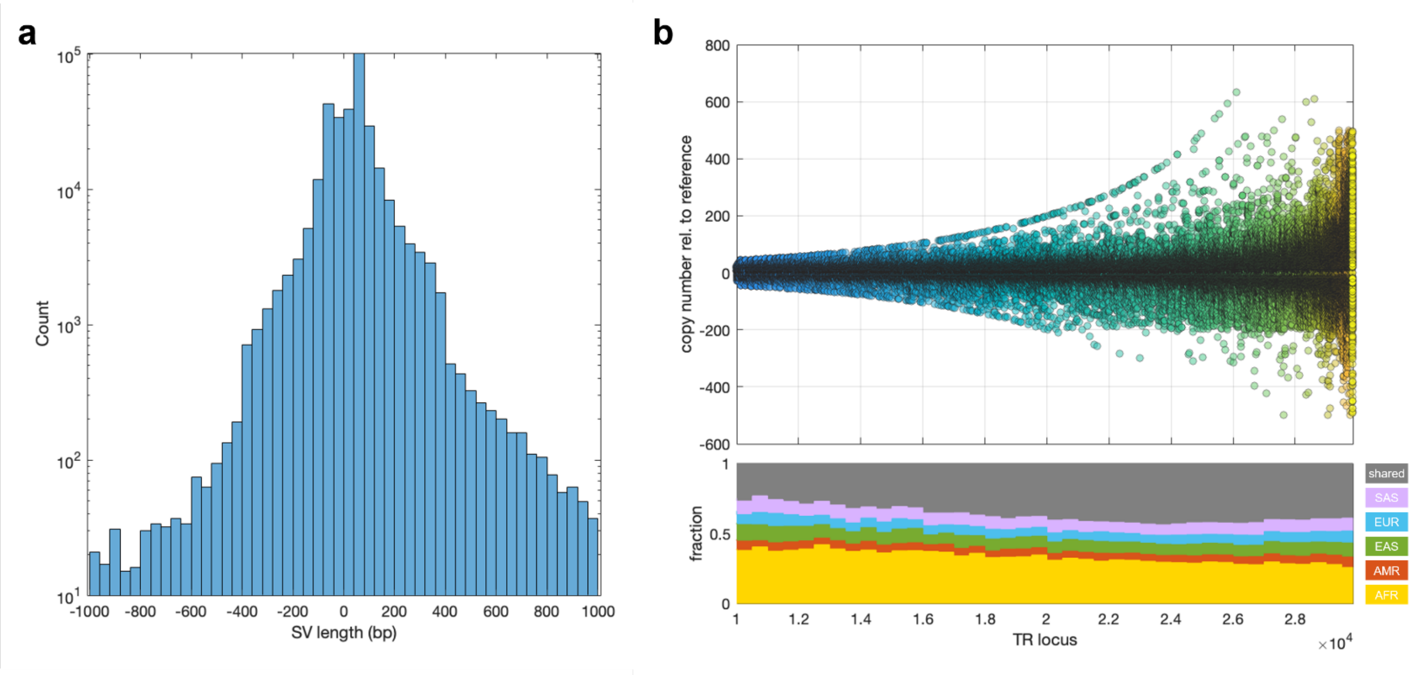

**Supplementary Figure 11. Tandem repeat-associated SVs in the gcSV-callset**

**a,** The length distribution of tandem repeat-associated SVs in gcSV-callset, shown on the logarithmic scale. **b,** Copy number variations (upper panel) and the fractions of population unique SVs (lower panel) at each tandem repeat locus. The tandem repeat loci are sorted on the x-axis by increasing variance.

**Supplementary Table 1. The availability of the datasets used for benchmark**

| No. | Dataset | Type | Accession |
|---|---|---|---|
| 1 | *HG002 LRS 30X* | PacBio HiFi long reads | https://ftp-trace.ncbi.nlm.nih.gov/ReferenceSamples/giab/data/AshkenazimTrio/HG002_NA24385_son/PacBio_CCS_15kb/alignment/HG002.Sequel.15kb.pbmm2.hs37d5.whatshap.haplotag.RTG.10x.trio.bam |
| 2 | *HG002 SRS 60X* | Illumina short reads | https://ftp-trace.ncbi.nlm.nih.gov/ReferenceSamples/giab/data/AshkenazimTrio/HG002_NA24385_son/NIST_HiSeq_HG002_Homogeneity-10953946/NHGRI_Illumina300X_AJtrio_novoalign_bams/HG002.hs37d5.60x.1.bam |
| 3 | *HG002 SRS 35X* | Illumina short reads | https://opendata.nist.gov/pdrsrv/mds2-2336/input_fastqs/HG002.novaseq.pcr-free.35x.R1.fastq.gz https://opendata.nist.gov/pdrsrv/mds2-2336/input_fastqs/HG002.novaseq.pcr-free.35x.R2.fastq.gz |
| 4 | *1KGP phase4 whole genome sequencing (3202 samples)* | Illumina short reads | https://ftp.1000genomes.ebi.ac.uk/vol1/ftp/data_collections/1000_genomes_project/data/ |
| 5 | *NIST HG002 Q100 benchmark (V1.0)* | Ground truth | https://ftp-trace.ncbi.nlm.nih.gov/ReferenceSamples/giab/data/AshkenazimTrio/analysis/NIST_HG002_DraftBenchmark_defrabbV0.015-20240215/ |
| 6 | *CMRG benchmark* | Genomic regions | https://ftp.ncbi.nlm.nih.gov/giab/ftp/release/AshkenazimTrio/HG002_NA24385_son/CMRG_v1.00/GRCh37/StructuralVariant/ |
| 7 | *RepeatMasker regions* | Genomic regions | https://hgdownload.soe.ucsc.edu/goldenPath/hg19/bigZips/hg19.fa.out.gz |
| 8 | *EASY and HARD regions for short read* | Genomic regions | https://ftp-trace.ncbi.nlm.nih.gov/ReferenceSamples/giab/release/genome-stratifications/v3.3/GRCh37@all/Union/ |

**Supplementary Table 2. The yields of various approaches using long read sequencing data [a]**

| Tool | Coverage | Precision [b] | Recall [b] | F1-Score-SV [b] | GT concordance [b] | Precision-GT [b] | Recall-GT [b] | F1-Score-GT [b] |
|---|---|---|---|---|---|---|---|---|
| **gcSV(D) [c]** | 30x | **91.30%** | **82.48%** | **86.67%** | **94.48%** | **86.27%** | **77.93%** | **81.89%** |
| **SVDSS(D)** | 30x | 85.10% | 76.29% | 80.46% | 53.98% | 45.93% | 41.18% | 43.43% |
| **Sniffles(M) [d]** | 30x | 90.83% | 65.17% | 75.89% | 81.01% | 73.58% | 52.79% | 61.47% |
| **cuteSV(M)** | 30x | 90.83% | 65.84% | 76.34% | 79.55% | 72.25% | 52.37% | 60.73% |
| **gcSV(D)** | 25x | **91.16%** | **82.00%** | **86.34%** | **94.23%** | **85.90%** | **77.27%** | **81.36%** |
| **SVDSS(D)** | 25x | 85.36% | 76.00% | 80.41% | 54.59% | 46.60% | 41.49% | 43.89% |
| **Sniffles(M)** | 25x | 90.82% | 64.86% | 75.67% | 80.85% | 73.42% | 52.44% | 61.18% |
| **cuteSV(M)** | 25x | 90.74% | 65.36% | 75.99% | 79.05% | 71.73% | 51.67% | 60.07% |
| **gcSV(D)** | 20x | **91.07%** | **81.35%** | **85.94%** | **93.87%** | **85.49%** | **76.36%** | **80.67%** |
| **SVDSS(D)** | 20x | 85.78% | 75.57% | 80.35% | 55.08% | 47.25% | 41.62% | 44.26% |
| **Sniffles(M)** | 20x | 90.66% | 64.33% | 75.26% | 80.53% | 73.01% | 51.81% | 60.61% |
| **cuteSV(M)** | 20x | 90.67% | 64.80% | 75.58% | 78.36% | 71.05% | 50.77% | 59.22% |
| **gcSV(D)** | 15x | **90.70%** | **80.13%** | **85.09%** | **92.99%** | **84.34%** | **74.51%** | **79.12%** |
| **SVDSS(D)** | 15x | 85.97% | 74.73% | 79.95% | 55.78% | 47.96% | 41.69% | 44.60% |
| **Sniffles(M)** | 15x | 90.62% | 63.26% | 74.51% | 79.66% | 72.19% | 50.39% | 59.35% |
| **cuteSV(M)** | 15x | 90.54% | 63.36% | 74.55% | 76.83% | 69.56% | 48.68% | 57.28% |
| **gcSV(D)** | 10x | 90.24% | **77.14%** | **83.18%** | **90.74%** | **81.88%** | **70.00%** | **75.48%** |
| **SVDSS(D)** | 10x | 85.71% | 71.29% | 77.84% | 57.19% | 49.02% | 40.77% | 44.52% |
| **Sniffles(M)** | 10x | **90.35%** | 60.70% | 72.61% | 77.33% | 69.86% | 46.94% | 56.15% |
| **cuteSV(M)** | 10x | 90.22% | 60.93% | 72.74% | 73.49% | 66.30% | 44.77% | 53.45% |
| **gcSV(D)** | 9x | 90.19% | **76.02%** | **82.50%** | **90.05%** | **81.21%** | **68.46%** | **74.29%** |
| **SVDSS(D)** | 9x | 85.87% | 69.90% | 77.06% | 57.83% | 49.66% | 40.42% | 44.57% |
| **Sniffles(M)** | 9x | **90.43%** | 59.81% | 72.00% | 76.44% | 69.13% | 45.72% | 55.04% |
| **cuteSV(M)** | 9x | 90.38% | 60.18% | 72.25% | 72.30% | 65.34% | 43.51% | 52.24% |
| **gcSV(D)** | 8x | 90.11% | **74.65%** | **81.66%** | **88.97%** | **80.18%** | **66.42%** | **72.65%** |
| **SVDSS(D)** | 8x | 85.89% | 67.96% | 75.88% | 58.21% | 50.00% | 39.56% | 44.17% |
| **Sniffles(M)** | 8x | **90.34%** | 58.77% | 71.21% | 75.36% | 68.08% | 44.28% | 53.66% |
| **cuteSV(M)** | 8x | 90.23% | 59.17% | 71.47% | 70.72% | 63.81% | 41.85% | 50.55% |
| **gcSV(D)** | 7x | 90.06% | **73.04%** | **80.66%** | **88.23%** | **79.46%** | **64.44%** | **71.16%** |
| **SVDSS(D)** | 7x | 86.07% | 65.34% | 74.29% | 58.56% | 50.40% | 38.26% | 43.50% |

| | | | | | | | | |
|---|---|---|---|---|---|---|---|---|
| **Sniffles(M)** | 7x | **90.32%** | 57.28% | 70.11% | 73.94% | 66.78% | 42.35% | 51.83% |
| **cuteSV(M)** | 7x | 89.99% | 58.02% | 70.56% | 68.98% | 62.08% | 40.03% | 48.67% |
| **gcSV(D)** | 6x | 90.09% | **70.89%** | **79.34%** | **87.06%** | **78.44%** | **61.72%** | **69.08%** |
| **SVDSS(D)** | 6x | 86.22% | 61.73% | 71.95% | 58.48% | 50.42% | 36.10% | 42.08% |
| **Sniffles(M)** | 6x | **90.12%** | 55.48% | 68.68% | 72.05% | 64.93% | 39.97% | 49.49% |
| **cuteSV(M)** | 6x | 89.76% | 56.49% | 69.34% | 67.02% | 60.16% | 37.86% | 46.47% |
| **gcSV(D)** | 5x | **90.22%** | **67.49%** | **77.22%** | **84.71%** | **76.42%** | **57.17%** | **65.41%** |
| **SVDSS(D)** | 5x | 86.62% | 56.22% | 68.19% | 58.11% | 50.33% | 32.67% | 39.62% |
| **Sniffles(M)** | 5x | 89.96% | 52.99% | 66.69% | 69.19% | 62.24% | 36.66% | 46.15% |
| **cuteSV(M)** | 5x | 89.56% | 54.37% | 67.66% | 64.28% | 57.57% | 34.95% | 43.49% |
| **gcSV(D)** | 4x | **90.47%** | **63.36%** | **74.53%** | **82.33%** | **74.48%** | **52.16%** | **61.35%** |
| **SVDSS(D)** | 4x | 86.89% | 48.99% | 62.65% | 56.96% | 49.49% | 27.90% | 35.69% |
| **Sniffles(M)** | 4x | 89.58% | 50.23% | 64.36% | 66.12% | 59.22% | 33.21% | 42.55% |
| **cuteSV(M)** | 4x | 88.99% | 52.03% | 65.66% | 61.45% | 54.69% | 31.97% | 40.35% |
| **gcSV(D)** | 3x | **90.79%** | **56.74%** | **69.84%** | **79.96%** | **72.60%** | **45.37%** | **55.84%** |
| **SVDSS(D)** | 3x | 87.21% | 39.25% | 54.13% | 54.89% | 47.87% | 21.54% | 29.71% |
| **Sniffles(M)** | 3x | 89.46% | 46.06% | 60.81% | 62.26% | 55.70% | 28.68% | 37.86% |
| **cuteSV(M)** | 3x | 88.59% | 48.16% | 62.40% | 58.02% | 51.40% | 27.94% | 36.20% |
| **gcSV(D)** | 2x | **90.92%** | **46.46%** | **61.49%** | **75.75%** | **68.87%** | **35.19%** | **46.58%** |
| **SVDSS(D)** | 2x | 87.62% | 25.67% | 39.70% | 50.81% | 44.52% | 13.04% | 20.17% |
| **Sniffles(M)** | 2x | 89.28% | 39.31% | 54.59% | 56.50% | 50.44% | 22.21% | 30.84% |
| **cuteSV(M)** | 2x | 88.14% | 41.52% | 56.45% | 53.63% | 47.27% | 22.27% | 30.28% |
| **gcSV(D)** | 1x | **91.51%** | **28.99%** | **44.03%** | **71.06%** | **65.03%** | **20.60%** | **31.29%** |
| **SVDSS(D)** | 1x | 87.72% | 9.61% | 17.32% | 44.62% | 39.14% | 4.29% | 7.73% |
| **Sniffles(M)** | 1x | 89.09% | 26.25% | 40.55% | 47.46% | 42.28% | 12.46% | 19.24% |
| **cuteSV(M)** | 1x | 87.51% | 27.96% | 42.38% | 45.94% | 40.20% | 12.85% | 19.47% |

(a) Ground truth callset: NIST HG002 Q100 benchmark (V1.0)

(b) The precisions, sensitivities and F1-scores of various approaches are assessed by Truvari with the following parameter setting "truvari bench --passonly -p 0 --dup-to-in"

(c) gcSV(D), SVDSS(D) indicate the tools were running with default parameter settings.

(d) Sniffles(M) and cuteSV(M) indicate the tools were running in "max sensitive setting" for low coverage sequencing data, i.e., (sniffles --minsupport 1 --qc-output-all --qc-coverage 1 --long-dup-coverage 1 --detect-large-ins True) for Sniffles and (cuteSV --genotype -s 1) for cuteSV

**Supplementary Table 3. The yields of various approaches using hybrid sequencing data [a]**

| Tool | Coverage | Precision [b] | Recall [b] | F1-Score-SV [b] | GT concordance [b] | Precision-GT [b] | Recall-GT [b] | F1-Score-GT [b] |
|---|---|---|---|---|---|---|---|---|
| **gcSV(D)** [c] | 1x+60x [d] | 90.62% | 48.92% | 63.54% | 79.44% | 71.99% | 38.86% | 50.47% |
| **gcSV(D)** | 2x+60x | 90.25% | 57.84% | 70.50% | 80.54% | 72.69% | 46.59% | 56.78% |
| **gcSV(D)** | 3x+60x | 90.60% | 63.69% | 74.79% | 82.80% | 75.01% | 52.73% | 61.93% |
| **gcSV(D)** | 4x+60x | 90.69% | 67.54% | 77.42% | 83.95% | 76.13% | 56.70% | 64.99% |
| **gcSV(D)** | 5x+60x | 90.69% | 70.05% | 79.04% | 85.52% | 77.56% | 59.91% | 67.60% |
| **gcSV(D)** | 10x+60x | **90.85%** | **77.36%** | **83.57%** | **90.68%** | **82.39%** | **70.15%** | **75.78%** |
| **gcSV(D)** | 1x+30x | 90.88% | 45.70% | 60.81% | 79.09% | 71.87% | 36.14% | 48.10% |
| **gcSV(D)** | 2x+30x | 90.47% | 55.95% | 69.14% | 80.38% | 72.72% | 44.97% | 55.58% |
| **gcSV(D)** | 3x+30x | 90.68% | 62.41% | 73.93% | 82.57% | 74.87% | 51.53% | 61.05% |
| **gcSV(D)** | 4x+30x | 90.82% | 66.67% | 76.89% | 83.72% | 76.04% | 55.82% | 64.38% |
| **gcSV(D)** | 5x+30x | 90.83% | 69.35% | 78.65% | 85.34% | 77.52% | 59.18% | 67.12% |
| **gcSV(D)** | 10x+30x | **90.93%** | **77.11%** | **83.45%** | **90.67%** | **82.44%** | **69.91%** | **75.66%** |
| **Blend-Seq** | 4x+30x | NA | NA | NA | NA | 65.17% | 41.79% | 50.92% |

(a) Ground truth callset: NIST HG002 Q100 benchmark (V1.0)

(b) The precisions, sensitivities and F1-scores of various approaches are assessed by Truvari with the following parameter setting "truvari bench --passonly -p 0 --dup-to-in"

(c) gcSV was running with default parameter setting.

(d) "Ax+Bx" means hybrid SV-calling using Ax coverage PacBio HiFi reads plus Bx coverage Illumina short reads

**Supplementary Table 4. The yields of various approaches using short read sequencing data [a]**

| Tool | Coverage | Precision [b] | Recall [b] | F1-Score-SV [b] | GT concordance [b] | Precision-GT [b] | Recall-GT [b] | F1-Score-GT [b] |
|---|---|---|---|---|---|---|---|---|
| **gcSV [c]** | 60x | **91.48%** | **35.39%** | **51.03%** | 81.11% | **74.20%** | **28.70%** | **41.39%** |
| **Manta [c]** | 60x | 87.54% | 23.29% | 36.79% | 81.06% | 70.96% | 18.88% | 29.82% |
| **Lumpy [c]** | 60x | 69.18% | 10.19% | 17.77% | 0.00% | 0.00% | 0.00% | 0.00% |
| **Delly [c]** | 60x | 61.42% | 11.43% | 19.28% | **91.91%** | 56.45% | 10.51% | 17.72% |
| **gcSV** | 35x [d] | **91.04%** | **30.07%** | **45.21%** | **83.64%** | **76.15%** | **25.15%** | **37.81%** |
| **DRAGEN** | 35x | 86.44% | 29.99% | 44.53% | 78.37% | 67.75% | 23.50% | 34.90% |
| **GATK-SV** | 35x | 78.30% | 21.26% | 33.44% | 81.88% | 64.11% | 17.41% | 27.38% |
| **gcSV** | 30x | **92.02%** | **29.78%** | **44.99%** | 81.59% | **75.08%** | **24.29%** | **36.71%** |
| **Manta** | 30x | 89.74% | 19.09% | 31.48% | 82.64% | 74.16% | 15.78% | 26.02% |
| **Lumpy** | 30x | 80.92% | 9.51% | 17.03% | 0.00% | 0.00% | 0.00% | 0.00% |
| **Delly** | 30x | 73.51% | 10.50% | 18.38% | **93.57%** | 68.78% | 9.83% | 17.20% |
| **gcSV** | 15x [d] | **93.22%** | **17.69%** | **29.73%** | 82.92% | 77.29% | **14.67%** | **24.65%** |
| **Manta** | 15x | 92.42% | 13.20% | 23.10% | 86.76% | **80.19%** | 11.45% | 20.04% |
| **Lumpy** | 15x | 91.80% | 6.23% | 11.66% | 0.00% | 0.00% | 0.00% | 0.00% |
| **Delly** | 15x | 82.53% | 8.92% | 16.10% | **93.88%** | 77.48% | 8.37% | 15.12% |

(a) Ground truth callset: NIST HG002 Q100 benchmark (V1.0)

(b) The precisions, sensitivities and F1-scores of various approaches are assessed by Truvari with the following parameter setting "truvari bench --passonly -p 0 --dup-to-in"

(c) All the tools were running with their default parameter settings.

(d) The 30x and 15x datasets were obtained by downsampling the full (60x) Illumina dataset.

**Supplementary Table 5. The yields of various approaches for SV sizes and types (long reads) [a]**

| Tool | SV Size | SV Type | Precision [b] | Recall [b] | F1-Score-SV [b] | GT concordance [b] | Precision-GT [b] | Recall-GT [b] | F1-Score-GT [b] |
|---|---|---|---|---|---|---|---|---|---|
| **gcSV [c]** | 50-200bp | INS | 88.91% | **84.29%** | **86.53%** | **94.77%** | **84.26%** | **79.88%** | **82.01%** |
| **SVDSS [c]** | 50-200bp | INS | 80.70% | 78.55% | 79.61% | 52.25% | 42.17% | 41.04% | 41.60% |
| **Sniffles [c]** | 50-200bp | INS | **89.68%** | 64.94% | 75.33% | 80.90% | 72.55% | 52.54% | 60.94% |
| **cuteSV [c]** | 50-200bp | INS | 88.99% | 61.40% | 72.66% | 80.13% | 71.31% | 49.20% | 58.23% |
| **gcSV** | 200-500bp | INS | 91.63% | **88.84%** | **90.21%** | **93.91%** | **86.04%** | **83.43%** | **84.71%** |
| **SVDSS** | 200-500bp | INS | 83.94% | 83.01% | 83.47% | 48.78% | 40.95% | 40.50% | 40.72% |
| **Sniffles** | 200-500bp | INS | 92.02% | 69.35% | 79.09% | 76.04% | 69.97% | 52.74% | 60.14% |
| **cuteSV** | 200-500bp | INS | **92.86%** | 62.25% | 74.54% | 73.31% | 68.07% | 45.64% | 54.64% |
| **gcSV** | 500-1kbp | INS | 89.20% | **89.39%** | **89.30%** | **92.94%** | **82.90%** | **83.07%** | **82.99%** |
| **SVDSS** | 500-1kbp | INS | 82.93% | 83.49% | 83.21% | 31.68% | 26.27% | 26.45% | 26.36% |
| **Sniffles** | 500-1kbp | INS | 88.22% | 62.42% | 73.11% | 60.20% | 53.11% | 37.58% | 44.01% |
| **cuteSV** | 500-1kbp | INS | **90.21%** | 54.35% | 67.83% | 57.33% | 51.72% | 31.16% | 38.89% |
| **gcSV** | 1k-2kbp | INS | **91.92%** | **88.28%** | **90.06%** | **94.59%** | **86.95%** | **83.50%** | **85.19%** |
| **SVDSS** | 1k-2kbp | INS | 87.11% | 81.06% | 83.98% | 34.71% | 30.23% | 28.13% | 29.15% |
| **Sniffles** | 1k-2kbp | INS | 89.00% | 57.62% | 69.95% | 62.60% | 55.71% | 36.07% | 43.79% |
| **cuteSV** | 1k-2kbp | INS | 89.46% | 45.18% | 60.04% | 57.29% | 51.25% | 25.88% | 34.39% |
| **gcSV** | 2k-5kbp | INS | 93.34% | **81.13%** | **86.81%** | **93.49%** | **87.26%** | **75.85%** | **81.16%** |
| **SVDSS** | 2k-5kbp | INS | 91.88% | 62.64% | 74.50% | 26.31% | 24.17% | 16.48% | 19.60% |
| **Sniffles** | 2k-5kbp | INS | 89.89% | 53.71% | 67.24% | 39.58% | 35.58% | 21.26% | 26.61% |
| **cuteSV** | 2k-5kbp | INS | **94.64%** | 33.33% | 49.30% | 45.28% | 42.86% | 15.09% | 22.33% |
| **gcSV** | >5kbp | INS | 96.74% | **25.57%** | **40.45%** | **82.02%** | **79.35%** | **20.98%** | **33.18%** |
| **SVDSS** | >5kbp | INS | **100.00%** | 0.29% | 0.57% | 0.00% | 0.00% | 0.00% | 0.00% |
| **Sniffles** | >5kbp | INS | 78.95% | 4.31% | 8.17% | 20.00% | 15.79% | 0.86% | 1.63% |
| **cuteSV** | >5kbp | INS | **100.00%** | 4.60% | 8.79% | 31.25% | 31.25% | 1.44% | 2.75% |
| **gcSV** | 50-200bp | DEL | 90.90% | **76.98%** | **83.36%** | **94.71%** | **86.09%** | **72.90%** | **78.95%** |
| **SVDSS** | 50-200bp | DEL | 86.03% | 71.36% | 78.01% | 64.02% | 55.07% | 45.68% | 49.94% |
| **Sniffles** | 50-200bp | DEL | 89.57% | 63.81% | 74.53% | 89.00% | 79.71% | 56.79% | 66.32% |
| **cuteSV** | 50-200bp | DEL | **91.67%** | 59.48% | 72.15% | 85.76% | 78.62% | 51.01% | 61.88% |
| **gcSV** | 200-500bp | DEL | 93.39% | **84.96%** | **88.97%** | **95.55%** | **89.23%** | **81.18%** | **85.02%** |
| **SVDSS** | 200-500bp | DEL | 89.75% | 80.33% | 84.78% | 77.92% | 69.93% | 62.59% | 66.06% |
| **Sniffles** | 200-500bp | DEL | 92.51% | 77.67% | 84.44% | 94.29% | 87.23% | 73.24% | 79.62% |

| | | | | | | | | | |
|---|---|---|---|---|---|---|---|---|---|
| **cuteSV** | 200-500bp | DEL | **93.52%** | 71.85% | 81.26% | 92.16% | 86.19% | 66.22% | 74.90% |
| **gcSV** | 500-1kbp | DEL | **91.89%** | **76.26%** | **83.35%** | **94.61%** | **86.94%** | **72.15%** | **78.86%** |
| **SVDSS** | 500-1kbp | DEL | 81.14% | 74.77% | 77.82% | 59.75% | 48.48% | 44.67% | 46.50% |
| **Sniffles** | 500-1kbp | DEL | 84.60% | 70.84% | 77.11% | 91.29% | 77.23% | 64.67% | 70.40% |
| **cuteSV** | 500-1kbp | DEL | 85.71% | 58.32% | 69.41% | 87.50% | 75.00% | 51.03% | 60.73% |
| **gcSV** | 1k-2kbp | DEL | **94.01%** | 79.46% | 86.13% | **98.13%** | **92.25%** | **77.98%** | **84.52%** |
| **SVDSS** | 1k-2kbp | DEL | 91.72% | **82.44%** | **86.83%** | 52.71% | 48.34% | 43.45% | 45.77% |
| **Sniffles** | 1k-2kbp | DEL | 91.67% | 75.30% | 82.68% | 93.28% | 85.51% | 70.24% | 77.12% |
| **cuteSV** | 1k-2kbp | DEL | 91.54% | 54.76% | 68.53% | 90.22% | 82.59% | 49.40% | 61.82% |
| **gcSV** | 2k-5kbp | DEL | 90.00% | **66.44%** | **76.45%** | **93.43%** | **84.09%** | **62.08%** | **71.43%** |
| **SVDSS** | 2k-5kbp | DEL | 93.94% | 62.42% | 75.00% | 59.68% | 56.06% | 37.25% | 44.76% |
| **Sniffles** | 2k-5kbp | DEL | **95.53%** | 57.38% | 71.70% | 90.06% | 86.03% | 51.68% | 64.57% |
| **cuteSV** | 2k-5kbp | DEL | 94.57% | 40.94% | 57.14% | 91.80% | 86.82% | 37.58% | 52.46% |
| **gcSV** | >5kbp | DEL | **100.00%** | **45.15%** | **62.21%** | 89.25% | **89.25%** | **40.29%** | **55.52%** |
| **SVDSS** | >5kbp | DEL | **100.00%** | 0.97% | 1.92% | 50.00% | 50.00% | 0.49% | 0.96% |
| **Sniffles** | >5kbp | DEL | 95.65% | 10.68% | 19.21% | 40.91% | 39.13% | 4.37% | 7.86% |
| **cuteSV** | >5kbp | DEL | 90.00% | 8.74% | 15.93% | **94.44%** | 85.00% | 8.25% | 15.04% |

(a) Ground truth callset: NIST HG002 Q100 benchmark (V1.0)

(b) The precisions, sensitivities and F1-scores of various approaches are assessed by Truvari with the following parameter setting "truvari bench --passonly -p 0 --dup-to-in"

(c) All the tools were running with their default parameter settings.

**Supplementary Table 6. The yields of various approaches in various genomic regions (long reads) [a]**

| Tool | Genomic Regions | Precision [b] | Recall [b] | F1-Score-SV [b] | GT concordance [b] | Precision-GT [b] | Recall-GT [b] | F1-Score-GT [b] |
|---|---|---|---|---|---|---|---|---|
| **gcSV [c]** | Simple repeat | **86.95%** | **78.62%** | **82.57%** | **93.91%** | **81.65%** | **73.82%** | **77.54%** |
| **SVDSS [c]** | Simple repeat | 80.94% | 70.39% | 75.30% | 37.93% | 30.70% | 26.70% | 28.56% |
| **Sniffles [c]** | Simple repeat | 84.83% | 54.96% | 66.71% | 71.81% | 60.92% | 39.47% | 47.90% |
| **cuteSV [c]** | Simple repeat | 85.37% | 49.63% | 62.77% | 68.53% | 58.51% | 34.01% | 43.02% |
| **gcSV** | LINE | **92.99%** | **84.57%** | **88.58%** | **93.61%** | **87.05%** | **79.17%** | **82.92%** |
| **SVDSS** | LINE | 85.27% | 81.02% | 83.09% | 61.71% | 52.63% | 50.00% | 51.28% |
| **Sniffles** | LINE | 90.09% | 70.63% | 79.18% | 80.99% | 72.97% | 57.20% | 64.13% |
| **cuteSV** | LINE | 91.49% | 63.58% | 75.02% | 79.53% | 72.76% | 50.57% | 59.67% |
| **gcSV** | SINE | **94.24%** | **80.26%** | **86.69%** | **92.03%** | **86.73%** | **73.86%** | **79.78%** |
| **SVDSS** | SINE | 86.34% | 78.23% | 82.08% | 56.12% | 48.45% | 43.90% | 46.06% |
| **Sniffles** | SINE | 92.37% | 63.76% | 75.44% | 72.42% | 66.90% | 46.17% | 54.64% |
| **cuteSV** | SINE | 92.50% | 56.82% | 70.40% | 71.16% | 65.82% | 40.43% | 50.09% |
| **gcSV** | Low complexity | **85.52%** | **78.17%** | **81.68%** | **95.08%** | **81.30%** | **74.32%** | **77.66%** |
| **SVDSS** | Low complexity | 78.93% | 71.74% | 75.16% | 39.86% | 31.46% | 28.59% | 29.96% |
| **Sniffles** | Low complexity | 84.14% | 53.57% | 65.46% | 69.50% | 58.48% | 37.23% | 45.50% |
| **cuteSV** | Low complexity | 84.92% | 45.21% | 59.01% | 69.68% | 59.17% | 31.50% | 41.12% |
| **gcSV** | LTR | **93.01%** | **79.96%** | **85.99%** | **93.54%** | **86.99%** | **74.79%** | **80.43%** |
| **SVDSS** | LTR | 88.55% | 76.69% | 82.19% | 71.80% | 63.58% | 55.06% | 59.02% |
| **Sniffles** | LTR | 87.73% | 71.62% | 78.86% | 86.89% | 76.23% | 62.24% | 68.52% |
| **cuteSV** | LTR | 91.10% | 62.66% | 74.25% | 87.04% | 79.29% | 54.54% | 64.63% |
| **gcSV** | Satellite | **66.77%** | **54.09%** | **59.76%** | **90.22%** | **60.24%** | **48.80%** | **53.92%** |
| **SVDSS** | Satellite | 45.53% | 51.44% | 48.31% | 50.93% | 23.19% | 26.20% | 24.60% |
| **Sniffles** | Satellite | 49.50% | 47.60% | 48.53% | 79.80% | 39.50% | 37.98% | 38.73% |
| **cuteSV** | Satellite | 47.81% | 39.42% | 43.21% | 79.88% | 38.19% | 31.49% | 34.52% |
| **gcSV** | DNA | **94.00%** | **81.03%** | **87.04%** | **97.34%** | **91.50%** | **78.88%** | **84.72%** |
| **SVDSS** | DNA | 86.10% | 82.76% | 84.40% | 70.31% | 60.54% | 58.19% | 59.34% |
| **Sniffles** | DNA | 90.31% | 76.29% | 82.71% | 87.01% | 78.57% | 66.38% | 71.96% |
| **cuteSV** | DNA | 93.06% | 69.40% | 79.51% | 86.96% | 80.92% | 60.34% | 69.14% |
| **gcSV** | RNA | 90.00% | **90.00%** | **90.00%** | **100.00%** | **90.00%** | **90.00%** | **90.00%** |
| **SVDSS** | RNA | 85.71% | 60.00% | 70.59% | 33.33% | 28.57% | 20.00% | 23.53% |
| **Sniffles** | RNA | **100.00%** | 50.00% | 66.67% | 60.00% | 60.00% | 30.00% | 40.00% |

| | | | | | | | | |
|---|---|---|---|---|---|---|---|---|
| **cuteSV** | RNA | **100.00%** | 40.00% | 57.14% | 75.00% | 75.00% | 30.00% | 42.86% |
| **gcSV** | CMRG | 95.05% | 94.58% | **94.81%** | **96.35%** | **91.58%** | **91.13%** | **91.36%** |
| **SVDSS** | CMRG | 89.45% | **96.06%** | 92.64% | 78.46% | 70.18% | 75.37% | 72.68% |
| **Sniffles** | CMRG | **98.37%** | 89.16% | 93.54% | 91.71% | 90.22% | 81.77% | 85.79% |
| **cuteSV** | CMRG | 98.18% | 79.80% | 88.04% | 91.36% | 89.70% | 72.91% | 80.43% |

(a) Ground truth callset: NIST HG002 Q100 benchmark (V1.0)

(b) The precisions, sensitivities and F1-scores of various approaches are assessed by Truvari with the following parameter setting "truvari bench --passonly -p 0 --dup-to-in"

(c) All the tools were running with their default parameter settings.

**Supplementary Table 7. The yields of gcSV for various SV sizes and types (hybrid sequencing) [a]**

| Coverage | SV Size | SV Type | Precision [b] | Recall [b] | F1-Score-SV [b] | GT concordance [b] | Precision-GT [b] | Recall-GT [b] | F1-Score-GT [b] |
|---|---|---|---|---|---|---|---|---|---|
| 1x+60x | 50-200bp | INS | 89.30% | 47.78% | 62.25% | 75.14% | 67.10% | 35.90% | 46.77% |
| 2x+60x | 50-200bp | INS | 88.36% | 57.95% | 69.99% | 78.17% | 69.07% | 45.30% | 54.72% |
| 3x+60x | 50-200bp | INS | 88.43% | 64.41% | 74.53% | 81.23% | 71.83% | 52.32% | 60.54% |
| 4x+60x | 50-200bp | INS | 88.51% | 68.95% | 77.51% | 83.16% | 73.61% | 57.34% | 64.46% |
| 5x+60x | 50-200bp | INS | 88.50% | 71.69% | 79.21% | 85.14% | 75.35% | 61.04% | 67.44% |
| 10x+60x | 50-200bp | INS | 88.25% | 79.70% | 83.76% | 91.16% | 80.45% | 72.66% | 76.35% |
| 1x+30x | 50-200bp | INS | 89.66% | 44.44% | 59.43% | 74.43% | 66.74% | 33.08% | 44.23% |
| 2x+30x | 50-200bp | INS | 88.74% | 56.02% | 68.68% | 77.59% | 68.86% | 43.47% | 53.29% |
| 3x+30x | 50-200bp | INS | 88.67% | 63.02% | 73.68% | 80.95% | 71.78% | 51.01% | 59.64% |
| 4x+30x | 50-200bp | INS | 88.73% | 67.94% | 76.95% | 82.85% | 73.51% | 56.29% | 63.76% |
| 5x+30x | 50-200bp | INS | 88.69% | 70.94% | 78.83% | 84.86% | 75.26% | 60.20% | 66.89% |
| 10x+30x | 50-200bp | INS | 88.33% | 79.32% | 83.58% | 91.19% | 80.54% | 72.33% | 76.22% |
| 1x+60x | 50-200bp | DEL | 88.08% | 51.04% | 64.63% | 84.99% | 74.86% | 43.38% | 54.93% |
| 2x+60x | 50-200bp | DEL | 88.50% | 57.85% | 69.97% | 85.19% | 75.39% | 49.28% | 59.60% |
| 3x+60x | 50-200bp | DEL | 89.52% | 62.56% | 73.65% | 87.53% | 78.36% | 54.76% | 64.47% |
| 4x+60x | 50-200bp | DEL | 89.90% | 65.56% | 75.82% | 88.08% | 79.18% | 57.74% | 66.78% |
| 5x+60x | 50-200bp | DEL | 90.27% | 67.78% | 77.43% | 89.28% | 80.60% | 60.52% | 69.13% |
| 10x+60x | 50-200bp | DEL | 91.09% | 73.70% | 81.48% | 92.76% | 84.50% | 68.36% | 75.58% |
| 1x+30x | 50-200bp | DEL | 88.55% | 47.71% | 62.01% | 84.48% | 74.80% | 40.30% | 52.38% |
| 2x+30x | 50-200bp | DEL | 88.90% | 56.02% | 68.73% | 84.63% | 75.24% | 47.41% | 58.17% |
| 3x+30x | 50-200bp | DEL | 89.59% | 61.54% | 72.96% | 86.91% | 77.86% | 53.48% | 63.41% |
| 4x+30x | 50-200bp | DEL | 90.02% | 64.89% | 75.42% | 87.87% | 79.10% | 57.02% | 66.27% |
| 5x+30x | 50-200bp | DEL | 90.43% | 67.30% | 77.17% | 89.19% | 80.65% | 60.02% | 68.82% |
| 10x+30x | 50-200bp | DEL | 91.08% | 73.54% | 81.38% | 92.75% | 84.47% | 68.21% | 75.47% |
| 1x+60x | 200-500bp | INS | 93.89% | 49.57% | 64.89% | 75.60% | 70.98% | 37.48% | 49.05% |
| 2x+60x | 200-500bp | INS | 92.73% | 59.75% | 72.68% | 76.93% | 71.34% | 45.97% | 55.91% |
| 3x+60x | 200-500bp | INS | 92.44% | 66.44% | 77.31% | 78.91% | 72.94% | 52.43% | 61.01% |
| 4x+60x | 200-500bp | INS | 92.18% | 71.21% | 80.35% | 80.70% | 74.39% | 57.46% | 64.84% |
| 5x+60x | 200-500bp | INS | 91.77% | 74.08% | 81.98% | 82.41% | 75.63% | 61.05% | 67.56% |
| 10x+60x | 200-500bp | INS | 91.49% | 82.55% | 86.79% | 88.83% | 81.26% | 73.33% | 77.09% |
| 1x+30x | 200-500bp | INS | 93.46% | 45.80% | 61.48% | 75.41% | 70.48% | 34.54% | 46.36% |
| 2x+30x | 200-500bp | INS | 92.47% | 57.78% | 71.12% | 77.23% | 71.41% | 44.62% | 54.92% |
| 3x+30x | 200-500bp | INS | 92.35% | 65.06% | 76.34% | 78.62% | 72.61% | 51.16% | 60.02% |

| | | | | | | | | | |
|---|---|---|---|---|---|---|---|---|---|
| 4x+30x | 200-500bp | INS | 92.11% | 70.29% | 79.74% | 80.45% | 74.11% | 56.55% | 64.15% |
| 5x+30x | 200-500bp | INS | 91.78% | 73.50% | 81.63% | 82.29% | 75.53% | 60.48% | 67.18% |
| 10x+30x | 200-500bp | INS | 91.52% | 82.43% | 86.73% | 88.86% | 81.32% | 73.25% | 77.07% |
| 1x+60x | 200-500bp | DEL | 88.64% | 65.29% | 75.19% | 90.90% | 80.58% | 59.35% | 68.35% |
| 2x+60x | 200-500bp | DEL | 89.15% | 71.27% | 79.21% | 89.72% | 79.98% | 63.94% | 71.07% |
| 3x+60x | 200-500bp | DEL | 90.14% | 75.43% | 82.13% | 90.29% | 81.38% | 68.11% | 74.15% |
| 4x+60x | 200-500bp | DEL | 91.17% | 77.21% | 83.61% | 91.31% | 83.24% | 70.50% | 76.34% |
| 5x+60x | 200-500bp | DEL | 92.12% | 78.48% | 84.76% | 92.33% | 85.06% | 72.46% | 78.26% |
| 10x+60x | 200-500bp | DEL | 92.92% | 82.45% | 87.37% | 93.92% | 87.27% | 77.44% | 82.06% |
| 1x+30x | 200-500bp | DEL | 89.77% | 62.98% | 74.03% | 90.69% | 81.42% | 57.12% | 67.14% |
| 2x+30x | 200-500bp | DEL | 89.77% | 69.69% | 78.46% | 90.26% | 81.02% | 62.90% | 70.82% |
| 3x+30x | 200-500bp | DEL | 90.32% | 74.51% | 81.66% | 90.94% | 82.14% | 67.76% | 74.26% |
| 4x+30x | 200-500bp | DEL | 91.46% | 76.82% | 83.50% | 91.06% | 83.29% | 69.96% | 76.04% |
| 5x+30x | 200-500bp | DEL | 92.26% | 78.17% | 84.63% | 92.16% | 85.03% | 72.04% | 78.00% |
| 10x+30x | 200-500bp | DEL | 93.16% | 82.41% | 87.46% | 93.87% | 87.45% | 77.36% | 82.10% |
| 1x+60x | 500-1kbp | INS | 89.05% | 35.35% | 50.61% | 63.84% | 56.84% | 22.57% | 32.31% |
| 2x+60x | 500-1kbp | INS | 87.85% | 49.79% | 63.56% | 68.81% | 60.46% | 34.27% | 43.74% |
| 3x+60x | 500-1kbp | INS | 86.22% | 59.27% | 70.25% | 73.54% | 63.40% | 43.58% | 51.66% |
| 4x+60x | 500-1kbp | INS | 85.32% | 64.39% | 73.39% | 75.24% | 64.20% | 48.45% | 55.22% |
| 5x+60x | 500-1kbp | INS | 84.77% | 68.01% | 75.47% | 77.02% | 65.29% | 52.38% | 58.13% |
| 10x+60x | 500-1kbp | INS | 85.65% | 80.64% | 83.07% | 86.46% | 74.05% | 69.72% | 71.82% |
| 1x+30x | 500-1kbp | INS | 88.78% | 32.35% | 47.42% | 66.24% | 58.81% | 21.43% | 31.41% |
| 2x+30x | 500-1kbp | INS | 87.78% | 48.34% | 62.35% | 70.24% | 61.65% | 33.95% | 43.79% |
| 3x+30x | 500-1kbp | INS | 86.29% | 58.64% | 69.83% | 74.14% | 63.98% | 43.48% | 51.77% |
| 4x+30x | 500-1kbp | INS | 85.60% | 63.98% | 73.22% | 75.73% | 64.82% | 48.45% | 55.45% |
| 5x+30x | 500-1kbp | INS | 85.13% | 67.55% | 75.32% | 77.39% | 65.88% | 52.28% | 58.30% |
| 10x+30x | 500-1kbp | INS | 85.82% | 80.18% | 82.90% | 86.31% | 74.07% | 69.20% | 71.55% |
| 1x+60x | 500-1kbp | DEL | 89.93% | 48.41% | 62.94% | 88.42% | 79.51% | 42.80% | 55.65% |
| 2x+60x | 500-1kbp | DEL | 89.49% | 55.70% | 68.66% | 88.93% | 79.58% | 49.53% | 61.06% |
| 3x+60x | 500-1kbp | DEL | 90.22% | 60.37% | 72.34% | 89.47% | 80.73% | 54.02% | 64.73% |
| 4x+60x | 500-1kbp | DEL | 91.58% | 62.99% | 74.64% | 88.43% | 80.98% | 55.70% | 66.00% |
| 5x+60x | 500-1kbp | DEL | 91.91% | 65.79% | 76.69% | 90.91% | 83.55% | 59.81% | 69.72% |
| 10x+60x | 500-1kbp | DEL | 91.73% | 70.47% | 79.70% | 94.16% | 86.37% | 66.36% | 75.05% |
| 1x+30x | 500-1kbp | DEL | 89.68% | 47.10% | 61.76% | 87.30% | 78.29% | 41.12% | 53.92% |
| 2x+30x | 500-1kbp | DEL | 89.26% | 54.39% | 67.60% | 89.69% | 80.06% | 48.79% | 60.63% |

| | | | | | | | | | |
|---|---|---|---|---|---|---|---|---|---|
| 3x+30x | 500-1kbp | DEL | 90.60% | 59.44% | 71.78% | 89.31% | 80.91% | 53.08% | 64.11% |
| 4x+30x | 500-1kbp | DEL | 91.41% | 61.68% | 73.66% | 88.79% | 81.16% | 54.77% | 65.40% |
| 5x+30x | 500-1kbp | DEL | 92.00% | 64.49% | 75.82% | 89.86% | 82.67% | 57.94% | 68.13% |
| 10x+30x | 500-1kbp | DEL | 91.93% | 70.28% | 79.66% | 94.41% | 86.80% | 66.36% | 75.21% |
| 1x+60x | 1k-2kbp | INS | 92.37% | 32.73% | 48.34% | 61.98% | 57.25% | 20.29% | 29.96% |
| 2x+60x | 1k-2kbp | INS | 90.40% | 45.00% | 60.08% | 68.74% | 62.14% | 30.93% | 41.30% |
| 3x+60x | 1k-2kbp | INS | 92.31% | 53.02% | 67.35% | 71.94% | 66.41% | 38.14% | 48.45% |
| 4x+60x | 1k-2kbp | INS | 91.75% | 59.15% | 71.93% | 73.48% | 67.41% | 43.46% | 52.85% |
| 5x+60x | 1k-2kbp | INS | 91.07% | 63.48% | 74.81% | 76.70% | 69.86% | 48.69% | 57.39% |
| 10x+60x | 1k-2kbp | INS | 91.65% | 75.20% | 82.62% | 85.73% | 78.57% | 64.47% | 70.83% |
| 1x+30x | 1k-2kbp | INS | 93.77% | 31.20% | 46.82% | 61.27% | 57.45% | 19.12% | 28.69% |
| 2x+30x | 1k-2kbp | INS | 90.93% | 44.27% | 59.55% | 68.64% | 62.41% | 30.39% | 40.87% |
| 3x+30x | 1k-2kbp | INS | 92.66% | 52.39% | 66.94% | 71.94% | 66.67% | 37.69% | 48.16% |
| 4x+30x | 1k-2kbp | INS | 91.94% | 58.61% | 71.59% | 73.23% | 67.33% | 42.92% | 52.42% |
| 5x+30x | 1k-2kbp | INS | 91.24% | 62.94% | 74.49% | 75.93% | 69.28% | 47.79% | 56.56% |
| 10x+30x | 1k-2kbp | INS | 91.83% | 75.02% | 82.58% | 85.22% | 78.26% | 63.93% | 70.37% |
| 1x+60x | 1k-2kbp | DEL | 96.09% | 65.77% | 78.09% | 95.02% | 91.30% | 62.50% | 74.20% |
| 2x+60x | 1k-2kbp | DEL | 97.05% | 68.45% | 80.28% | 91.30% | 88.61% | 62.50% | 73.30% |
| 3x+60x | 1k-2kbp | DEL | 96.36% | 70.83% | 81.65% | 94.12% | 90.69% | 66.67% | 76.84% |
| 4x+60x | 1k-2kbp | DEL | 95.67% | 72.32% | 82.37% | 93.83% | 89.76% | 67.86% | 77.29% |
| 5x+60x | 1k-2kbp | DEL | 94.51% | 71.73% | 81.56% | 93.78% | 88.63% | 67.26% | 76.48% |
| 10x+60x | 1k-2kbp | DEL | 94.03% | 75.00% | 83.44% | 96.03% | 90.30% | 72.02% | 80.13% |
| 1x+30x | 1k-2kbp | DEL | 97.21% | 62.20% | 75.86% | 95.22% | 92.56% | 59.23% | 72.23% |
| 2x+30x | 1k-2kbp | DEL | 98.20% | 64.88% | 78.14% | 91.74% | 90.09% | 59.52% | 71.68% |
| 3x+30x | 1k-2kbp | DEL | 97.03% | 68.15% | 80.07% | 93.89% | 91.10% | 63.99% | 75.17% |
| 4x+30x | 1k-2kbp | DEL | 96.34% | 70.54% | 81.44% | 93.67% | 90.24% | 66.07% | 76.29% |
| 5x+30x | 1k-2kbp | DEL | 95.16% | 70.24% | 80.82% | 94.49% | 89.92% | 66.37% | 76.37% |
| 10x+30x | 1k-2kbp | DEL | 93.98% | 74.40% | 83.06% | 95.60% | 89.85% | 71.13% | 79.40% |
| 1x+60x | 2k-5kbp | INS | 94.70% | 33.71% | 49.72% | 58.58% | 55.48% | 19.75% | 29.13% |
| 2x+60x | 2k-5kbp | INS | 93.04% | 42.01% | 57.89% | 61.38% | 57.10% | 25.79% | 35.53% |
| 3x+60x | 2k-5kbp | INS | 93.41% | 48.18% | 63.57% | 66.84% | 62.44% | 32.20% | 42.49% |
| 4x+60x | 2k-5kbp | INS | 93.32% | 52.70% | 67.36% | 67.78% | 63.25% | 35.72% | 45.66% |
| 5x+60x | 2k-5kbp | INS | 93.53% | 56.35% | 70.33% | 69.87% | 65.34% | 39.37% | 49.14% |
| 10x+60x | 2k-5kbp | INS | 93.54% | 67.42% | 78.36% | 79.48% | 74.35% | 53.58% | 62.28% |
| 1x+30x | 2k-5kbp | INS | 93.39% | 30.19% | 45.63% | 54.58% | 50.97% | 16.48% | 24.90% |

| | | | | | | | | | |
|---|---|---|---|---|---|---|---|---|---|
| 2x+30x | 2k-5kbp | INS | 92.58% | 39.25% | 55.12% | 60.26% | 55.79% | 23.65% | 33.22% |
| 3x+30x | 2k-5kbp | INS | 93.15% | 46.16% | 61.73% | 65.94% | 61.42% | 30.44% | 40.71% |
| 4x+30x | 2k-5kbp | INS | 92.76% | 51.57% | 66.29% | 67.07% | 62.22% | 34.59% | 44.46% |
| 5x+30x | 2k-5kbp | INS | 93.21% | 55.22% | 69.35% | 70.16% | 65.39% | 38.74% | 48.66% |
| 10x+30x | 2k-5kbp | INS | 93.53% | 67.30% | 78.27% | 79.81% | 74.65% | 53.71% | 62.47% |
| 1x+60x | 2k-5kbp | DEL | 98.66% | 74.16% | 84.67% | 93.67% | 92.41% | 69.46% | 79.31% |
| 2x+60x | 2k-5kbp | DEL | 98.25% | 75.50% | 85.39% | 94.22% | 92.58% | 71.14% | 80.46% |
| 3x+60x | 2k-5kbp | DEL | 98.68% | 75.17% | 85.33% | 92.41% | 91.19% | 69.46% | 78.86% |
| 4x+60x | 2k-5kbp | DEL | 98.23% | 74.50% | 84.73% | 92.34% | 90.71% | 68.79% | 78.24% |
| 5x+60x | 2k-5kbp | DEL | 96.83% | 71.81% | 82.47% | 92.99% | 90.05% | 66.78% | 76.69% |
| 10x+60x | 2k-5kbp | DEL | 91.12% | 65.44% | 76.17% | 93.85% | 85.51% | 61.41% | 71.48% |
| 1x+30x | 2k-5kbp | DEL | 98.58% | 70.13% | 81.96% | 94.74% | 93.40% | 66.44% | 77.65% |
| 2x+30x | 2k-5kbp | DEL | 98.62% | 72.15% | 83.33% | 94.42% | 93.12% | 68.12% | 78.68% |
| 3x+30x | 2k-5kbp | DEL | 99.07% | 71.81% | 83.27% | 92.99% | 92.13% | 66.78% | 77.43% |
| 4x+30x | 2k-5kbp | DEL | 98.60% | 71.14% | 82.65% | 92.92% | 91.63% | 66.11% | 76.80% |
| 5x+30x | 2k-5kbp | DEL | 97.14% | 68.46% | 80.31% | 93.63% | 90.95% | 64.09% | 75.20% |
| 10x+30x | 2k-5kbp | DEL | 91.04% | 64.77% | 75.69% | 93.78% | 85.38% | 60.74% | 70.98% |
| 1x+60x | >5kbp | INS | 92.05% | 23.28% | 37.16% | 56.79% | 52.27% | 13.22% | 21.10% |
| 2x+60x | >5kbp | INS | 95.70% | 25.57% | 40.36% | 57.30% | 54.84% | 14.66% | 23.13% |
| 3x+60x | >5kbp | INS | 97.87% | 26.44% | 41.63% | 56.52% | 55.32% | 14.94% | 23.53% |
| 4x+60x | >5kbp | INS | 97.96% | 27.59% | 43.05% | 55.21% | 54.08% | 15.23% | 23.77% |
| 5x+60x | >5kbp | INS | 96.97% | 27.59% | 42.95% | 55.21% | 53.54% | 15.23% | 23.71% |
| 10x+60x | >5kbp | INS | 97.83% | 25.86% | 40.91% | 58.89% | 57.61% | 15.23% | 24.09% |
| 1x+30x | >5kbp | INS | 88.73% | 18.10% | 30.07% | 49.21% | 43.66% | 8.91% | 14.80% |
| 2x+30x | >5kbp | INS | 90.24% | 21.26% | 34.42% | 50.00% | 45.12% | 10.63% | 17.21% |
| 3x+30x | >5kbp | INS | 91.95% | 22.99% | 36.78% | 48.75% | 44.83% | 11.21% | 17.93% |
| 4x+30x | >5kbp | INS | 94.57% | 25.00% | 39.55% | 50.57% | 47.83% | 12.64% | 20.00% |
| 5x+30x | >5kbp | INS | 93.41% | 24.43% | 38.72% | 50.59% | 47.25% | 12.36% | 19.59% |
| 10x+30x | >5kbp | INS | 97.75% | 25.00% | 39.82% | 57.47% | 56.18% | 14.37% | 22.88% |
| 1x+60x | >5kbp | DEL | 93.15% | 66.02% | 77.27% | 97.79% | 91.10% | 64.56% | 75.57% |
| 2x+60x | >5kbp | DEL | 93.20% | 66.50% | 77.62% | 97.81% | 91.16% | 65.05% | 75.92% |
| 3x+60x | >5kbp | DEL | 95.17% | 66.99% | 78.63% | 97.83% | 93.10% | 65.53% | 76.92% |
| 4x+60x | >5kbp | DEL | 95.17% | 66.99% | 78.63% | 97.83% | 93.10% | 65.53% | 76.92% |
| 5x+60x | >5kbp | DEL | 96.48% | 66.50% | 78.74% | 97.81% | 94.37% | 65.05% | 77.01% |
| 10x+60x | >5kbp | DEL | 99.08% | 52.43% | 68.57% | 95.37% | 94.50% | 50.00% | 65.40% |

| | | | | | | | | | |
|---|---|---|---|---|---|---|---|---|---|
| 1x+30x | >5kbp | DEL | 95.59% | 63.11% | 76.02% | 98.46% | 94.12% | 62.14% | 74.85% |
| 2x+30x | >5kbp | DEL | 95.62% | 63.59% | 76.38% | 98.47% | 94.16% | 62.62% | 75.22% |
| 3x+30x | >5kbp | DEL | 96.32% | 63.59% | 76.61% | 98.47% | 94.85% | 62.62% | 75.44% |
| 4x+30x | >5kbp | DEL | 96.32% | 63.59% | 76.61% | 98.47% | 94.85% | 62.62% | 75.44% |
| 5x+30x | >5kbp | DEL | 96.99% | 62.62% | 76.11% | 98.45% | 95.49% | 61.65% | 74.93% |
| 10x+30x | >5kbp | DEL | 99.07% | 51.94% | 68.15% | 95.33% | 94.44% | 49.51% | 64.97% |

(a) Ground truth callset: NIST HG002 Q100 benchmark (V1.0), and gcSV was running with its default parameters.
(b) The precisions, sensitivities and F1-scores of various approaches are assessed by Truvari with the following parameter setting "truvari bench --passonly -p 0 --dup-to-in"

**Supplementary Table 8. The yields of gcSV in various genomic regions (hybrid sequencing) [a]**

| Coverage | Region | Precision [b] | Recall [b] | F1-Score-SV [b] | GT concordance [b] | Precision-GT [b] | Recall-GT [b] | F1-Score-GT [b] |
|---|---|---|---|---|---|---|---|---|
| 1x+60x | Simple repeat | 86.17% | 38.42% | 53.15% | 71.48% | 61.59% | 27.46% | 37.99% |
| 2x+60x | Simple repeat | 85.55% | 48.15% | 61.62% | 76.22% | 65.21% | 36.70% | 46.97% |
| 3x+60x | Simple repeat | 85.87% | 55.21% | 67.21% | 79.50% | 68.27% | 43.89% | 53.43% |
| 4x+60x | Simple repeat | 86.09% | 60.18% | 70.84% | 81.33% | 70.01% | 48.94% | 57.61% |
| 5x+60x | Simple repeat | 86.29% | 63.32% | 73.04% | 83.34% | 71.92% | 52.77% | 60.88% |
| 10x+60x | Simple repeat | 86.13% | 71.98% | 78.42% | 89.07% | 76.71% | 64.11% | 69.85% |
| 1x+30x | Simple repeat | 86.55% | 35.82% | 50.67% | 71.67% | 62.03% | 25.68% | 36.32% |
| 2x+30x | Simple repeat | 85.85% | 46.72% | 60.51% | 76.26% | 65.46% | 35.62% | 46.14% |
| 3x+30x | Simple repeat | 86.01% | 54.27% | 66.55% | 79.54% | 68.42% | 43.17% | 52.94% |
| 4x+30x | Simple repeat | 86.23% | 59.52% | 70.43% | 81.16% | 69.98% | 48.30% | 57.16% |
| 5x+30x | Simple repeat | 86.49% | 62.85% | 72.80% | 83.22% | 71.98% | 52.31% | 60.59% |
| 10x+30x | Simple repeat | 86.24% | 71.74% | 78.32% | 89.06% | 76.80% | 63.89% | 69.75% |
| 1x+60x | LINE | 93.34% | 59.10% | 72.38% | 78.07% | 72.87% | 46.14% | 56.50% |
| 2x+60x | LINE | 92.12% | 64.92% | 76.16% | 79.24% | 72.99% | 51.44% | 60.35% |
| 3x+60x | LINE | 91.27% | 68.31% | 78.14% | 81.17% | 74.09% | 55.45% | 63.43% |
| 4x+60x | LINE | 91.84% | 71.19% | 80.21% | 81.94% | 75.25% | 58.33% | 65.72% |
| 5x+60x | LINE | 92.14% | 72.99% | 81.46% | 82.31% | 75.84% | 60.08% | 67.05% |
| 10x+60x | LINE | 92.78% | 79.99% | 85.91% | 88.94% | 82.52% | 71.14% | 76.41% |
| 1x+30x | LINE | 93.29% | 54.32% | 68.66% | 77.56% | 72.35% | 42.13% | 53.25% |
| 2x+30x | LINE | 91.93% | 61.52% | 73.71% | 78.76% | 72.41% | 48.46% | 58.06% |
| 3x+30x | LINE | 91.14% | 65.64% | 76.32% | 80.96% | 73.79% | 53.14% | 61.78% |
| 4x+30x | LINE | 92.00% | 69.19% | 78.98% | 81.71% | 75.17% | 56.53% | 64.53% |
| 5x+30x | LINE | 92.15% | 71.30% | 80.39% | 82.03% | 75.60% | 58.49% | 65.95% |
| 10x+30x | LINE | 92.76% | 79.73% | 85.75% | 89.16% | 82.70% | 71.09% | 76.46% |
| 1x+60x | SINE | 91.20% | 53.29% | 67.27% | 70.71% | 64.48% | 37.68% | 47.57% |
| 2x+60x | SINE | 91.09% | 59.33% | 71.86% | 68.55% | 62.44% | 40.67% | 49.26% |
| 3x+60x | SINE | 91.15% | 62.86% | 74.41% | 70.03% | 63.83% | 44.02% | 52.11% |
| 4x+60x | SINE | 91.47% | 65.43% | 76.29% | 72.58% | 66.39% | 47.49% | 55.37% |
| 5x+60x | SINE | 92.13% | 67.88% | 78.17% | 74.63% | 68.75% | 50.66% | 58.33% |
| 10x+60x | SINE | 92.64% | 73.74% | 82.12% | 82.73% | 76.63% | 61.00% | 67.93% |
| 1x+30x | SINE | 91.92% | 47.61% | 62.73% | 68.34% | 62.82% | 32.54% | 42.87% |
| 2x+30x | SINE | 91.84% | 56.58% | 70.02% | 67.44% | 61.94% | 38.16% | 47.22% |
| 3x+30x | SINE | 91.90% | 61.06% | 73.37% | 69.25% | 63.64% | 42.28% | 50.81% |

| | | | | | | | | |
|---|---|---|---|---|---|---|---|---|
| 4x+30x | SINE | 92.03% | 64.23% | 75.66% | 71.60% | 65.90% | 45.99% | 54.17% |
| 5x+30x | SINE | 92.32% | 66.87% | 77.56% | 73.97% | 68.29% | 49.46% | 57.37% |
| 10x+30x | SINE | 92.77% | 73.62% | 82.09% | 82.53% | 76.56% | 60.77% | 67.76% |
| 1x+60x | Low complexity | 85.61% | 27.09% | 41.16% | 73.31% | 62.76% | 19.86% | 30.17% |
| 2x+60x | Low complexity | 84.60% | 40.23% | 54.53% | 74.91% | 63.38% | 30.14% | 40.85% |
| 3x+60x | Low complexity | 84.23% | 47.89% | 61.06% | 78.14% | 65.81% | 37.42% | 47.71% |
| 4x+60x | Low complexity | 84.14% | 53.57% | 65.46% | 80.89% | 68.07% | 43.33% | 52.95% |
| 5x+60x | Low complexity | 84.20% | 57.56% | 68.38% | 82.22% | 69.23% | 47.32% | 56.22% |
| 10x+60x | Low complexity | 85.70% | 69.48% | 76.74% | 89.46% | 76.66% | 62.16% | 68.65% |
| 1x+30x | Low complexity | 85.53% | 24.98% | 38.66% | 71.80% | 61.41% | 17.93% | 27.76% |
| 2x+30x | Low complexity | 84.58% | 38.87% | 53.26% | 74.88% | 63.33% | 29.11% | 39.88% |
| 3x+30x | Low complexity | 84.47% | 47.00% | 60.39% | 77.72% | 65.65% | 36.53% | 46.94% |
| 4x+30x | Low complexity | 84.34% | 52.86% | 64.99% | 80.37% | 67.79% | 42.49% | 52.24% |
| 5x+30x | Low complexity | 84.44% | 56.81% | 67.92% | 82.31% | 69.50% | 46.76% | 55.91% |
| 10x+30x | Low complexity | 85.86% | 69.01% | 76.52% | 89.39% | 76.75% | 61.69% | 68.40% |
| 1x+60x | LTR | 92.51% | 59.92% | 72.73% | 84.68% | 78.34% | 50.74% | 61.59% |
| 2x+60x | LTR | 91.69% | 65.19% | 76.20% | 84.30% | 77.30% | 54.96% | 64.24% |
| 3x+60x | LTR | 90.38% | 67.41% | 77.22% | 85.76% | 77.51% | 57.81% | 66.22% |
| 4x+60x | LTR | 90.42% | 68.67% | 78.06% | 85.56% | 77.36% | 58.76% | 66.79% |
| 5x+60x | LTR | 91.72% | 70.15% | 79.50% | 87.67% | 80.41% | 61.50% | 69.70% |
| 10x+60x | LTR | 92.82% | 76.37% | 83.80% | 91.57% | 85.00% | 69.94% | 76.74% |
| 1x+30x | LTR | 91.29% | 55.27% | 68.86% | 82.82% | 75.61% | 45.78% | 57.03% |
| 2x+30x | LTR | 91.56% | 62.97% | 74.63% | 83.58% | 76.53% | 52.64% | 62.37% |
| 3x+30x | LTR | 90.30% | 65.82% | 76.14% | 85.10% | 76.85% | 56.01% | 64.80% |
| 4x+30x | LTR | 90.83% | 67.93% | 77.73% | 85.25% | 77.43% | 57.91% | 66.26% |
| 5x+30x | LTR | 92.20% | 69.83% | 79.47% | 86.86% | 80.08% | 60.65% | 69.03% |
| 10x+30x | LTR | 93.05% | 76.27% | 83.83% | 91.42% | 85.07% | 69.73% | 76.64% |
| 1x+60x | Satellite | 53.19% | 30.05% | 38.40% | 77.60% | 41.28% | 23.32% | 29.80% |
| 2x+60x | Satellite | 51.21% | 35.58% | 41.99% | 79.05% | 40.48% | 28.12% | 33.19% |
| 3x+60x | Satellite | 52.79% | 38.70% | 44.66% | 82.61% | 43.61% | 31.97% | 36.89% |
| 4x+60x | Satellite | 53.27% | 41.11% | 46.40% | 78.95% | 42.06% | 32.45% | 36.64% |
| 5x+60x | Satellite | 53.51% | 43.99% | 48.28% | 84.15% | 45.03% | 37.02% | 40.63% |
| 10x+60x | Satellite | 53.70% | 48.80% | 51.13% | 84.24% | 45.24% | 41.11% | 43.07% |
| 1x+30x | Satellite | 51.72% | 28.85% | 37.04% | 77.50% | 40.09% | 22.36% | 28.70% |
| 2x+30x | Satellite | 51.41% | 35.10% | 41.71% | 79.45% | 40.85% | 27.88% | 33.14% |

| | | | | | | | | |
|---|---|---|---|---|---|---|---|---|
| 3x+30x | Satellite | 53.14% | 38.70% | 44.78% | 82.61% | 43.89% | 31.97% | 37.00% |
| 4x+30x | Satellite | 53.63% | 40.87% | 46.38% | 78.82% | 42.27% | 32.21% | 36.56% |
| 5x+30x | Satellite | 54.01% | 43.75% | 48.34% | 84.07% | 45.40% | 36.78% | 40.64% |
| 10x+30x | Satellite | 53.72% | 48.56% | 51.01% | 84.16% | 45.21% | 40.87% | 42.93% |
| 1x+60x | DNA | 92.77% | 66.38% | 77.39% | 87.01% | 80.72% | 57.76% | 67.34% |
| 2x+60x | DNA | 90.34% | 68.53% | 77.94% | 84.28% | 76.14% | 57.76% | 65.69% |
| 3x+60x | DNA | 90.11% | 70.69% | 79.23% | 87.20% | 78.57% | 61.64% | 69.08% |
| 4x+60x | DNA | 90.76% | 71.98% | 80.29% | 88.02% | 79.89% | 63.36% | 70.67% |
| 5x+60x | DNA | 90.05% | 74.14% | 81.32% | 89.53% | 80.63% | 66.38% | 72.81% |
| 10x+60x | DNA | 92.31% | 77.59% | 84.31% | 93.89% | 86.67% | 72.84% | 79.16% |
| 1x+30x | DNA | 92.36% | 62.50% | 74.55% | 86.90% | 80.25% | 54.31% | 64.78% |
| 2x+30x | DNA | 89.71% | 67.67% | 77.15% | 85.35% | 76.57% | 57.76% | 65.85% |
| 3x+30x | DNA | 89.50% | 69.83% | 78.45% | 88.27% | 79.01% | 61.64% | 69.25% |
| 4x+30x | DNA | 90.66% | 71.12% | 79.71% | 89.09% | 80.77% | 63.36% | 71.01% |
| 5x+30x | DNA | 90.53% | 74.14% | 81.52% | 90.12% | 81.58% | 66.81% | 73.46% |
| 10x+30x | DNA | 92.31% | 77.59% | 84.31% | 93.89% | 86.67% | 72.84% | 79.16% |
| 1x+60x | RNA | 100.00% | 50.00% | 66.67% | 60.00% | 60.00% | 30.00% | 40.00% |
| 2x+60x | RNA | 100.00% | 50.00% | 66.67% | 60.00% | 60.00% | 30.00% | 40.00% |
| 3x+60x | RNA | 100.00% | 50.00% | 66.67% | 60.00% | 60.00% | 30.00% | 40.00% |
| 4x+60x | RNA | 100.00% | 50.00% | 66.67% | 80.00% | 80.00% | 40.00% | 53.33% |
| 5x+60x | RNA | 100.00% | 50.00% | 66.67% | 80.00% | 80.00% | 40.00% | 53.33% |
| 10x+60x | RNA | 85.71% | 60.00% | 70.59% | 66.67% | 57.14% | 40.00% | 47.06% |
| 1x+30x | RNA | 100.00% | 40.00% | 57.14% | 75.00% | 75.00% | 30.00% | 42.86% |
| 2x+30x | RNA | 100.00% | 50.00% | 66.67% | 60.00% | 60.00% | 30.00% | 40.00% |
| 3x+30x | RNA | 100.00% | 50.00% | 66.67% | 60.00% | 60.00% | 30.00% | 40.00% |
| 4x+30x | RNA | 100.00% | 50.00% | 66.67% | 80.00% | 80.00% | 40.00% | 53.33% |
| 5x+30x | RNA | 100.00% | 50.00% | 66.67% | 80.00% | 80.00% | 40.00% | 53.33% |
| 10x+30x | RNA | 85.71% | 60.00% | 70.59% | 66.67% | 57.14% | 40.00% | 47.06% |

(a) Ground truth callset: NIST HG002 Q100 benchmark (V1.0), and gcSV was running with its default parameters.
(b) The precisions, sensitivities and F1-scores of various approaches are assessed by Truvari with the following parameter setting "truvari bench --passonly -p 0 --dup-to-in"

**Supplementary Table 9. The yields of various approaches in the EASY/HARD regions for short reads [a]**

| Tools | Coverage | Region | Precision [b] | Recall [b] | F1-Score-SV [b] | GT concordance [b] | Precision-GT [b] | Recall-GT [b] | F1-Score-GT [b] |
|---|---|---|---|---|---|---|---|---|---|
| **gcSV [c]** | 60x | EASY | **96.98%** | **67.01%** | **79.26%** | **83.25%** | **80.73%** | **55.79%** | **65.98%** |
| **Delly [c]** | 60x | EASY | 73.79% | 13.29% | 22.52% | 93.46% | 68.97% | 12.42% | 21.05% |
| **Lumpy [c]** | 60x | EASY | 77.78% | 10.43% | 18.40% | 0.00% | 0.00% | 0.00% | 0.00% |
| **Manta [c]** | 60x | EASY | 90.88% | 35.89% | 51.46% | 87.68% | 79.69% | 31.47% | 45.12% |
| **gcSV** | 35X | EASY | **97.08%** | **60.69%** | **74.69%** | 84.98% | **82.50%** | **51.58%** | **63.47%** |
| **DRAGEN** | 35X | EASY | 92.90% | 49.68% | 64.74% | 84.82% | 78.80% | 42.14% | 54.91% |
| **GATK-SV** | 35X | EASY | 86.06% | 40.30% | 54.90% | **85.79%** | 73.82% | 34.58% | 47.10% |
| **gcSV** | 30x | EASY | **95.89%** | **55.66%** | **70.44%** | 81.79% | 78.43% | **45.53%** | **57.61%** |
| **Delly** | 30x | EASY | 81.41% | 12.30% | 21.36% | **94.55%** | 76.97% | 11.62% | 20.20% |
| **Lumpy** | 30x | EASY | 86.27% | 9.99% | 17.90% | 0.00% | 0.00% | 0.00% | 0.00% |
| **Manta** | 30x | EASY | 92.07% | 29.43% | 44.61% | 88.86% | **81.82%** | 26.15% | 39.64% |
| **gcSV** | 15x | EASY | **95.73%** | **32.31%** | **48.32%** | 79.40% | 76.01% | **25.66%** | **38.37%** |
| **Delly** | 15x | EASY | 85.94% | 9.56% | 17.21% | **94.55%** | **81.25%** | 9.04% | 16.27% |
| **Lumpy** | 15x | EASY | 89.90% | 6.86% | 12.74% | 0.00% | 0.00% | 0.00% | 0.00% |
| **Manta** | 15x | EASY | 94.09% | 21.36% | 34.82% | 91.28% | 85.89% | 19.50% | 31.78% |
| **gcSV** | 60x | HARD | **85.26%** | **23.49%** | **36.83%** | 75.01% | **63.95%** | **17.62%** | **27.63%** |
| **Delly** | 60x | HARD | 52.32% | 3.59% | 6.72% | **81.73%** | 42.76% | 2.93% | 5.49% |
| **Lumpy** | 60x | HARD | 42.90% | 2.47% | 4.67% | 0.00% | 0.00% | 0.00% | 0.00% |
| **Manta** | 60x | HARD | 82.89% | 13.96% | 23.90% | 70.21% | 58.20% | 9.80% | 16.78% |
| **gcSV** | 35X | HARD | **83.06%** | 18.39% | 30.11% | 77.98% | **64.78%** | **14.34%** | **23.48%** |
| **DRAGEN** | 35X | HARD | 77.65% | **19.00%** | **30.53%** | 68.62% | 53.28% | 13.04% | 20.95% |
| **GATK-SV** | 35X | HARD | 75.76% | 17.59% | 28.55% | **80.20%** | 60.76% | 14.09% | 22.88% |
| **gcSV** | 30x | HARD | **86.13%** | **18.41%** | **30.33%** | 74.73% | **64.36%** | **13.75%** | **22.66%** |
| **Delly** | 30x | HARD | 59.97% | 2.79% | 5.33% | **84.14%** | 50.46% | 2.35% | 4.49% |
| **Lumpy** | 30x | HARD | 47.93% | 1.97% | 3.79% | 0.00% | 0.00% | 0.00% | 0.00% |
| **Manta** | 30x | HARD | 85.39% | 10.25% | 18.30% | 70.09% | 59.84% | 7.18% | 12.82% |
| **gcSV** | 15x | HARD | **86.26%** | **9.62%** | **17.31%** | 76.69% | **66.16%** | **7.38%** | **13.28%** |
| **Delly** | 15x | HARD | 66.27% | 1.95% | 3.79% | **83.90%** | 55.60% | 1.64% | 3.18% |
| **Lumpy** | 15x | HARD | 42.37% | 0.87% | 1.71% | 0.00% | 0.00% | 0.00% | 0.00% |
| **Manta** | 15x | HARD | 87.71% | 5.62% | 10.57% | 72.95% | 63.98% | 4.10% | 7.71% |

(a) Ground truth callset: NIST HG002 Q100 benchmark (V1.0)

(b) The precisions, sensitivities and F1-scores of various approaches are assessed by Truvari with the following parameter setting “truvari bench --passonly -p 0 --dup-to-in”

(c) All the tools were running in the default parameters.

**Supplementary Table 10. The yields of various approaches for various SV sizes and types (short reads) [a]**

| Tools | SV Size | Precision [b] | Recall [b] | F1-Score-SV [b] | GT concordance [b] | Precision-GT [b] | Recall-GT [b] | F1-Score-GT [b] |
|---|---|---|---|---|---|---|---|---|
| | | | | 60X-INS | | | | |
| **gcSV [c]** | 50-200bp | 92.25% | **32.04%** | **47.56%** | 70.83% | **65.34%** | **22.69%** | **33.69%** |
| **Delly [c]** | 50-200bp | **94.15%** | 1.89% | 3.70% | **75.71%** | 71.28% | 1.43% | 2.80% |
| **Lumpy [c]** | 50-200bp | 52.54% | 0.33% | 0.66% | 0.00% | 0.00% | 0.00% | 0.00% |
| **Manta [c]** | 50-200bp | 86.29% | 21.93% | 34.97% | 65.54% | 56.56% | 14.37% | 22.92% |
| **gcSV** | 200-500bp | **96.59%** | **31.88%** | **47.93%** | **74.59%** | **72.05%** | **23.78%** | **35.75%** |
| **Delly** | 200-500bp | 39.49% | 1.27% | 2.46% | 53.23% | 21.02% | 0.68% | 1.31% |
| **Lumpy** | 200-500bp | 43.75% | 1.02% | 1.99% | 0.00% | 0.00% | 0.00% | 0.00% |
| **Manta** | 200-500bp | 94.29% | 10.31% | 18.58% | 64.85% | 61.14% | 6.68% | 12.05% |
| **gcSV** | 500-1kbp | **94.39%** | **14.80%** | **25.59%** | 56.64% | **53.47%** | **8.39%** | **14.50%** |
| **Delly** | 500-1kbp | 36.65% | 3.00% | 5.55% | 67.80% | 24.84% | 2.04% | 3.76% |
| **Lumpy** | 500-1kbp | 47.75% | 2.74% | 5.19% | 0.00% | 0.00% | 0.00% | 0.00% |
| **Manta** | 500-1kbp | 75.00% | 2.33% | 4.52% | **68.89%** | 51.67% | 1.60% | 3.11% |
| **gcSV** | 1k-2kbp | **87.65%** | **12.80%** | **22.34%** | 54.93% | **48.15%** | **7.03%** | **12.27%** |
| **Delly** | 1k-2kbp | 23.53% | 0.99% | 1.90% | 83.33% | 19.61% | 0.83% | 1.59% |
| **Lumpy** | 1k-2kbp | 25.93% | 1.26% | 2.41% | 0.00% | 0.00% | 0.00% | 0.00% |
| **Manta** | 1k-2kbp | 38.10% | 0.72% | 1.42% | **100.00%** | 38.10% | 0.72% | 1.42% |
| **gcSV** | 2k-5kbp | **94.19%** | **20.38%** | **33.51%** | 58.64% | **55.23%** | **11.95%** | **19.65%** |
| **Delly** | 2k-5kbp | 21.43% | 1.01% | 1.92% | 66.67% | 14.29% | 0.67% | 1.28% |
| **Lumpy** | 2k-5kbp | 18.37% | 1.13% | 2.13% | 0.00% | 0.00% | 0.00% | 0.00% |
| **Manta** | 2k-5kbp | 21.05% | 0.50% | 0.98% | **100.00%** | 21.05% | 0.50% | 0.98% |
| **gcSV** | >5kbp | **82.50%** | **18.97%** | **30.84%** | 57.58% | **47.50%** | **10.92%** | **17.76%** |
| **Delly** | >5kbp | 7.77% | 2.01% | 3.20% | 75.00% | 5.83% | 1.51% | 2.40% |
| **Lumpy** | >5kbp | 7.29% | 2.01% | 3.15% | 0.00% | 0.00% | 0.00% | 0.00% |
| **Manta** | >5kbp | 15.00% | 0.86% | 1.63% | **100.00%** | 15.00% | 0.86% | 1.63% |
| | | | | 60X-DEL | | | | |
| **gcSV** | 50-200bp | **86.92%** | **41.67%** | **56.33%** | 89.17% | **77.51%** | **37.15%** | **50.23%** |
| **Delly** | 50-200bp | 66.53% | 13.70% | 22.73% | **90.58%** | 60.26% | 12.41% | 20.59% |
| **Lumpy** | 50-200bp | 71.23% | 10.33% | 18.04% | 0.00% | 0.00% | 0.00% | 0.00% |
| **Manta** | 50-200bp | 85.80% | 31.03% | 45.57% | 84.82% | 72.77% | 26.32% | 38.65% |
| **gcSV** | 200-500bp | 88.26% | **56.85%** | **69.15%** | 95.66% | 84.43% | **54.38%** | **66.15%** |
| **Delly** | 200-500bp | 64.67% | 44.54% | 52.75% | 98.10% | 63.44% | 43.69% | 51.75% |

| | | | | | | | | |
|---|---|---|---|---|---|---|---|---|
| **Lumpy** | 200-500bp | 78.52% | 45.39% | 57.53% | 0.00% | 0.00% | 0.00% | 0.00% |
| **Manta** | 200-500bp | **89.88%** | 49.33% | 63.70% | **96.64%** | **86.86%** | 47.67% | 61.55% |
| **gcSV** | 500-1kbp | **94.64%** | **39.63%** | **55.86%** | 92.45% | **87.50%** | **36.64%** | **51.65%** |
| **Delly** | 500-1kbp | 36.32% | 42.43% | 39.14% | 87.22% | 31.68% | 37.01% | 34.14% |
| **Lumpy** | 500-1kbp | 50.75% | 44.30% | 47.31% | 0.00% | 0.00% | 0.00% | 0.00% |
| **Manta** | 500-1kbp | 88.00% | 32.90% | 47.89% | **96.59%** | 85.00% | 31.78% | 46.26% |
| **gcSV** | 1k-2kbp | **97.12%** | 60.12% | **74.26%** | 96.53% | **93.75%** | **58.04%** | **71.69%** |
| **Delly** | 1k-2kbp | 64.04% | **72.62%** | 68.06% | **97.95%** | 62.73% | 71.13% | 66.67% |
| **Lumpy** | 1k-2kbp | 71.22% | 71.43% | 71.32% | 0.00% | 0.00% | 0.00% | 0.00% |
| **Manta** | 1k-2kbp | 88.98% | 62.50% | 73.43% | 97.14% | 86.44% | 60.71% | 71.33% |
| **gcSV** | 2k-5kbp | **98.19%** | 72.82% | 83.62% | 94.93% | **93.21%** | 69.13% | **79.38%** |
| **Delly** | 2k-5kbp | 83.90% | 82.21% | 83.05% | 93.47% | 78.42% | 76.85% | 77.63% |
| **Lumpy** | 2k-5kbp | 85.86% | **83.56%** | **84.69%** | 0.00% | 0.00% | 0.00% | 0.00% |
| **Manta** | 2k-5kbp | 92.18% | 75.17% | 82.81% | **95.09%** | 87.65% | **71.48%** | 78.74% |
| **gcSV** | >5kbp | **98.52%** | 64.56% | **78.01%** | **97.74%** | **96.30%** | 63.11% | **76.25%** |
| **Delly** | >5kbp | 59.62% | 76.70% | 67.09% | 96.20% | 57.36% | 73.79% | 64.54% |
| **Lumpy** | >5kbp | 63.86% | **77.18%** | 69.89% | 0.00% | 0.00% | 0.00% | 0.00% |
| **Manta** | >5kbp | 90.85% | 67.48% | 77.44% | 96.40% | 87.58% | **65.05%** | 74.65% |
| | | | | **35X-INS** | | | | |
| **dragen** | 50-200bp | 79.95% | **30.54%** | **44.19%** | 64.70% | 51.72% | **19.76%** | 28.59% |
| **gcSV** | 50-200bp | **92.82%** | 25.11% | 39.53% | **74.36%** | **69.02%** | 18.67% | **29.39%** |
| **dragen** | 200-500bp | 94.54% | 21.06% | 34.44% | 69.85% | 66.03% | 14.71% | 24.06% |
| **gcSV** | 200-500bp | **96.24%** | **28.53%** | **44.01%** | **79.66%** | **76.66%** | **22.72%** | **35.05%** |
| **dragen** | 500-1kbp | 90.91% | 7.40% | 13.69% | 56.67% | 51.52% | 4.20% | 7.76% |
| **gcSV** | 500-1kbp | **93.29%** | **7.55%** | **13.97%** | **61.44%** | **57.32%** | **4.64%** | **8.58%** |
| **dragen** | 1k-2kbp | 61.54% | 0.70% | 1.38% | **87.50%** | 53.85% | 0.61% | 1.20% |
| **gcSV** | 1k-2kbp | **85.71%** | **6.27%** | **11.68%** | 63.89% | **54.76%** | **4.00%** | **7.46%** |
| **dragen** | 2k-5kbp | 40.00% | 0.25% | 0.50% | **100.00%** | 40.00% | 0.25% | 0.50% |
| **gcSV** | 2k-5kbp | **82.83%** | **10.38%** | **18.45%** | 63.41% | **52.53%** | **6.58%** | **11.70%** |
| **dragen** | >5kbp | 50.00% | 0.86% | 1.69% | **100.00%** | 50.00% | 0.86% | 1.69% |
| **gcSV** | >5kbp | **56.86%** | **8.33%** | **14.54%** | 62.07% | **35.29%** | **5.17%** | **9.02%** |
| | | | | **35X-DEL** | | | | |

| | | | | | | | | |
|---|---|---|---|---|---|---|---|---|
| **dragen** | 50-200bp | 84.96% | 36.67% | 51.23% | 82.48% | 70.07% | 30.25% | 42.25% |
| **gcSV** | 50-200bp | **88.10%** | **39.04%** | **54.11%** | **86.98%** | **76.63%** | **33.96%** | **47.06%** |
| **dragen** | 200-500bp | **93.49%** | **54.29%** | **68.69%** | **96.28%** | **90.01%** | **52.27%** | **66.13%** |
| **gcSV** | 200-500bp | 87.65% | 52.60% | 65.75% | 95.44% | 83.66% | 50.21% | 62.75% |
| **dragen** | 500-1kbp | 88.35% | **38.06%** | **53.20%** | **93.18%** | 82.33% | **35.47%** | **49.58%** |
| **gcSV** | 500-1kbp | **91.63%** | 34.08% | 49.68% | 93.40% | **85.58%** | 31.83% | 46.41% |
| **dragen** | 1k-2kbp | 92.44% | **59.46%** | **72.37%** | 96.82% | 89.50% | **57.57%** | **70.07%** |
| **gcSV** | 1k-2kbp | **96.91%** | 50.81% | 66.67% | **97.87%** | **94.85%** | 49.73% | 65.25% |
| **dragen** | 2k-5kbp | 96.97% | **73.68%** | **83.74%** | 94.20% | 91.34% | **69.41%** | **78.88%** |
| **gcSV** | 2k-5kbp | **98.97%** | 63.16% | 77.11% | **94.27%** | **93.30%** | 59.54% | 72.69% |
| **dragen** | >5kbp | **97.97%** | **71.08%** | **82.39%** | 96.55% | 94.59% | **68.63%** | **79.55%** |
| **gcSV** | >5kbp | 97.39% | 54.90% | 70.22% | **99.11%** | **96.52%** | 54.41% | 69.59% |

(a) Ground truth callset: NIST HG002 Q100 benchmark (V1.0)

(b) The precisions, sensitivities and F1-scores of various approaches are assessed by Truvari with the following parameter setting "truvari bench --passonly -p 0 --dup-to-in"

(c) All the tools were running in the default parameters.

**Supplementary Table 11. OMIM, dbVar, ClinVar annotated pathogenic SVs discovered by gcSV in 1KGP dataset**

| CHR | POS | AFR-AC | AMR-AC | EAS-AC | EUR-AC | SAS-AC | SV-type | SV-length | Database | Related gene | Related diseases |
|---|---|---|---|---|---|---|---|---|---|---|---|
| chr3 | 101,264,508 | 0 | 0 | 0 | 1 | 0 | Deletion | -9394 | dbVar: nssv15120116 | IMPG2 | Retinal dystrophy |
| chr7 | 4,769,694 | 1 | 0 | 0 | 0 | 0 | Deletion | -42046 | dbVar: nssv15143360 | AP5Z1;FOXK1;MIR4656;RADIL | Spastic paraplegia; autosomal recessive |
| chr21 | 42,477,225 | 32 | 12 | 24 | 11 | 9 | Insertion | 50 | CLN: 469626 | RSPH1 | Primary ciliary dyskinesia |
| chrX | 107,948,288 | 0 | 0 | 0 | 1 | 0 | Deletion | -42623 | morbid: IRS4 | TEX13B | Hypothyroidism; congenital, nongoitrous |

**Supplementary Table 12. All the resources used in this study**

| Tools | Version | URL | Category |
|---|---|---|---|
| **BWA-MEM** | 0.7.15-r1140 | https://github.com/lh3/bwa | SRS aligner |
| **pbmm2** | 0.10.0 | https://github.com/PacificBiosciences/pbmm2 | LRS aligner |
| **Manta** | 1.6.0 | https://github.com/Illumina/manta | SRS SV caller |
| **Delly** | 1.1.6 | https://github.com/dellytools/delly | SRS SV caller |
| **Lumpy** | 0.2.13 | https://github.com/arq5x/lumpy-sv | SRS SV caller |
| **GATK-SV** | - | https://github.com/broadinstitute/gatk-sv | SRS SV caller |
| **DRAGEN** [a] | - | https://github.com/srbehera/DRAGEN_Analysis | SRS SV caller |
| **Blend-Seq** [b] | - | https://github.com/broadinstitute/blend_seq_paper | Hybrid SV caller |
| **Sniffles** | 2.2 | https://github.com/fritzsedlazeck/Sniffles | LRS SV caller |
| **SVDSS** | 2.0.0 | https://github.com/Parsoa/SVDSS | LRS SV caller |
| **cuteSV** | 2.1.1 | https://github.com/tjiangHIT/cuteSV | LRS SV caller |
| **Truvari** | 4.2.2 | https://github.com/ACEnglish/truvari | Benchmarking tool |
| **annotSV** | 3.0.9 | https://github.com/lgmgeo/AnnotSV | Annotation tool |
| **samtools** | 1.10 | https://github.com/samtools/samtools | SAM processing tool |
| **bedtools** | 2.27.1 | https://github.com/arq5x/bedtools2 | BED processing tool |
| **bcftools** | 1.10.2 | https://github.com/samtools/bcftools | VCF sorting tool |
| **bgzip** | 1.17 | https://github.com/samtools/htslib | VCF compression tool |
| **tabix** | 1.17 | https://github.com/samtools/htslib | VCF indexing tool |

(a) The VCF data for Dragen was downloaded from the following address: https://zenodo.org/records/10428664/files/HG002_DRAGEN_SV_hg19.vcf.gz .
(b) The GitHub page for Blend-Seq is temporarily inaccessible; the data of Blend-Seq and GATK-SV were obtained from the following paper: https://www.biorxiv.org/content/10.1101/2024.11.01.621515v1?versioned=true.

### Supplementary Notes

#### 1. The commands and parameters used for read alignment

**BWA-MEM**

```
bwa mem -t 16 -K 100000000 -Y ref.fa read1.fq read2.fq
```

**pbmm2**

```
pbmm2 align --alignment-threads 14 --sort-threads 2 --sort-memory 16G --preset CCS --sort
```

#### 2. Down sample with samtools.

**samtools**

```
samtools view -s 0.5 -b -o input.bam output.bam
```

#### 3. The commands and parameters used for LRS SV calling

**gcSV**

```
gcSV call -c input.bam -r ref.fa -o output.vcf 2> /dev/null
cat output.vcf | bcftools sort | bgzip -cf > output.vcf.gz && tabix output.vcf.gz
```

**sniffle2 max sensitive mode**

```
sniffles --input input.bam --vcf output.vcf --minsupport 1 --qc-output-all --qc-coverage 1 --long-dup-coverage 1 --detect-large-ins True --reference ref.fa
cat output.vcf | bcftools sort | bgzip -cf > output.vcf.gz && tabix output.vcf.gz
```

**sniffle2 default mode**

```
sniffles --input input.bam --vcf output.vcf --reference ref.fa
cat output.vcf | bcftools sort | bgzip -cf > output.vcf.gz && tabix output.vcf.gz
```

**SVDSS**

```
SVDSS_linux_x86-64 index --reference ref.fa --index ref.fmd
SVDSS_linux_x86-64 smooth --reference ref.fa --bam input.bam > SMO.bam && samtools index SMO.bam
SVDSS_linux_x86-64 search --index ref.fmd --bam SMO.bam > SPE.txt
SVDSS_linux_x86-64 call --reference ref.fa --bam SMO.bam --sfs SPE.txt > output.vcf
cat output.vcf | bcftools sort | bgzip -cf > output.vcf.gz && tabix output.vcf.gz
```

**cuteSV2 max sensitive mode**

```
cuteSV --genotype -s 1 input.bam ref.fa output.vcf ./work_dir
cat output.vcf | bcftools sort | bgzip -cf > output.vcf.gz && tabix output.vcf.gz
```

**cuteSV2 default mode**

```
cuteSV --genotype input.bam ref.fa output.vcf ./work_dir
cat output.vcf | bcftools sort | bgzip -cf > output.vcf.gz && tabix output.vcf.gz
```

#### 4. The commands and parameters used for SRS SV calling

**gcSV**

```
gcSV ngs_fa_stat ref.fa > ref.stat.txt
gcSV ngs_trans_reads ref.fa input.bam TL.bam
samtools sort --output-fmt=BAM -o TL.sort.bam TL.bam
gcSV call -n input.bam -L TL.sort.bam -r ref.fa -I ref.stat.txt -o output.vcf 2> /dev/null
cat output.vcf | grep -v -E "SVTYPE=BND” | bcftools sort | bgzip -cf > output.vcf.gz && tabix output.vcf.gz
```

**manta**

```
configManta.py --bam input.bam --referenceFasta ref.fa --runDir ./manta_work
./manta_work/runWorkflow.py -m local -j 16
zcat ./manta_work/results/variants/diploidSV.vcf.gz | grep -v -E "SVTYPE=BND” | bcftools sort | bgzip -cf > output.vcf.gz && tabix output.vcf.gz
```

**delly**

```
delly call -g ref.fa input.bam > output.vcf
cat output.vcf | grep -v -E "SVTYPE=BND” | bcftools sort | bgzip -cf > output.vcf.gz && tabix output.vcf.gz
```

**lumpy**

```
samtools view -F 1294 input.bam > DIS.bam
samtools sort -@ 16 DIS.bam -o DIS.sort.bam && samtools index DIS.sort.bam
samtools view -h input.bam | ./lumpy-sv/scripts/extractSplitReads_BwaMem -i stdin | samtools view -Sb - > SIP.bam
samtools sort -@ 16 SIP.bam -o SIP.sort.bam && samtools index SIP.sort.bam
lumpyexpress -B input.bam -S SIP.sort.bam -D DIS.sort.bam -o output.vcf
cat output.vcf | grep -v -E "SVTYPE=BND|SVTYPE=TRA|SVTYPE=INV"  | bcftools sort | bgzip -cf > output.vcf.gz && tabix output.vcf.gz
```

### 5. The commands and parameters used for Hybrid SV calling

**gcSV**

```
gcSV ngs_fa_stat ref.fa > ref.stat.txt
gcSV ngs_trans_reads ref.fa NGS.input.bam TL.bam
samtools sort --output-fmt=BAM -o TL.sort.bam TL.bam
gcSV call -c TGS.input.bam-n NGS.input.bam -L TL.sort.bam -r ref.fa -I ref.stat.txt -o output.vcf 2> /dev/null
cat output.vcf | grep -v -E "SVTYPE=BND” | bcftools sort | bgzip -cf > output.vcf.gz && tabix output.vcf.gz
```

### 6. The commands and parameters used for 1KGP re-analysis

**SURVIVOR (SV merging)**

```
SURVIVOR merge vcf_list.txt 500 1 1 1 0 30 output.vcf
```

**AnnotSV**

```
AnnotSV -SVinputFile inputFile -outputDir output -outputFile ./output/{sample_name}.csv -genomeBuild GRCh38 -SVminSize 30
```

**gcSV VNTR analysis**

```
gcSV tools vntr_analysis ref.fa single_sample_vcf_list.txt VNTR_region.bed 1kgp_sample_info.txt
```

### 7. The commands and parameters used for benchmarking

**truvari bench**

```
truvari bench --passonly -p 0 -P 0.7/0.9/0.95/0.99/1--dup-to-in -c cmp.vcf.gz -b base.vcf.gz -o ./result.dir --reference ref.fa --includebed region.bed
```

**truvari refine**

```
truvari bench --passonly --pick ac --dup-to-ins --reference ref.fa --includebed region.bed -c cmp.vcf.gz -b base.vcf.gz -o ./result.dir

truvari refine --recount --use-region-coords --use-original-vcfs --align mafft --reference ref.fa

--regions ./result.dir/candidate.refine.bed ./output_dir

truvari ga4gh --input ./output_dir --output ./output_dir/combined_result --with-refine
```
